# Generative inverse design steers human hematopoietic cell fate *in vitro*

**DOI:** 10.64898/2026.10.07.756550

**Authors:** Licyel Paulas-Condori, Alessandro Palma, Lorenzo Consoli, Juan Antonio Rubio-Lara, Olivia Hidajat, Weizhong Zheng, Dominik Klein, Deborah G. Martinuzzi, Zhimeng Wu, Andrea Dittadi, Nicola K. Wilson, Berthold Göttgens, Fabian J. Theis

## Abstract

Inverse design has advanced the discovery of molecules and materials. Yet it remains largely unexplored for steering cell fate, where inputs are continuous, combinatorial culture conditions and outputs are distributions of cell states. Human hematopoiesis, in which combinations and doses of growth factors steer stem cells toward each blood lineage, is a demanding test case. While forward models can evaluate discrete conditions, the vast combinatorial space renders searching for a specific, target-producing design an intractable inverse problem. Here, we introduce LabCompass, an inverse-modeling framework that optimizes experimental designs toward candidates expected to produce specified target cell states, using a flow-matching generative model guided by cellular phenotypes predicted by a forward surrogate. We applied LabCompass to primary human CD34+ *in vitro* cultures, where continuous cytokine and small-molecule parameters govern hematopoietic cell fate. In a lab-in-the-loop setting, we experimentally validated model-driven designs and used these measurements to update the model across four successive loops. LabCompass recovered established lineage-inducing designs, improved predictions with new data, and enriched target populations across lineages. Incorporating new parameters expanded the design space and enriched rare populations beyond initial reach. Across loops, we generated a 78-million-cell spectral flow cytometry dataset and identified distinct culture conditions converging on the same target cell states. Our results establish generative inverse modeling as a compelling paradigm for navigating high-dimensional, continuous experimental spaces, and demonstrate that virtual cell models should be judged not only by how accurately they predict cellular responses, but also by how effectively they guide experiments toward desired biological outcomes.

## Introduction

Designing biological experiments is complex because the space of possible conditions is often continuous and combinatorial, making exhaustive testing impractical. Outcomes often depend on combinations and doses of signals, creating a vast search space for experimental design. Predictive models of cellular perturbation response, known as virtual cell models^1^, offer a computational alternative. These models have captured single-cell transcriptomic responses to genetic^2^, chemical^3–6^, cytokine perturbations^7^, and cellular responses at scale^8–10^. However, most methods are limited to forward modeling: predicting the outcome of a given set of inputs rather than identifying which inputs will produce a desired phenotype^11^. This limitation becomes increasingly important as design spaces grow, particularly when dosages and experimental conditions are continuous, yet exhaustive forward search requires discretization^12^.

Inverse modeling offers a natural alternative: instead of forward prediction, inputs are optimized to achieve a desired output, with experimentally tested designs feeding back into subsequent rounds in a lab-in-the-loop workflow. Related approaches have reduced experimental burden in property-driven molecular design^13–15^ and Bayesian search of culture media^16–18^. In cellular phenotypic optimization, however, existing approaches have largely relied on retrieval or discrete search^19,20^, ranking predefined candidate designs, such as gene perturbations^21^, by their ability to induce a target state^22^. Such fixed catalogues cannot generate novel experimental designs and scale poorly to continuous, high-dimensional phenotypic optimization tasks. This limitation reflects a broader bottleneck in biology: phenotypic outputs are often governed by vast, non-linear combinatorial spaces that rapidly exceed what exhaustive experimental, and indeed computational, testing can cover.

Generating defined populations from hematopoietic stem/progenitor cells (HSPCs) is important for transfusion medicine, cell and gene therapy, disease modeling, and drug screening, using approaches that include HSC (hematopoietic stem cell) expansion, directed differentiation, and 3D hematopoietic and bone marrow organoids^23–25^. What blood cells are produced in any of those settings depends on combinations and doses of signals rather than single factors, yet developing new culture conditions largely relies on an empirical trial-and-error approach, complicated by the notion that the same factor can favor expansion or differentiation depending on concentration and timing, and that each condition yields a distribution of co-existing states rather than a single cell type. Protocol development is consequently slow and material-intensive, with the current gold-standard protocol for cord blood-derived megakaryocytes distilled painstakingly and optimized from 1,500 umbilical cord blood samples^26^. For most hematopoietic fates, it also remains unknown whether a target state is reachable through one singular combination of signals or several distinct routes.

Here we introduce LabCompass, a generative inverse-modeling framework for high-dimensional experimental design. Rather than retrieving candidates from a fixed catalogue, LabCompass learns from experimentally observed designs and uses this knowledge to generate novel designs that remain realistic and experimentally feasible. Based on flow matching^27–29^, it combines a generative model with a forward surrogate that predicts cellular phenotypes. During generation, surrogate gradients steer sampling toward designs predicted to achieve a specified target state^30^, while a generative prior constrains sampling to plausible regions. Because optimization occurs at inference time, the same model can be directed toward new phenotypic objectives without retraining.

We show that LabCompass can generate experimental designs predicted to enrich specific cell types, including megakaryocyte, early lymphoid, monocytic, and erythroid populations. Generating a 78-million-cell dataset, we identified distinct culture conditions that produced similar cell-type distributions, revealing multiple routes to the same cellular fate. Inverse design has previously reshaped how molecules and materials are discovered. LabCompass shows that the same logic applies to the conditions under which cells are grown.

### LabCompass enables experimental design optimization through generative inverse modeling

We formulate cellular fate prediction as a perturbation modeling problem, where perturbations correspond to experimental designs and a forward model maps each design to its resulting cellular response. We adopt a generative formulation, combining a generative model of cellular states, conditioned on the experimental design *x* ϵ *X*, with a phenotype prediction function, for example, a classifier mapping cells to their type, that reads out the response *y* ϵ *Y* of interest, in this case the resulting cell-type compositions.

The forward model consists of a perturbation model and a phenotypic readout function *h*. *F* maps designs *x* ϵ *X* to distributions over *ε*, generating synthetic cellular profiles *ê*∼*p*(· ∣*x*); *h* then extracts a cellular property of interest, *ŷ* = *h* (*ê*), which can range from discrete descriptors, such as cell-type frequencies, to continuous ones, such as a target cell surface marker expression value, or cell states. This formulation flexibly accommodates any representation of cells, while adapting to versatile definitions of the system’s response (**Fig. 1A**). By predicting the experimental response across a large set of unexplored designs *in silico*, forward models can identify designs inducing a desired response. Designs are ranked by how their predicted response differs from the desired one and selected for experimental validation (**Supplementary Fig. 1A**). However, exhaustive enumeration and random search quickly become intractable in large combinatorial or continuous design spaces.

**Figure 1.**
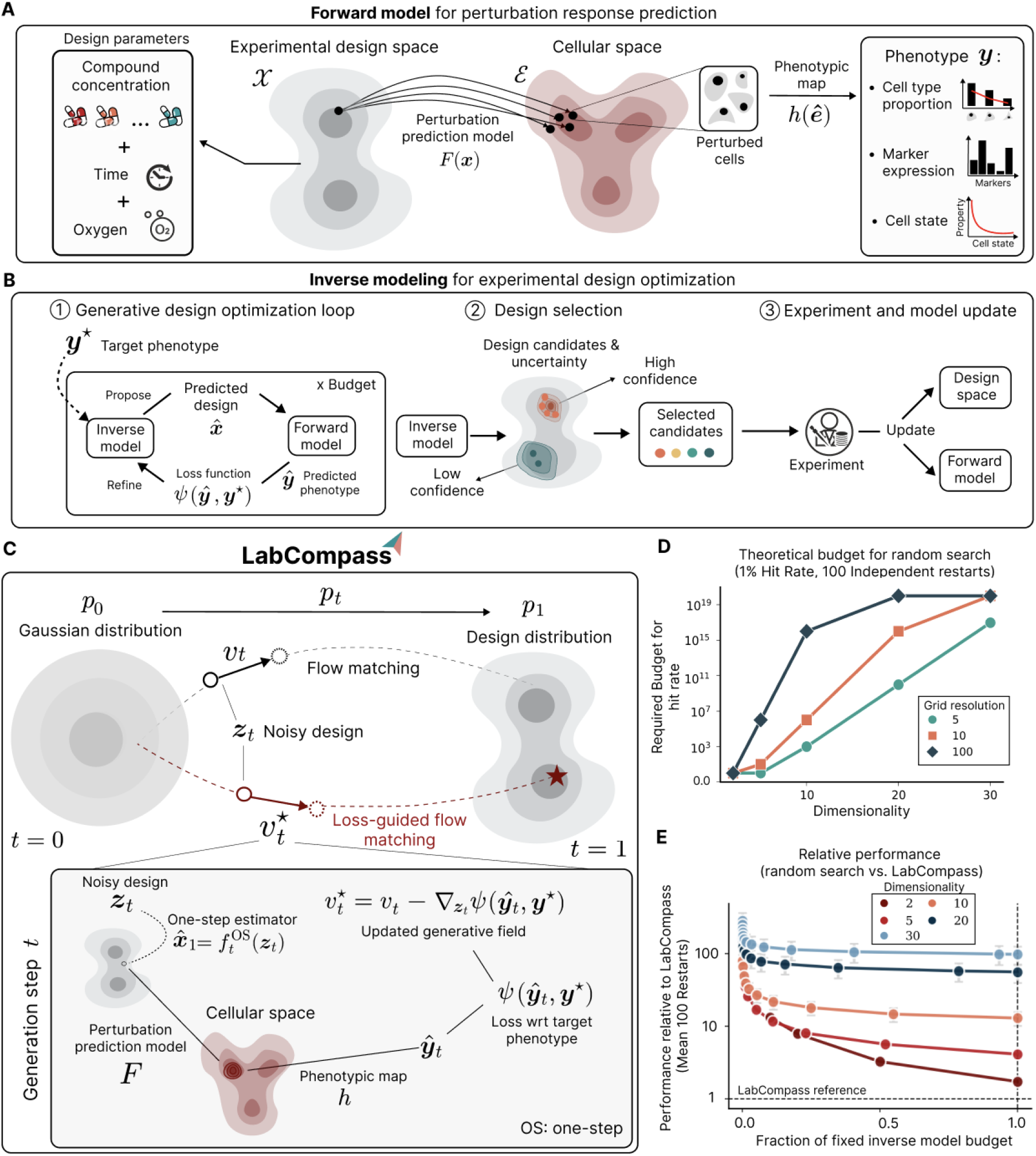
LabCompass couples forward surrogate modeling with inverse generative sampling to enable target-guided design generation. **A:** Forward model for perturbation response prediction. Experimental designs are mapped to conditional distributions over the cellular state space via a surrogate forward model. The phenotypic readout is then evaluated on the cellular states. **B:** Inverse modeling for experimental design optimization. (1) Given a target phenotype, the inverse model iteratively refines candidate designs based on the predicted phenotypic response, within a fixed budget. (2) Designs are ranked by predicted performance and uncertainty to balance exploration and exploitation, and are then selected for wet-lab evaluation. (3) New measurements are used to update both the design space (prior model) and the forward model. **C:** A single generative inverse step. The prior model interpolates between a standard Gaussian *t =* 0 at and the data distribution at *t =* 1, using the prior velocity field. LabCompass steers generation toward the target phenotype by mapping intermediate noisy designs to the estimated clean designs via a one-step estimator. The estimated data point is used to evaluate the forward model, compute the phenotypic loss, and backpropagate it. The resulting field yields samples minimizing the phenotypic loss function. **D:** Required number of function evaluations (y-axis) versus problem dimensionality (x-axis) to reach a 1% hit rate with 100 restarts for random search across three grid resolutions (5, 10, 100). The required budget increases polynomially with the resolution. **E:** Relative performance of random search versus LabCompass on the synthetic grid-optimization problem with resolution of 5, across dimensionalities (2–30) and relative budgets. LabCompass ran as a fixed-budget reference (surrogate evaluations; dashed line); random search’s budget increased until it matched this reference.

Conversely, inverse models use *F*, now treated as a differentiable surrogate for experimental outcomes, to identify regions of the design space likely to induce a target phenotype *y*^⋆^ ϵ y, reducing the number of uninformative virtual screens (**Supplementary Fig. 1B**). The inverse model produces a pool of candidates ranked by the forward surrogate’s predictive uncertainty, enabling both exploratory and exploitative design strategies (see **Supplementary Fig. 1C** and **Methods**): exploration prioritizes uncertain regions, while exploitation favors areas predicted to yield favorable outcomes with high confidence. Selected candidates are validated experimentally, with results looped back to update the design space and the forward surrogate (**Fig. 1B**).

LabCompass is our flow-matching-based inverse modeling framework for experimental design in single-cell biology. Rather than generating designs independent of the target phenotype and evaluating them only a posteriori, LabCompass lets the target state actively steer generation from the first step. Candidate designs arise from a flow *p*_*t*_ that gradually reshapes random noise into the learned distribution of experimental designs, guided by a velocity field *υ*_*t*_ and acting as a prior over plausible designs. At each step towards design generation, the forward surrogate predicts the cellular response induced by the current candidate and compares it to *y*^⋆^ via a loss *ψ*. This comparison and the gradient of the loss are used to adjust the generative trajectory, nudging it toward designs expected to yield the target phenotype while remaining anchored to realistic designs (**Fig. 1C**). Each generated design is therefore refined over many guided steps rather than produced in one shot. This optimization happens entirely at inference time, requiring no retraining of either model, and enables efficient amortized design optimization across new targets or objectives, avoiding the need to brute-force evaluate options across what is often a combinatorially large experimental space. LabCompass is not tied to any specific forward surrogate: any model differentiable with respect to *x* can be used, letting the framework incorporate future advances in forward modeling as they emerge (see “Inverse design of viral expression states in human cytomegalovirus (HCMV) infection” in the **Methods** and **Supplementary Fig. 2** for a continuous gene-expression-based optimization applied to single-cell transcriptomic data).

Because even a simple random search can be a surprisingly competitive baseline in black-box optimization, we asked whether LabCompass provides an advantage beyond uninformed exploration, particularly as the design space becomes more complex. We, therefore, designed a toy problem for a rigorous and controlled comparison with uniform random search: the design space was given by a *D*-dimensional hypercube, discretized at resolution *R* ; the objective was to sample a randomly selected point from the grid, chosen to be the optimization target. A theoretical analysis of this problem illustrates the unfavorable scaling of random search – the probability of sampling the target decreases rapidly with increasing dimensionality and resolution (**Fig. 1D, Supplementary Fig. 1D**; **Methods**). The experimental comparison between LabCompass and uniform random search aligned with the theoretical considerations: LabCompass outperformed random search across all tested dimensionalities and resolutions, with the performance gap broadly widening as dimensionality increased. For a resolution of 5, the relative performance increased monotonically from 1.73 in 2 dimensions to 97.91 in 30, providing empirical support for the theory (**Fig. 1E, Supplementary Fig. 1E**). These results highlighted the benefit of using the learned design prior together with feedback from the forward surrogate to focus exploration where favorable outcomes are more likely.

### A combinatorial screen explores the design space of human *in vitro* hematopoiesis

To train LabCompass in human HSPCs, we first established an experimental system that captures diverse hematopoietic states. Because current *in vitro* culture systems^24,25,31^ typically do not balance HSC self-renewal and multilineage differentiation (**Fig. 2A**), we screened 116 combinations of known hematopoietic-supportive molecules with varying oxygen concentrations, generating a dataset spanning a broad range of phenotypic outcomes. Efficiently navigating the hematopoietic design space requires a high-dimensional, scalable readout. While scRNA-seq provides cellular resolution of multilineage differentiation, spectral flow cytometry offers multi-dimensional resolution at higher throughput and lower cost. Even then, varying 10 parameters across 4 concentrations under two oxygen levels spans millions of possible designs.

**Figure 2.**
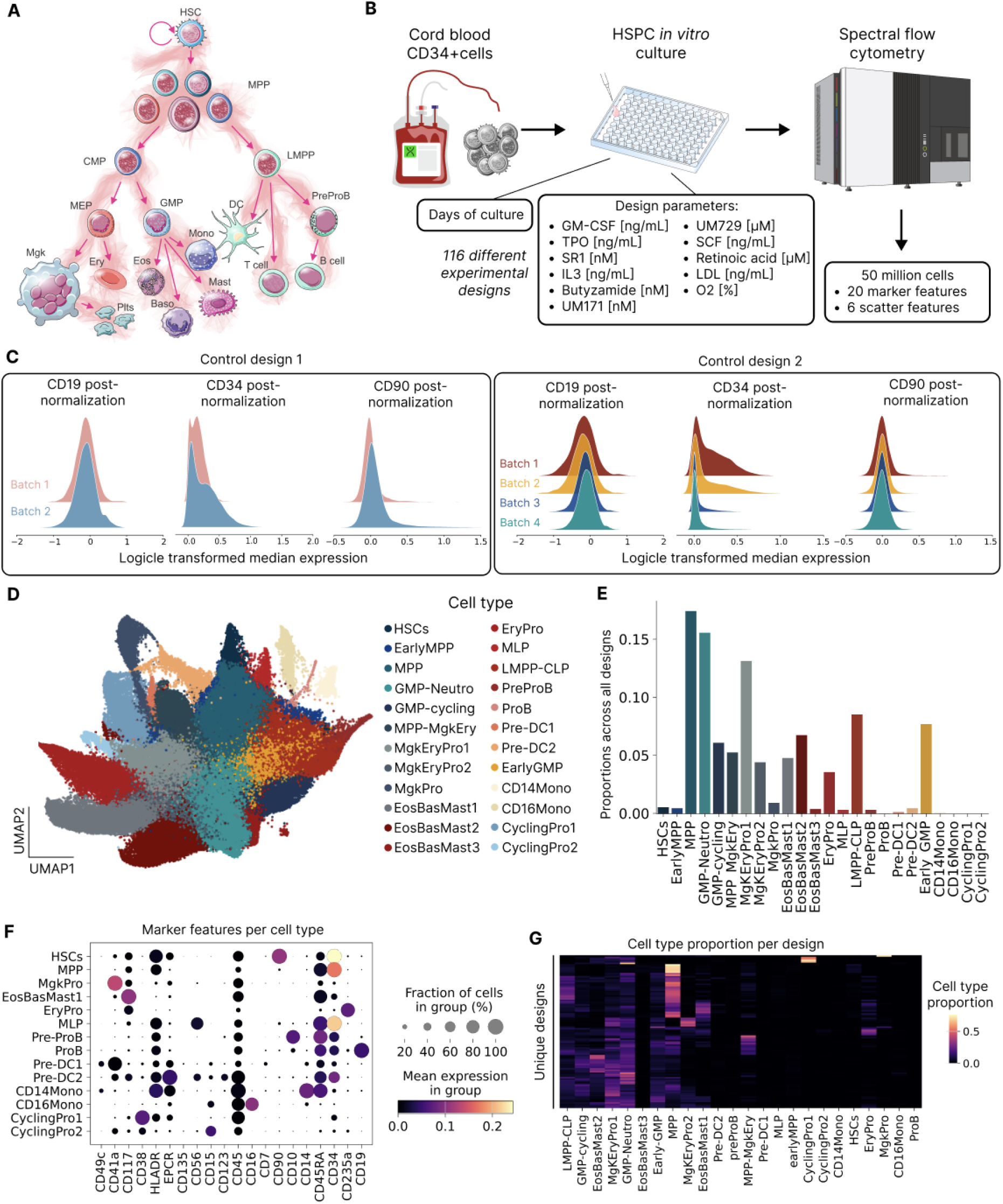
A high-dimensional combinatorial screen maps the design space of human *in vitro* hematopoiesis. **A:** Schematic of the hematopoietic landscape, depicting the differentiation hierarchy from hematopoietic stem cells (HSCs) through progenitor states to terminal lineages. MPP = Multipotent Progenitors, CMP = Common Myeloid Progenitors, LMPP = Lymphoid-Myeloid Primed Progenitors, MEP = Megakaryocyte-Erythroid Progenitors, GMP = Granulocyte-Monocyte Progenitors, Mgk = Megakaryocyte, Plts = Platelets, Ery = Erythrocyte, Eos = Eosinophil, Baso = Basophil, Mono = Monocyte, Mast = Mast Cell, DC = Dendritic cell. **B:** Experimental workflow: umbilical cord blood CD34^+^ cells are cultured in different HSPC *in vitro* culture conditions or designs (116 designs), which included 11 design parameters and days of culture, followed by spectral flow cytometry profiling of 50 million cells across 20 marker features and 6 scatter features. **C:** Logicle-transformed median expression of CD19, CD34, and CD90 across repeated measurements of two control designs (Control design 1, two batches (Design 240 & 280); Control design 2, four batches (Design 158, 205, 258 & 277)), demonstrating instrument stability and inter-batch reproducibility. **D:** UMAP visualization of the combined hematopoietic landscape from all 116 experimental designs, colored by cell type as identified by the spectral flow cytometry panel. EarlyMPP = Early Multipotent Progenitors, GMP-Neutro = GMP-neutrophil, MPP-MgkEry = MPP-Megakaryocyte-Erythroid, MgkEryPro1 = Megakaryocyte-Erythroid Progenitor 1, MgkEryPro2 = Megakaryocyte-Erythroid Progenitor 2, MgkPro = Megakaryocyte Progenitor, EosBasMast1 = Eosinophil-Basophil-Mast cell Progenitor 1, EosBasMast2 = Eosinophil-Basophil-Mast cell Progenitor 2, EosBasMast3 = Eosinophil-Basophil-Mast cell Progenitor 3, EryPro = Erythroid Progenitor, MLP = Multi-Lymphoid Progenitor, LMPP-CLP = Lymphoid-Primed Multipotent Progenitor-Common-Lymphoid-Progenitor, Pre-DC1 = Pre-Dendritic Cell 1, Pre-DC2 = Pre-Dendritic Cell 2, EarlyGMP = Early Granulocyte-Monocyte-Progenitor, CD14Mono = CD14+ monocyte, CD16Mono = CD16+ monocyte, CyclingPro1 = Cycling Progenitor 1, CyclingPro2 = Cycling Progenitor 2. **E:** Measured proportion of cells belonging to each hematopoietic lineage across all designs. **F:** Dot plot of marker feature expression per cell type. Dot size indicates the fraction of cells expressing each marker within a group; color indicates mean expression level. **G:** Heatmap of cell-type proportions across unique experimental designs in the dataset.

Primary CD34+ HSPCs, isolated from human umbilical cord blood, were cultured in basal media supplemented with diverse combinations and concentrations of 10 cytokines/small molecules as well as hypoxia (10%) vs. normoxia (20%), selected for their reported effects on distinct stages of hematopoiesis^25,32–41^ (**Fig. 2B**). Cellular output of each design was characterized using a standardized 20-color spectral flow cytometry panel that incorporated established hematopoietic markers^42–44^, profiling 50 million total cells across the screening (**Fig. 2B**).

Applying a standardized preprocessing pipeline (see **Methods**) mapped the spectral flow cytometry data into a continuous, shared multidimensional feature space. This allowed cells across all 116 conditions to be co-embedded and analyzed using standard single-cell tools, such as UMAP^45^ and unbiased clustering. To exclude batch effects and ensure consistency, control designs were periodically profiled using the spectral flow cytometer’s standardization mode (see **Methods**; **Fig. 2C**). Across the screen, we observed a broad spectrum of immunophenotypically defined hematopoietic populations, ranging from phenotypic HSCs (pHSC), which includes the population of CD34+CD45RA-CD90+EPCR+ cells, to more differentiated cell states, such as megakaryocyte progenitors (MgkPro), erythroid progenitors (EryPro), and monocytes (CD14 Mono and CD16 Mono) (**Fig. 2D**; **Supplementary Fig. 3**). The relative abundance of these populations varied across designs, with multipotent progenitors (MPPs), GMP-neutrophil (GMP-Neutro), and megakaryocyte-erythroid progenitor 1 (MgkEryPro1) among the most enriched populations, whereas monocytes, cycling progenitors, and proB populations were less abundant potentially due to the lack of cytokines to support these (**Fig. 2E, F**). Individual culture conditions selectively enriched distinct cell populations to varying degrees, highlighting that hematopoietic output is strongly shaped by the composition of the design (**Fig. 2G**).

Overall, the screen encompassed 116 designs with 10 biochemical design parameters at varying concentrations, plus oxygen level and days of culture, generating a high-dimensional immunophenotypic dataset. Each design is represented as a single multivariate vector of continuous concentrations, used by LabCompass to generate cellular fates for inverse modeling. Because these parameters are continuous, the resulting design space cannot be exhaustively enumerated, making it infeasible to fully explore this combinatorial parameter space. We, therefore, investigated whether LabCompass could learn predictive mappings from this screening dataset and use it to prioritize designs for enriching desired hematopoietic cell states.

### LabCompass recovers known hematopoietic lineages and learns from experimental feedback

To establish the feasibility of model-guided exploration, we designed an experimental loop (Loop 1) with three objectives: to test whether LabCompass could identify conditions known to enrich target cell types, assess whether the forward surrogate could reconstruct cell-type compositions of unseen designs, and determine whether experimental feedback could improve model predictions and uncertainty. We define a *loop* as a complete iteration of model prediction, design selection, experimental measurement, and model refinement. Here, we focused on megakaryocyte progenitors (MgkPro), for which enriching conditions have been reported^26,38,46,47^. Starting from 116 designs and their spectral flow cytometry measurements (**Supplementary Fig. 3, 4A-B**), we trained the forward model and design prior, then queried LabCompass for MgkPro-enriching designs (**Fig. 3A**). Generated designs recapitulated real experimental parameter ranges and induced cell states within the observed hematopoietic landscape, indicating qualitatively sensible inverse behavior before any experimental feedback and uncertainty calibration (**Supplementary Fig. 5A-C**).

**Figure 3.**
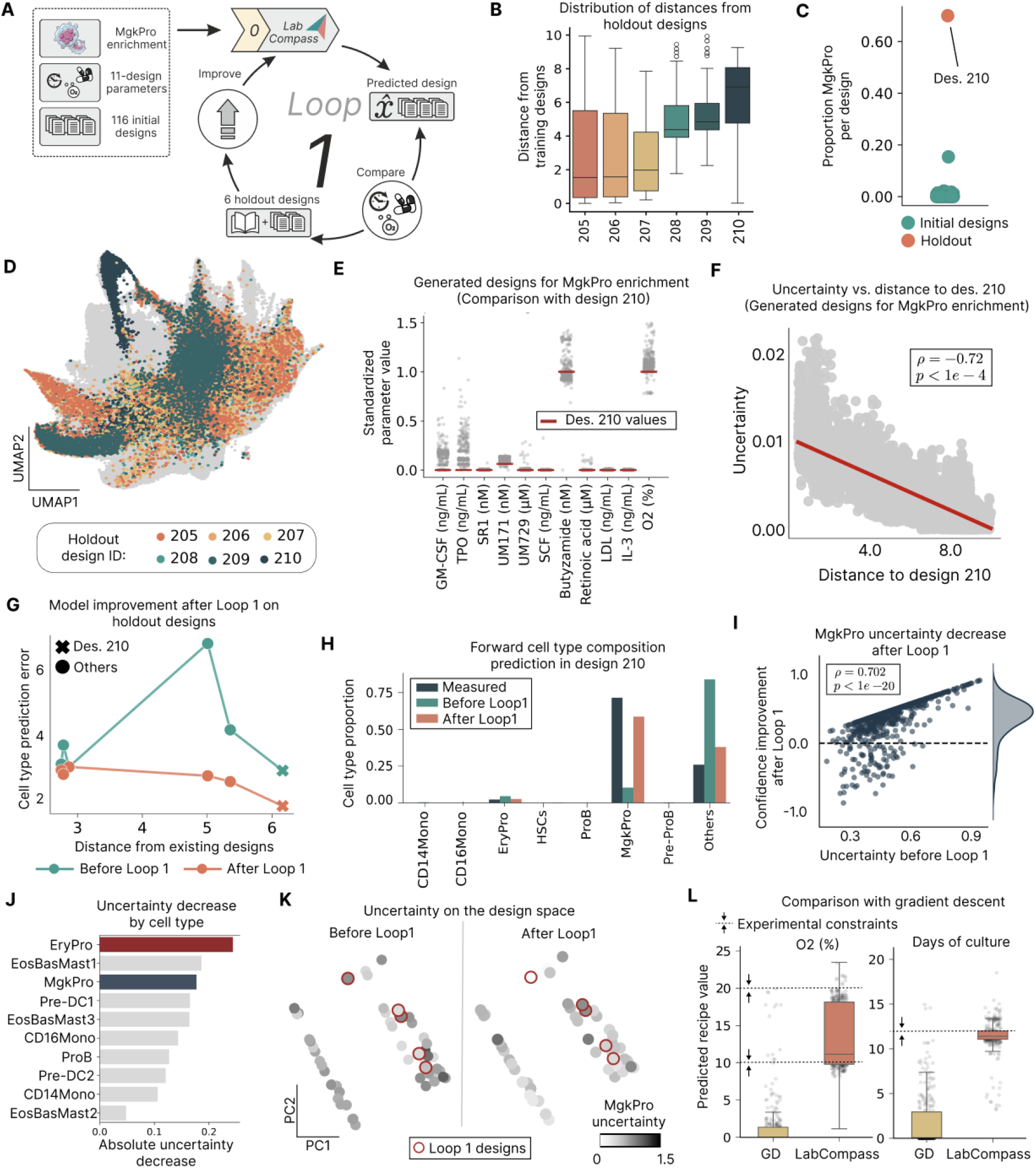
LabCompass enables model-guided exploration through lineage retrieval, forward prediction, and adaptive learning. **A:** Schematic of Loop 1: LabCompass is queried for MgkPro enrichment, using held-out designs as ground truth for retrieval. **B:** Euclidean distance from the training data for each of six held-out conditions (Des. 205–210), showing variable proximity to the training designs. **C:** Distribution of MgkPro proportions measured across all initial designs. Design 210 is marked in orange. **D:** UMAP visualization of the hematopoietic landscape, highlighting cells from each held-out design (Des. 205–210). **E:** Marginal distributions of molecular concentrations from LabCompass-optimized designs for MgkPro enrichment, using the original surrogate model. Experimental parameters of design 210 are marked with a red horizontal bar. **F:** Distance from design 210 in the experimental design space (x-axis) versus the surrogate model’s predictive uncertainty (y-axis). **G:** Forward-prediction error measured as cross-entropy versus average distance to existing designs, before and after observing the held-out designs (Des. 210 marked with ×; others with circles). **H:** Measured versus predicted cell-type composition for design 210, before and after model update. **I:** Predictive uncertainty before Loop 1 (x-axis) versus confidence improvement after Loop 1 (y-axis), for MgkPro-enriching designs generated by LabCompass. **J:** Mean absolute reduction in predictive uncertainty per cell type across experimental designs. **K:** PCA projection of experimental designs, colored by MgkPro-enrichment uncertainty, before and after inclusion of the held-out designs (outlined in red). **L:** Oxygenation level (O2, %) and days of culture for gradient-descent (GD)- and LabCompass-generated samples. Experimental constraints are indicated with dashed lines/arrows.

To evaluate retrieval, we held out six experimental designs (designs 205-210) of varying distances from the training designs (**Fig. 3B**). Design 210, selected from the literature based on reagent availability, was an exceptional megakaryocyte-inducing condition, yielding approximately 70% MgkPro where no initial design in the training set exceeded 16% (**Fig. 3C**), while the other held-out designs spanned diverse hematopoietic differentiation branches (**Fig. 3D**). Recovering design 210 without having observed it therefore constituted a stringent test of retrieval. In some cases, LabCompass-generated designs recapitulated the biochemical composition of design 210, including concentrations close to its defining conditions of UM171 (70 nM) and butyzamide (100 nM) (**Fig. 3E**; **Supplementary Fig. 6A, 6E**). Importantly, predictive uncertainty was negatively correlated with Euclidean distance to design 210 (Spearman ρ = −0.72, P < 0.0001; **Fig. 3F**), indicating that uncertainty concentrated around this experimentally relevant yet previously unseen region, naturally prioritizing it for exploration.

We next assessed whether the surrogate forward model could reconstruct general cell-type compositions for the held-out designs. Before model updating, reconstruction error varied non-monotonically with distance from the training distribution, without a consistent increase across all held-out designs (**Fig. 3G**). We, therefore, incorporated all six held-out measurements into the training set to improve predictions in previously underrepresented regions. The updated model showed reduced prediction error overall (**Fig. 3G**) and improved recovery of experimentally measured cell-type proportions and marker expression across individual cell types on designs 207, 208, and 210 (**Fig. 3H; Supplementary Fig. 6B-C**), demonstrating that experimental feedback can progressively refine the surrogate model and strengthen the optimization signal. Entropy of the predicted cell-type composition scaled with residual error, indicating that designs yielding more heterogeneous populations remain the hardest to predict (**Supplementary Fig. 6D**).

We further quantified how these new experimental measurements affected predictive uncertainty. Evaluating the generated designs under the surrogate forward model before and after data integration revealed substantially reduced uncertainty, with larger reductions occurring for designs that initially had greater uncertainty (Spearman ρ = 0.702, P < 10^−20^; **Fig. 3I**), indicating that experimental feedback was most informative where the model was initially least confident. Across cell types, uncertainty also decreased to varying degrees, with erythrocytes and megakaryocytes among the largest reductions, consistent with their shared erythroid–megakaryocyte branch (**Fig. 3J**). Upon observation of design 210, uncertainty reductions were concentrated in specific regions of the experimental design space (**Fig. 3K**), consistent with new measurements constraining nearby conditions while having limited influence on distant conditions.

Finally, we compared LabCompass with Gradient Descent (GD) to assess the benefit of flow-matching-based generative optimization for MgkPro enrichment. GD directly optimizes the condition-space variables by backpropagating the objective through the forward model, but because the surrogate is reliable primarily near the training distribution, its unconstrained trajectories can leave the experimentally supported design manifold^48^. In contrast, LabCompass steers a generative model over the condition space, constraining optimization to experimentally plausible regions. *In silico* comparison confirmed that LabCompass samples adhered more closely to the design manifold, whereas GD trajectories were prone to degenerate directions favored by the surrogate and to violate experimental constraints (**Fig. 3L; Supplementary Fig. 7A-B**). Strong gradient fields could also collapse the initialization distribution to a single point, eliminating solution diversity through *mode collapse* (**Supplementary Fig. 7C-D**). The generative formulation of LabCompass was therefore verified to better preserve diversity during optimization, unlike deterministic GD.

In summary, Loop 1 shows that LabCompass can retrieve designs known to drive specific lineages, reconstruct cell-type composition of unseen conditions, and effectively learn from new data.

### LabCompass generates cell-type-specific design patterns and improves its own certainty

Having established LabCompass’s ability to perform exploration and learn from new data, we transitioned from *in silico* validation in Loop 1 to the first model-informed wet-lab experimental loop. We designed these experiments to address the exploration–exploitation trade-off by testing whether high-confidence model predictions could reproducibly induce the desired differentiation fate experimentally, while assessing whether inclusion of uncertain but promising candidates could improve both performance and model confidence. This represents a critical step towards autonomous experimentation, in which models iteratively design experiments, learn from their outcomes, and improve subsequent predictions, thereby closing the model–wet-lab feedback loop (**Fig. 4A**).

**Figure 4.**
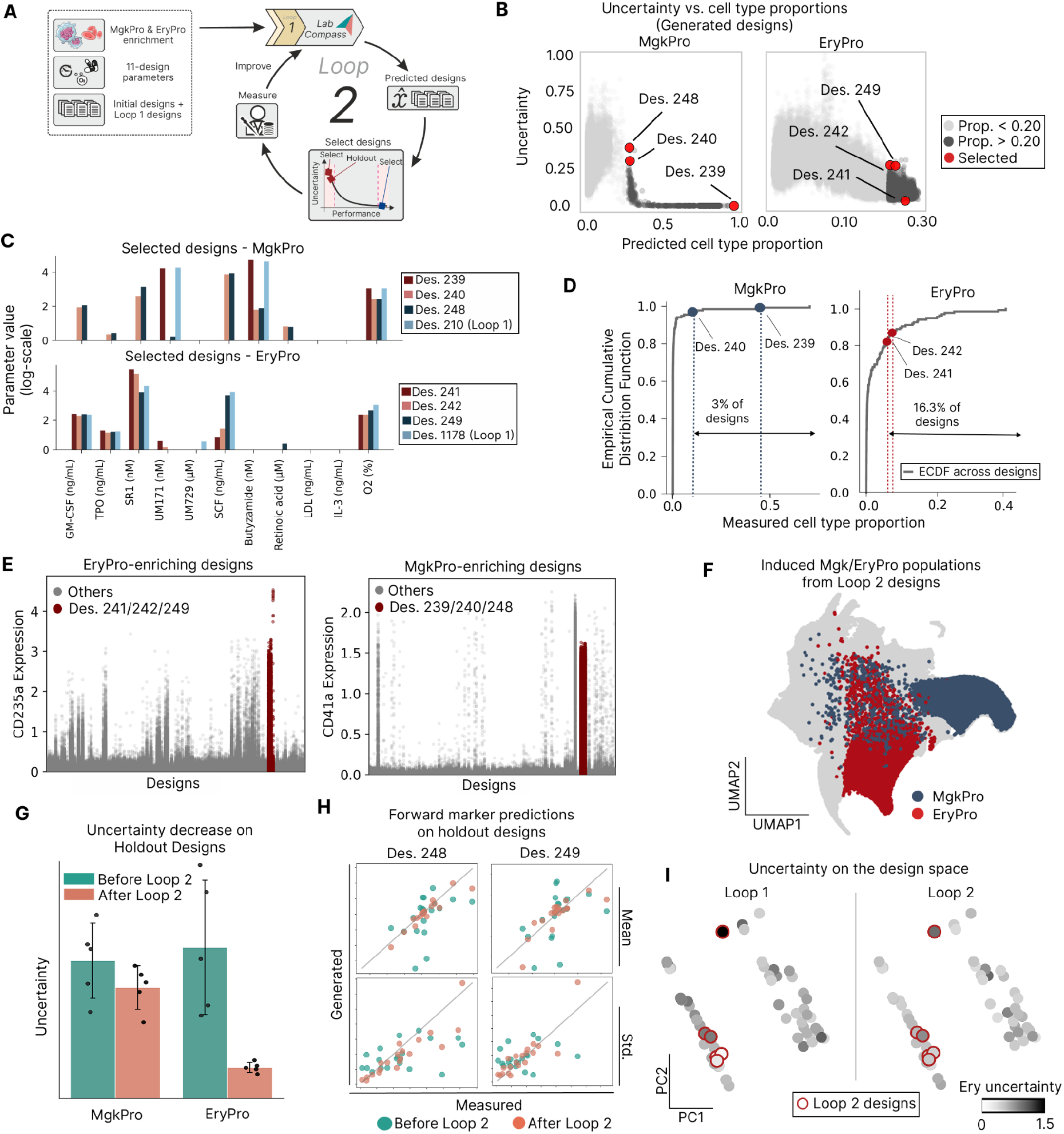
LabCompass reduces predictive uncertainty and improves accuracy by selecting informative designs. **A:** Schematic of Loop 2: LabCompass is queried for target cell-type enrichment. Generated designs are ranked by target proportion and predictive uncertainty: one exploitative and two exploratory design candidates are selected for wet-lab evaluation. **B:** Predicted target cell-type proportion versus predictive uncertainty for MgkPro (left) and EryPro (right) designs. Designs are shaded by predicted proportion (light gray, < 0.20; dark gray, > 0.20); red points mark candidates selected for experimental evaluation (Des. 239, 240, 248 for MgkPro; Des. 241, 242, 249 for EryPro). **C:** Log-transformed value of each design parameter, in the units indicated on the axis, across selected designs for MgkPro (top) and EryPro (bottom) enrichment, including the most enriching design from Loop 1 for each target cell type, for comparison. **D:** Empirical cumulative distribution function (ECDF) of measured target cell-type proportion across evaluated designs for MgkPro (left) and EryPro (right), with selected designs (Des. 239, 240; Des. 241, 242) and the top percentile they fall within indicated. **E:** Measured marker expression (CD235a for EryPro-enriching designs, left; CD41a for MgkPro-enriching designs, right) across all designs (gray), with generated designs highlighted (red; Des. 241/242/249 and Des. 239/240/248, respectively). **F:** UMAP visualization of the hematopoietic landscape, colored by target fate identity (MgkPro, EryPro) for induced populations from Loop 2 designs. **G:** Predictive uncertainty for MgkPro and EryPro enrichment on the held-out designs, before and after Loop 2. **H:** Measured versus predicted mean (top) and standard deviation (bottom) of marker expression for two held-out experiments (Des. 248, Des. 249), before and after Loop 2. **I:** PCA projection of Ery-enriching experimental designs for Loop 1 (left) and Loop 2 (right), colored by Ery-enrichment uncertainty. All Loop 2-generated designs are outlined in red.

For Loop 2, we expanded the target lineage panel to erythroid progenitors (EryPro), phenotypic HSCs (pHSCs), and pre-ProB cells (Pre-ProB), in addition to MgkPro. Using the updated surrogate forward model from Loop 1, we queried the inverse model to identify conditions predicted to enrich each target fate. For exploitation, we selected one high-confidence candidate per lineage, whereas exploration focused on two additional candidates with high predictive uncertainty but promising predicted enrichment (**Fig. 4B; Supplementary Fig. 8A**). The generated EryPro- and MgkPro-enriching designs selected in Loop 2 displayed diverse concentration patterns and generally distinct compound combinations relative to the corresponding Loop 1 reference design (Des. 210 for MgkPro, Des. 1178 for EryPro) (**Fig. 4C; Supplementary Fig. 8D**). On experimental validation, EryPro conditions reached measured target-cell proportions of 6.1% and 7.4%, ranking within the top 16.3% of all tested designs, while MgkPro conditions reached proportions of approximately 44% and 10%, respectively, ranking within the top 3% of all tested designs (**Fig. 4D**). Consistent with these compositions, the corresponding conditions showed high measured expression of the lineage-specific markers CD235a and CD41a, respectively (**Fig. 4E**). In contrast, pHSC- and Pre-ProB-enriching conditions showed limited enrichment, likely reflecting the absence of key medium parameters required for these fates in the initial design space (**Supplementary Fig. 8B–C**). Finally, UMAP visualization of the cells measured in Loop 2 showed qualitative enrichment of the MgkPro- and EryPro-enriching designs along the megakaryocytic and erythroid branches of the hematopoietic landscape respectively, consistent with their intended cellular fates (**Fig. 4F**).

To address the second objective, we used the high-confidence and one uncertain design per target lineage to update the forward model while holding out the remaining candidates (designs 248 for MgkPro and 249 for EryPro). We then compared predictions from the updated model with those from the original model, which had not seen the new experiments, to assess changes in predictive uncertainty and accuracy. The updated model showed reduced uncertainty for the held-out conditions by 18.4% for MgkPro and 66.1% for EryPro, showing that incorporating new measurements from these conditions reduced uncertainty in previously underexplored regions of the design space (**Fig. 4G**) and improved cell type prediction performance (**Supplementary Fig. 8E**). It also improved prediction of mean marker expression and its variability, increasing the average Pearson correlation with experimental measurements from 0.437 (P <0.05) to 0.873 (P <10^−5^) for the mean; for the standard deviation, the Pearson correlation was not significant before Loop 2, but increased to 0.872 (P<10^−5^) after observing the uncertain measurements (**Fig. 4H**). Together, these results show that incorporating uncertain but informative experiments substantially improved both model confidence and predictive accuracy.

After integrating all three designs per lineage, predictive uncertainty was also reduced in regions of the design space represented by experimental measurements, exemplified by the EryPro and MgkPro enrichment landscapes (**Fig. 4I; Supplementary Fig. 8F**). Overall, Loop 2 showed that LabCompass could identify informative experiments, learn from their outcomes, and improve both model confidence and predictive accuracy in a complex biological system.

### LabCompass integrates new design parameters to enrich rare cell types

The ability to extend the design space with new experimental parameters is crucial for uncovering underobserved regions of the hematopoietic landscape that would remain inaccessible within a static design space. We hypothesized that the initial set of design parameters was insufficient to effectively enrich the poorly represented lineages, such as monocyte and lymphoid branches (**Fig. 2E,G**). For the third iteration (Loop 3), we therefore introduced two additional parameters, M-CSF and a lymphoid (LY) cocktail^49^, to enrich these lineages. Two base designs were supplemented with different LY cocktail concentrations (1.5 and 15 µL/well): 245, generated in Loop 2 to enrich Pre-ProB cells with low uncertainty, and 247, a design selected for its ability to produce diverse cellular fates while retaining undifferentiated states, such as pHSCs and MPP cells (**Supplementary Fig. 9A-B**). Design 247 was also used for M-CSF supplementation, tested at three concentrations (0.2, 2, and 20 ng/mL). We experimentally evaluated these expanded conditions and used the resulting observations to update the forward model and prior (**Fig. 5A**).

**Figure 5.**
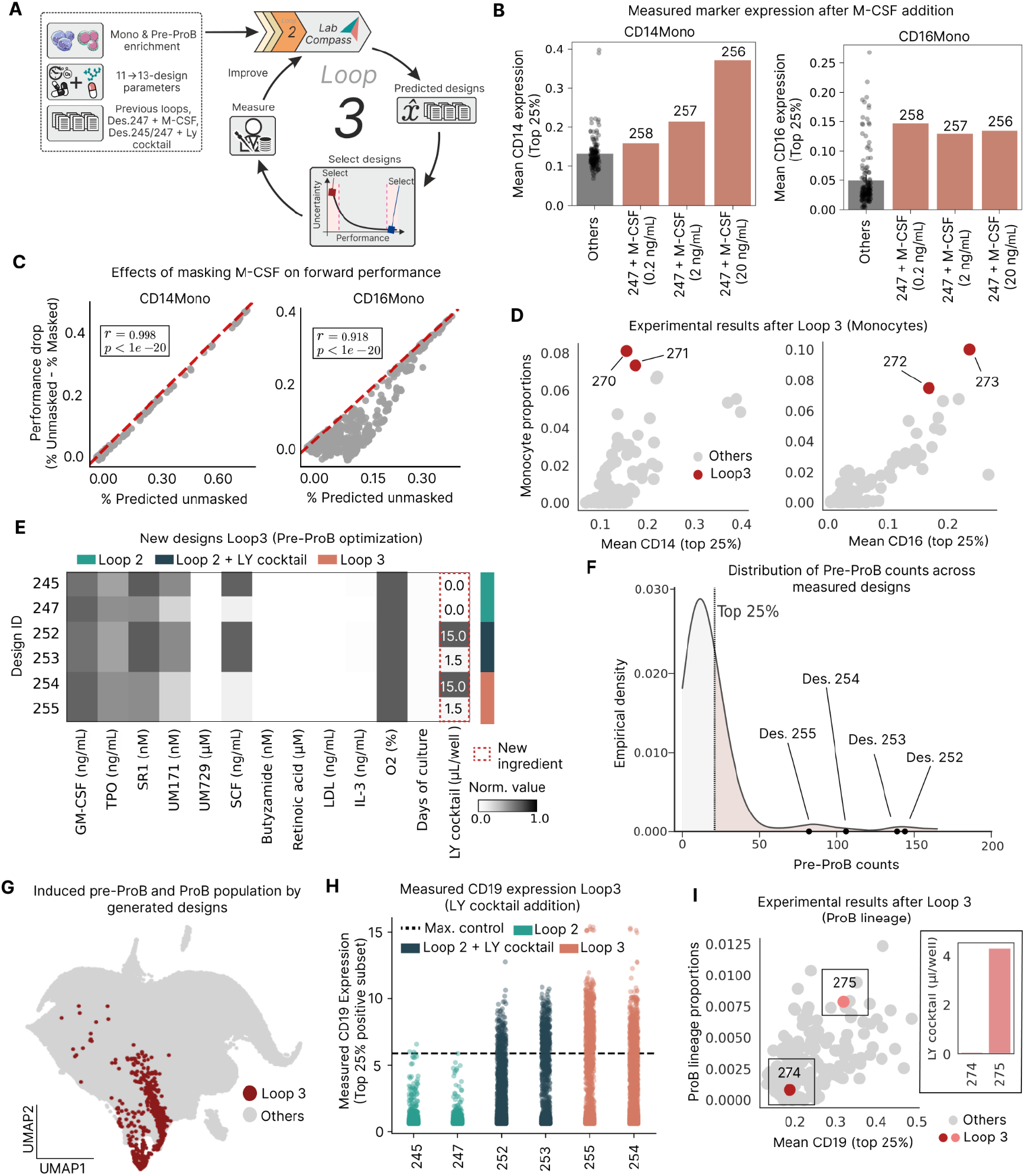
LabCompass enriches rare cell types through the incorporation of new design parameters. **A:** Schematic of Loop 3: new experimental parameters are added to the LabCompass design space to enrich for rare cell types. **B:** Mean CD14 (left) and CD16 (right) marker expression (top 25%) across M-CSF-supplemented conditions. Gray bars show the mean across all other experiments, with individual dots representing single realizations; colored bars show M-CSF conditions at 0.2, 2, and 20 ng/mL added on top of design 247 (Des. 258, 257, 256, respectively). **C:** Counterfactual masked evaluation for CD14Mono- (left) and CD16Mono-enriching (right) LabCompass optimization samples, showing predicted proportion (x-axis) versus the drop in predicted proportion after masking M-CSF (y-axis; red dashed line, bisector). **D:** Measured monocyte proportions versus measured mean CD14 (left) and CD16 (right) marker expression (top 25%) for experimentally validated inverse-design candidates from Loop 3 (red; Des. 270–273) compared with all other designs (gray). **E:** Heatmap of design parameters for the conditions generated for lymphoid (Pre-ProB) enrichment in Loop 3. The LY cocktail was added at two concentration levels (1.5 and 15 µL/well) to two base designs (Des. 245 and Des. 247), which are shown alongside them without supplementation; the new ingredient (LY cocktail) is outlined in red. **F:** Empirical density distribution of Pre-ProB counts across all measured designs. **G:** UMAP visualization computed on the Loop 3 measurements alone, with Pre-ProB/ProB cells from Loop 3 designs highlighted in red against all other Loop 3 cells (gray). **H:** Measured CD19 expression (top 25% of the subset positive for the marker) across the unsupplemented base designs (Des. 245, 247) and the LY-cocktail-supplemented designs (Des. 252, 253, 255, 254), colored by condition (Loop 2, Loop 2 + LY cocktail, Loop 3). Dashed line, maximum expression among positive cells in the control design. **I:** Measured Pre-ProB and ProB proportions versus measured mean CD19 expression (top 25%) for experimentally validated designs, with Des. 274 and Des. 275 (Loop 3) highlighted against all other designs (gray). Inset: LY cocktail concentration (µL/well) used in designs 274 and 275.

M-CSF favors monocyte lineage enrichment^50^, and indeed the M-CSF-supplemented conditions showed higher mean CD14 and CD16 expression than the average across all other designs, with CD14 expression increasing with M-CSF dose (**Fig. 5B, Supplementary Fig. 9D**). We then used the updated forward model to generate designs targeting both monocyte subtypes. To assess whether the predicted enrichment was attributable to M-CSF, we performed an *in silico* counterfactual experiment by masking M-CSF from the generated designs and re-evaluating them with the forward model. The resulting reduction in predicted target proportions strongly correlated with the initial predictions (Pearson r = 0.998 for CD14+ and 0.918 for CD16+; both P < 10^−20^), supporting M-CSF as an important driver of monocyte enrichment (**Fig. 5C**). We experimentally validated two designs for each monocyte subtype, spanning the exploitation–exploration trade-off: low-uncertainty, exploitative designs (270 and 272) and high-uncertainty, exploratory designs (271 and 273) for both monocyte lineages (**Supplementary Fig. 9C**). These designs yielded some of the highest CD14+ and CD16+ monocyte proportions observed in the whole dataset, together with increased marker expression (**Fig. 5D**).

In parallel, we introduced the LY cocktail to address the underrepresented Pre-ProB and ProB lineages (**Fig. 5E**). LY cocktail supplementation increased their abundance, with all supplemented conditions falling within the top 25% of Pre-ProB counts (**Fig. 5F**). These designs also uncovered previously depleted regions of the hematopoietic landscape, with Pre-ProB and ProB cells localizing to the lymphoid-specific branch of an embedding computed on the Loop 3 measurements (**Fig. 5G**), and exhibited increased CD19 expression relative to their unsupplemented base designs (245 and 247) and the control threshold (**Fig. 5H**). We incorporated these results into the forward model and generated new Pre-ProB-enriching designs with LabCompass (**Supplementary Fig. 9C,F**). Following the same exploitation–exploration selection strategy, we validated designs 274 and 275. Notably, only the high-uncertainty design 275, which incorporated the LY cocktail, increased Pre-ProB and ProB abundance and CD19 expression, consistent with LabCompass identifying the newly introduced parameter as an informative axis for lymphoid fate (**Fig. 5I**).

Together, these results demonstrate that LabCompass can integrate new design parameters to uncover and exploit previously inaccessible regions of the hematopoietic landscape, enabling enrichment of rare cell types.

### LabCompass generates alternative routes for *in vitro* cell fate optimization

A key research question in synthetic fate steering is whether models such as LabCompass can identify distinctly different experimental designs that yield the same target cell-type distribution (**Fig. 6A**). Validating this capability would demonstrate that the generative pipeline can move beyond a single solution, enabling broader exploration of the accessible design space. Such alternative routes are of practical value, as protocols reaching the same target population can differ in reagent availability, cost, or achievable efficiency, making some better suited than others to a given experimental setting. We focused on the megakaryocyte-erythroid lineages and CD16 monocytes given their enrichment in previous loops. To build a new up-to-date model, we first pooled datasets generated during prior optimization loops. Secondly, we expanded the design space along an axis suggested by our earlier experiments. High concentrations of the MPL agonist butyzamide contributed to megakaryocyte proliferation, a pattern also captured by the model (**Fig. 3E**), suggesting that TPO concentrations beyond the previous maximum of 2.5 ng/mL provide an alternative route to megakaryocyte enrichment^51,52^. The forward model indeed predicted increased proportions of megakaryocyte/erythroid cells which plateaued beyond 25 ng/mL TPO (**Fig. 6B, Supplementary Fig. 10**). Experimental validation supported this prediction, and the 25 ng/mL TPO condition was incorporated into the training dataset.

**Figure 6.**
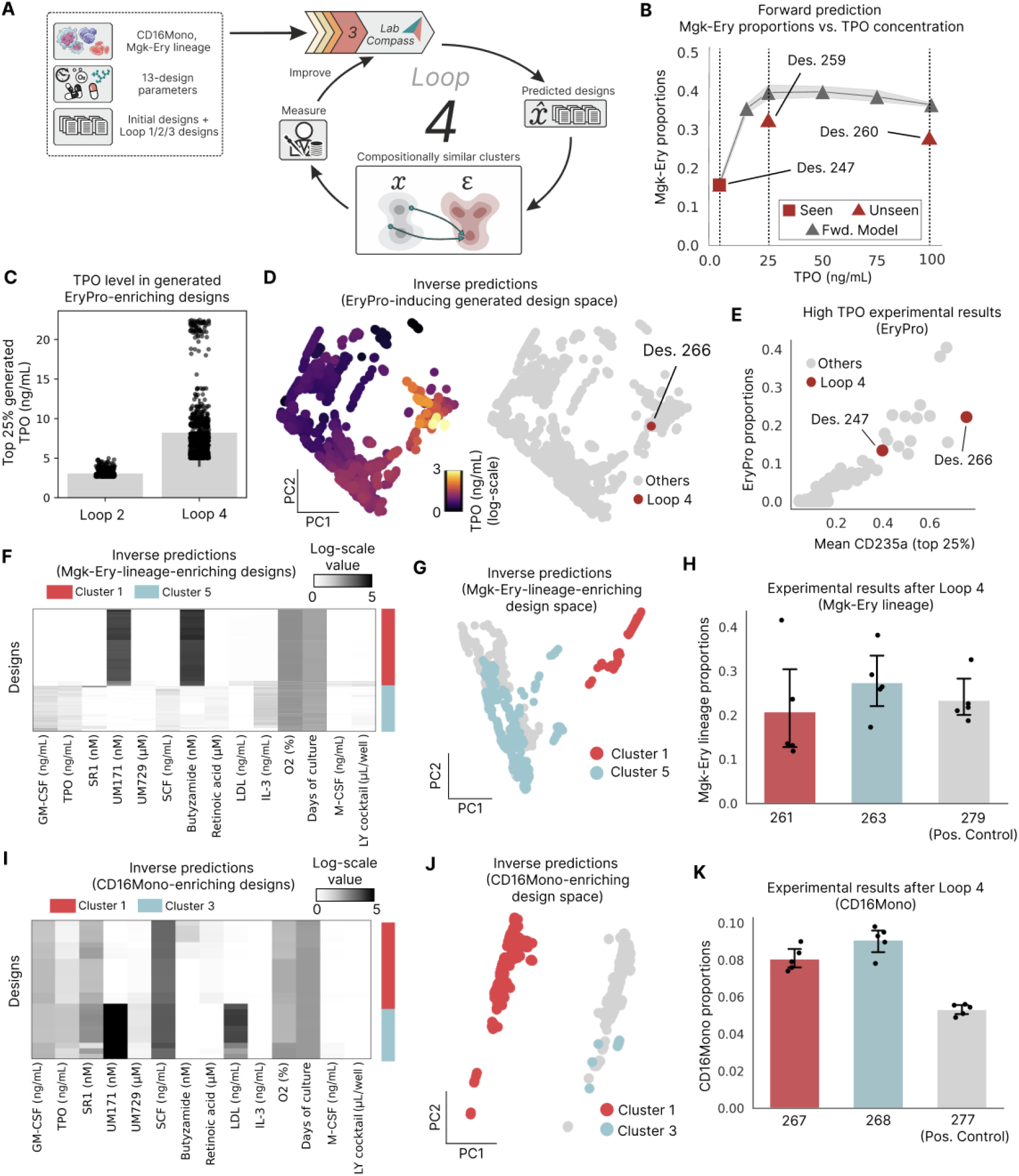
LabCompass identifies alternative differentiation trajectories converging on the same target cell state. **A:** Schematic of Loop 4: LabCompass generates designs predicted to enrich for Mgk-Ery lineages and CD16^+^ monocytes. Generated solutions are hierarchically clustered, and clusters with similar predicted cell-type proportions are compared; we select low-uncertainty solutions from each cluster for experimental validation. **B:** Forward-model predictions (gray triangles) versus measured Mgk-Ery proportions (red) across a titration of TPO concentrations, tested on control design 247. Squares indicate the TPO concentration seen during model training (Des. 247); triangles indicate unseen concentrations predicted by the model and subsequently validated experimentally (Des. 259, Des. 260). Validated experiments at TPO concentrations of 25 or 100 ng/mL were supplemented to base design 247. **C:** Top 25% TPO concentrations (ng/mL) among Ery-enriching designs generated in Loop 2 versus Loop 4. **D:** PCA projection of Ery-enriching designs generated in Loop 4. Left: points colored by predicted TPO log-scale concentration (ng/mL). Right: the same projection with Des. 266 (red), the lowest-uncertainty solution in a generated high-TPO region, highlighted against all other designs (gray). **E:** Measured EryPro proportion versus measured mean CD235a expression (top 25%) for validated recipes from Loop 4 (red) compared with all other validated designs (gray); Des. 266 and the base design 247 are labeled. **F:** Heatmap of design parameters for Mgk-Ery-lineage-enriching designs generated by LabCompass, drawn from two clusters (Cluster 1, Cluster 5) with similar predicted Mgk-Ery proportions. **G:** PCA projection of generated designs with predicted Mgk-Ery proportion > 5%, colored by cluster identity (Cluster 1, Cluster 5). **H:** Measured Mgk-Ery lineage proportions for the lowest-uncertainty solutions from Cluster 1 (Des. 261) and Cluster 5 (Des. 263), compared with the MgkPro-enriching positive control (Des. 279, a biological replicate of design 210). Each point represents one of five technical replicates. **I:** Heatmap of design parameters for CD16Mono-enriching designs generated by LabCompass, drawn from two clusters (Cluster 1, Cluster 3) with similar predicted cell-type proportions. **J:** PCA projection of generated designs with predicted CD16Mono proportion > 2%, colored by cluster identity (Cluster 1, Cluster 3). **K:** Measured CD16Mono proportions for the lowest-uncertainty solutions from Cluster 1 (Des. 267) and Cluster 3 (Des. 268), compared with the CD16Mono-enriching positive control (Des. 277, a biological replicate of design 247). Each point represents one of five technical replicates.

After training LabCompass on all the available data, we queried the model for designs predicted to induce high proportions of megakaryocyte-erythroid lineages and CD16 monocytes (**Supplementary Fig. 11A-F**), clustered the proposed solutions, and identified pairs of clusters with distinct parameter compositions but similar predicted cell type distributions. Low-uncertainty solutions were then selected for experimental validation. This analysis revealed a new optimization axis for erythroid induction: newly LabCompass-generated designs used TPO concentrations above those of the Loop 2 erythroid designs, which had remained near the original 2.5 ng/mL ceiling (**Fig. 6C**). Megakaryocyte enrichment was also affected, though to a lesser extent (**Supplementary Fig. 12A-B**). For both cell types, the addition of new TPO dosages led to an overall forward uncertainty reduction across the design space (**Supplementary Fig. 12C**). We validated design 266, the lowest-uncertainty solution among those exceeding the TPO concentrations used in Loop 2 (8.13 ng/mL), which occupied a distinct region of the design space (**Fig. 6D, Supplementary Fig. 12D-E**). Experimentally, it ranked among the top designs for CD235a expression and increased the erythroid population to 22%, roughly a 1.6-fold increase over the control condition (Des. 247) (**Fig. 6E**).

More broadly, clustering of the generated designs revealed distinct routes toward similar target distributions. For megakaryocyte-erythroid lineage optimization, two clusters exhibited similar predicted cell type proportions but distinct parameter compositions (**Fig. 6F-G, Supplementary Fig. 13A-B**). One route resembled design 210, combining UM171 and high butyzamide, whereas the alternative combined TPO and IL-3 (previously shown to promote megakaryocyte enrichment)^47,53^, lacked UM171, and had low butyzamide. The model also identified the LY cocktail as a component in some clusters, albeit at marginal levels (**Supplementary Fig. 13A)**. Notably, the LY cocktail contains IL-3 and SCF, which promote megakaryocyte and erythroid differentiation and proliferation *in vitro*^*54*^, suggesting that the model independently recovered a biologically relevant combination. The lowest-uncertainty designs from these clusters (designs 261 and 263) were experimentally validated and produced similar cumulative proportions of megakaryocytes and megakaryocyte-erythroid progenitors, achieving a mean enrichment between 20% and 30%, comparable to the MgkPro-enriching positive control (design 210 – here run as an independent biological repeat 279) (**Fig. 6H, Supplementary Fig. 14A-B**).

A similar analysis for CD16+ monocytes identified two clusters predicted to induce 2%–4% monocyte enrichment, differing primarily in the prevalence of Low Density Lipoproteins (LDL) and UM171 (**Fig. 6I-J, Supplementary Fig. 13C-D**). Designs 267 and 268 were experimentally validated, yielding similar mean CD16+ monocyte enrichments of 7.8% and 8.7%, respectively, both exceeding the positive control (design 247 - here run as an independent biological repeat 277) and the predicted range (**Fig. 6K, Supplementary Fig. 14A,C**).

Together, these results demonstrate that LabCompass can identify distinct experimental routes beyond trivial near-neighbors that converge on similar cellular outcomes, highlighting its potential for broader exploration of lineage optimization.

## Discussion

Developing new hematopoietic differentiation protocols is traditionally slow and material-intensive: the current gold-standard protocol for cord blood-derived megakaryocytes, for example, was distilled from 1,500 donor samples^26^. Here, LabCompass, a generative model conditioned on a target cell state, proposed designs that were tested in the laboratory, with each round of measurements improving the model that proposed the next. Within just four such loops, this approach recovered a known megakaryocyte-inducing condition without having observed it. It also incorporated new cytokines into the design space and optimized for CD16+ monocytes and early lymphoid lineages that were rare in the starting screen.

Strikingly, for the fates it reached, LabCompass showed the answer is several distinct routes rather than one: MgkEry (joint enrichment of MgkPro, MgkEryPro1, MgkEryPro2 and EryPro) and CD16+ monocyte lineages could each be steered via distinct signal combinations converging on the same target state. This has real practical value: if a specific molecule is prohibitively expensive, chemically unstable, or immunogenic for clinical manufacturing, inverse modeling can generate functionally equivalent recipes that substitute reagents while maintaining target cellular yield.

LabCompass’s forward predictions are also aligned with known biology. *In silico* M-CSF masking recovered a known biological dependency, and the model correctly predicted that TPO enrichment would plateau at higher doses, a pattern more consistent with an underlying dose-response mechanism than with simple linear extrapolation. This suggests that, given a mechanistic hypothesis, *in silico* predictions can be used to reduce experimental burden and reagent costs. Although the predictive accuracy of perturbation-response models remains an open challenge in the field^55^, we show that even potentially imperfect forward models can identify biologically meaningful directions in the design space, drive effective experimental optimization, and improve over successive loops.

By explicitly modeling uncertainty in the system’s response, LabCompass can identify promising unexplored regions of the experimental space and prioritize potentially informative experiments to refine its predictions. This feedback loop enables principled, model-guided exploration of complex biological systems and could improve experimental efficiency. It also connects experimental design to model-based sequential decision-making: the forward surrogate serves as a learned world model of the culture system, while the guided generative model proposes candidate experiments. Here, a simple selection rule balanced exploiting confident predictions with exploring uncertain regions; active learning strategies could adapt this balance across loops. Because the same factor can favor expansion or differentiation depending on timing, extending designs to time-varying dosing schedules would introduce a further challenge: choosing interventions whose effects depend on the current cellular state and influence subsequent responses. Such sequential control offers a natural direction for model-based reinforcement learning, using world models of cellular dynamics to optimize protocols over time.

This combination of forward prediction and inverse generation sets LabCompass apart from existing approaches, which typically select or optimize among predefined interventions. Rather than searching or optimizing over a fixed candidate space, generative inverse design proposes promising designs directly, reducing the computation needed to reach a target state. Lookup-based methods are constrained by their candidate space and do not naturally accommodate continuous dosages^19–21,46^; Bayesian optimization provides uncertainty estimates but relies on a probabilistic surrogate rather than a generative model^16,18,56^; and gradient-based inverse design offers greater flexibility but, without a generative prior, can move into regions unsupported by real designs, with plausibility assessed only after optimization^48^. In contrast, LabCompass optimizes within a learned generative model, constraining the optimization trajectory to plausible designs.

The LabCompass framework demonstrated here for human hematopoietic *in vitro* differentiation is broadly applicable to other experimental systems, including other stem cell differentiation systems, organoid cultures, and drug screening. It is also agnostic to the experimental readout. Applied instead to single-cell RNA sequencing data from human cytomegalovirus infection^57^, LabCompass recovered held-out infection phenotypes and identified host restriction factors, including STAT2 and IFNAR2, benchmarked against real Perturb-seq knockdown data (**Supplementary Fig. 2 and Methods**). We used spectral flow cytometry because it provides low-cost, same-day, batch-correction-free, multiparametric, and high-throughput measurements; but the flexible forward model has now also been shown to extend to transcriptomic readouts, and could in principle extend to image-based features^58^ as well.

As with any data-driven model, LabCompass’s performance depends on the quality and diversity of its training data and the presence of informative signals in the measurements. As a deep learning model, it also provides limited mechanistic interpretability. Our current implementation retrains the forward surrogate with newly acquired data at each experimental loop, introducing a computational burden that could be reduced through more efficient active-learning or fine-tuning strategies. Furthermore, LabCompass can only identify target cell states supported by the available design space: as shown in Loop 3, desired lineages could only be reached when the necessary experimental parameters were included. In such cases, prior knowledge remains essential for guiding the expansion of the design space. For example, pHSC enrichment was limited in this study, potentially reflecting the challenge of promoting HSC proliferation while simultaneously preserving self-renewal and stemness^23^. Nevertheless, incorporating new molecules that target HSC expansion pathways beyond those already represented might further improve enrichment of this rare population^23,25,59,60^.

Several directions remain for future development. In this study, LabCompass was queried with a single target cell type at a time. The underlying framework is not restricted to this setting, and extending queries to flexible, multi-target objectives, for example jointly enriching several lineages or balancing competing cell states, is a natural extension. The same holds for the design space, which can in principle combine discrete and continuous parameters, as long as the generative model can represent them. Expanding the design space to include the signals needed for full neutrophil and lymphoid maturation, as well as building on the progenitor states already reached, would be a natural next discovery test of this approach. More broadly, sequential inverse modeling could enable data-driven computational representations of cell culture systems, providing a step towards a digital twin of human *in vitro* hematopoiesis that could ultimately help guide the manufacture of clinically relevant primary human cell types that remain difficult to produce at scale. Finally, LabCompass could be coupled with language models at the design selection step, extending the current objective beyond cell-type proportions toward literature-grounded criteria in which generated candidates are assessed against published protocols, mechanistic plausibility, and practical constraints on their deployment^61^.

Realizing this vision requires evaluating virtual cell models not only by predictive accuracy, but by their ability to guide experiments toward specified biological outcomes. Standard forward models are typically benchmarked on predictive accuracy, a mathematically clean evaluation of individual predictions, but one that is practically insufficient for guiding experiment selection. With inverse modeling, predictive accuracy becomes a means to an end rather than the ultimate benchmark, enabling even imperfect models to drive successful experimental optimization. Systematically benchmarking models on inverse tasks will clarify their capacity to capture biologically relevant properties. Embedded in closed experimental loops alongside multimodal readouts, this approach will dramatically accelerate biological discovery.

## Supplementary Figures

**Supplementary Figure 1.**
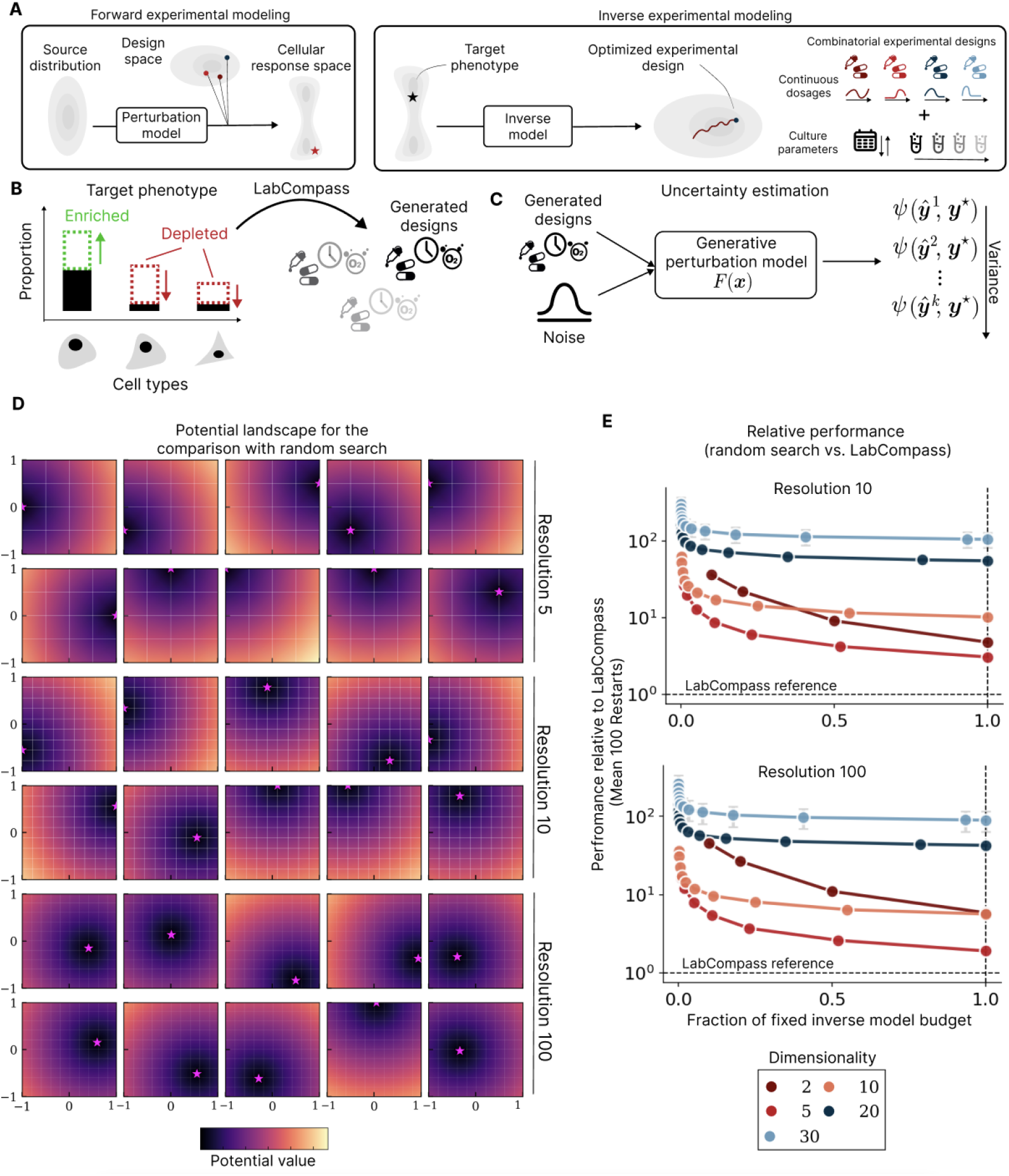
Forward and inverse experimental modeling in LabCompass, and benchmarking against random search. **A:** Schematic of forward and inverse perturbation modeling. Given an experimental design, the forward model maps a source distribution to the cellular response space. The inverse model takes a target phenotype as input, and traverses the experimental design space, including continuous dosages and culture parameters, to yield candidate designs. **B:** Representation of an inverse query. The target phenotype is a vector representing the desired cell-type proportions, with higher values for cell types to enrich and lower values for those to deplete. This vector is used to query LabCompass for candidate designs. **C:** Forward model uncertainty estimation. Generated designs, together with sampled noise, are passed through the generative perturbation model to produce predicted responses. Predicted responses are compared to the target phenotype using a loss function, and the variance of the loss across samples is used to rank generated designs. **D:** Potential landscape used for the comparison with uniform random search, shown in two dimensions across three discretization resolutions (5, 10, 100). Each panel shows one instance of the landscape, colored by potential value from low (dark) to high (light), with grid lines marking the discretization at each resolution and the magenta star marking the optimal design. **E:** Performance of random search relative to LabCompass on the synthetic grid-optimization problem, at resolutions of 10 and 100. The x-axis shows the fraction of the fixed inverse-model budget used by random search, and the y-axis shows random search’s performance relative to LabCompass, averaged over 100 restarts. Colors indicate the dimensionality of the design space (2 to 30), and LabCompass is shown as a fixed-budget reference, marked by the dashed horizontal line at 1.

**Supplementary Figure 2.**
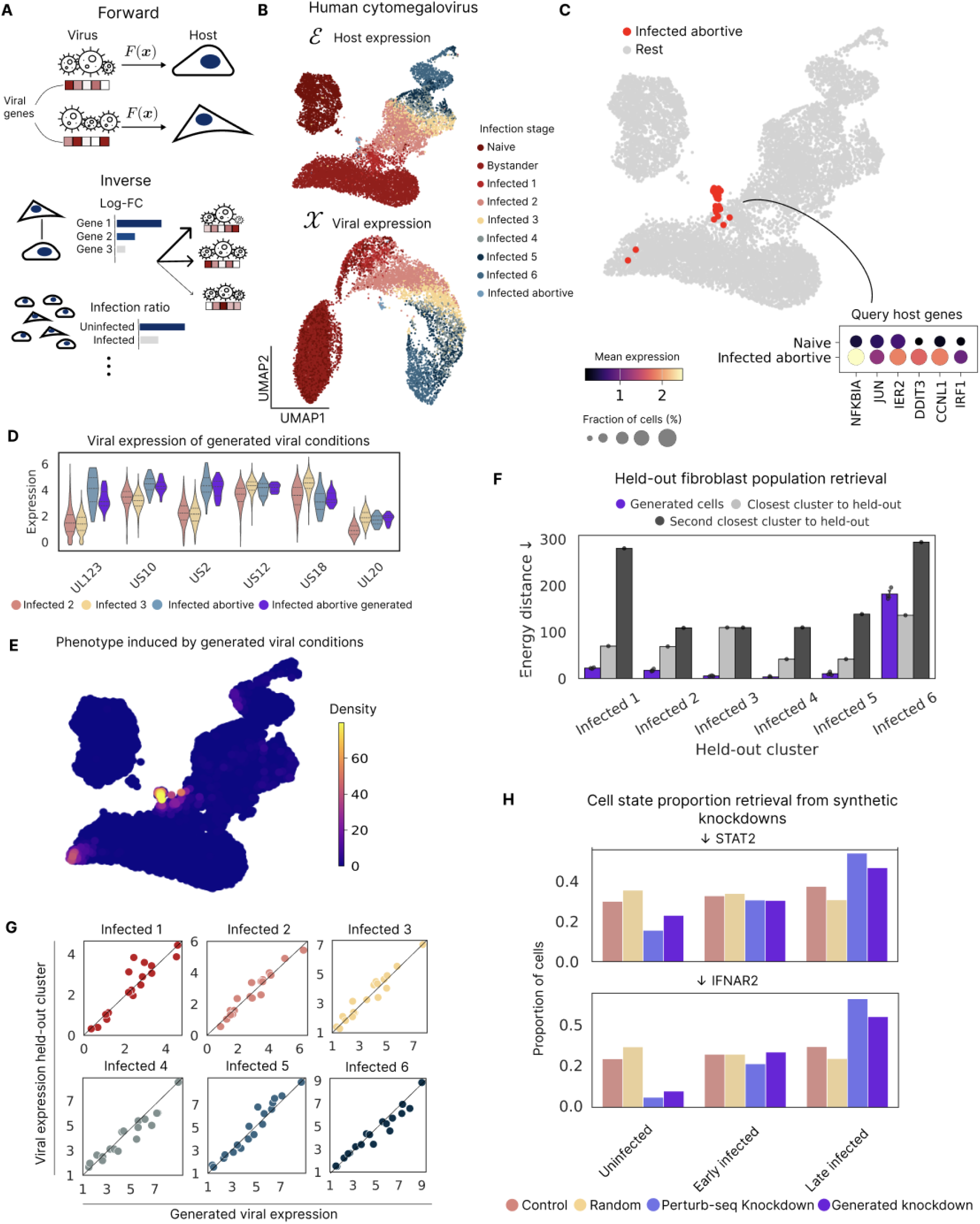
Inverse design recovers infection phenotypes and identifies host restriction factors in HCMV (human cytomegalovirus) infection. **A:** Schematic of the forward and inverse modeling tasks. LabCompass was applied to a published single-cell RNA sequencing time course of HCMV infection in human fibroblasts, profiled across 8 time points, together with its accompanying Perturb-seq host-factor screen^57^. Here the design space is the viral expression state of a cell and the cellular state space is its host expression profile, so the inverse task is to find viral expression states that induce a specified host transcriptional phenotype, specified as log_2_ fold changes of query host genes relative to naive cells. **B:** UMAP of host (top) and viral (bottom) expression, colored by infection stage (Naive, Bystander, Infected 1–6, Infected abortive), for the complete dataset. **C:** Infected abortive cells (red) versus all others (gray), with a dot plot of the query host genes used to define this target phenotype (NFKBIA, JUN, IER2, DDIT3, CCNL1 and IRF1). **D:** Viral gene expression across real Infected 2, Infected 3, and Infected abortive cells versus cells generated for the Infected abortive target, for a panel of HCMV genes spanning the canonical kinetic classes. **E:** UMAP density of cells generated for the Infected abortive target within the host expression landscape. **F:** Energy distance between generated cells and each held-out cluster, compared with the closest and second-closest real clusters and with a random subset of training viral states. Each of the clusters Infected 1 to 6 was removed in turn and the embeddings, prior and forward model refitted on the remaining data, so generated states were absent from training. Bars show the mean over three replicates with standard error. **G:** Held-out versus generated viral expression for each held-out cluster, compared gene by gene after decoding to gene space. **H:** Infection-stage composition for STAT2 (top) and IFNAR2 (bottom), comparing cells generated by guiding the viral state toward host states lacking the target gene with cells carrying the corresponding guide in the Perturb-seq screen, cells carrying non-targeting guides, and an equally sized random sample of screen cells. Stages are grouped as Uninfected, Early infected and Late infected.

**Supplementary Figure 3.**
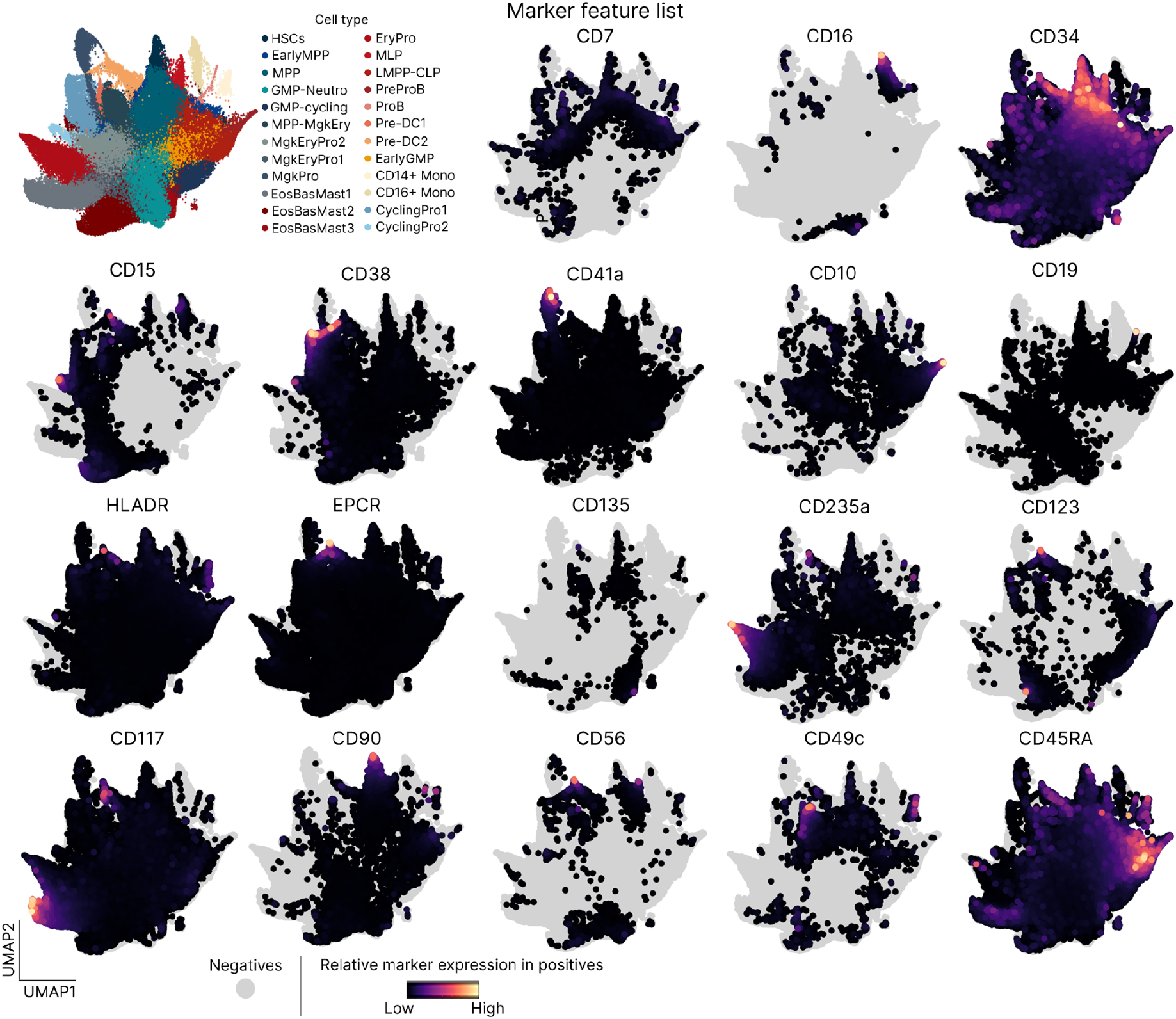
Marker expression across the hematopoietic landscape. Top left: UMAP of the combined hematopoietic landscape from all initial 116 experimental designs, colored by cell type, shown for reference. Remaining panels: the same UMAP, with cells positive for each of 18 markers in the immunophenotypic spectral flow panel colored by relative marker expression among positive cells, and negative cells in gray.

**Supplementary Figure 4.**
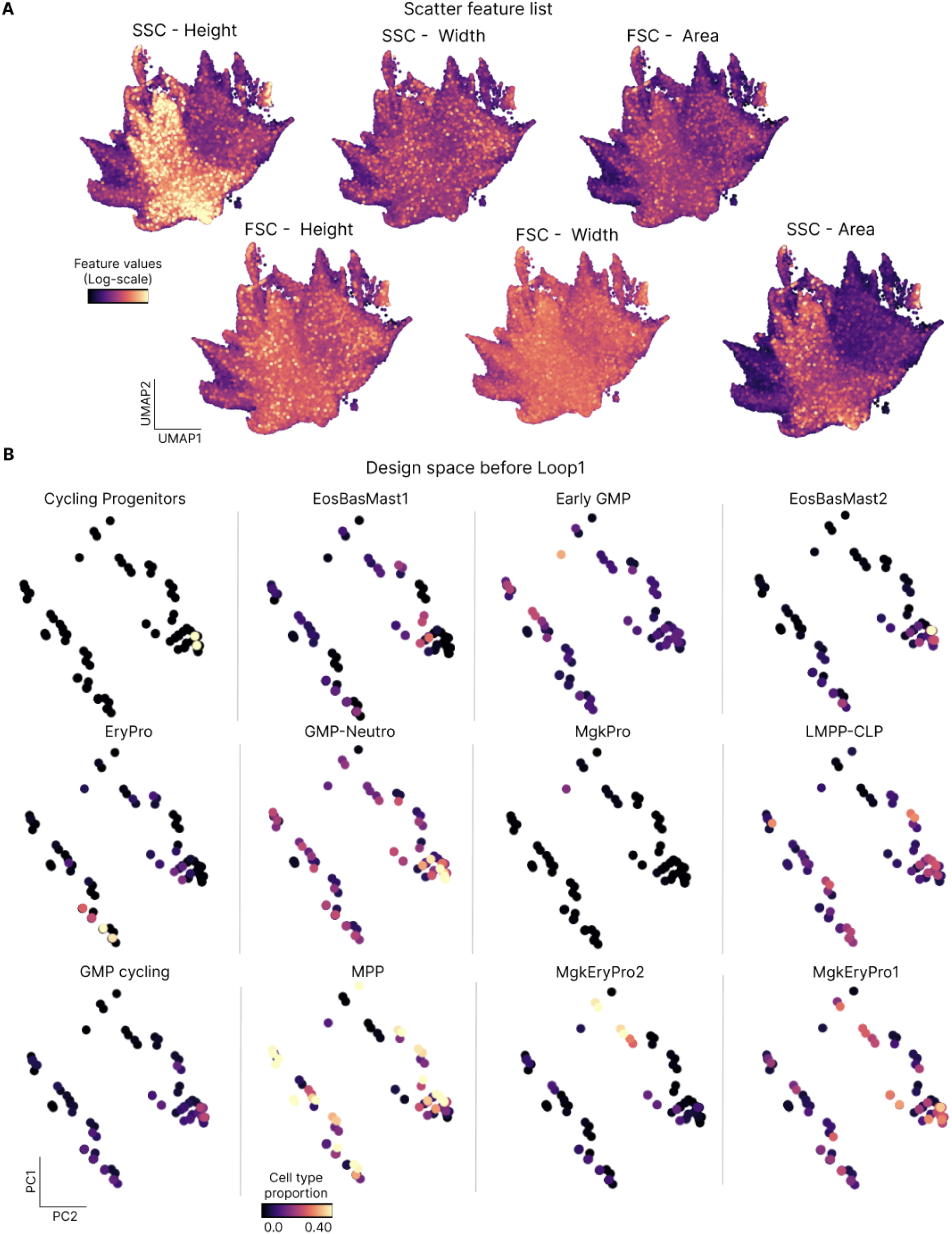
Morphological features across the hematopoietic landscape and cell-type localization in the initial design space. **A:** UMAP visualization of the hematopoietic landscape, colored by the morphological (scatter) features captured by spectral flow cytometry (SSC-Height, SSC-Width, FSC-Area, FSC-Height, FSC-Width, SSC-Area). **B:** Two-dimensional principal component projection of the initial design space, colored by the induced proportion of each cell type.

**Supplementary Figure 5.**
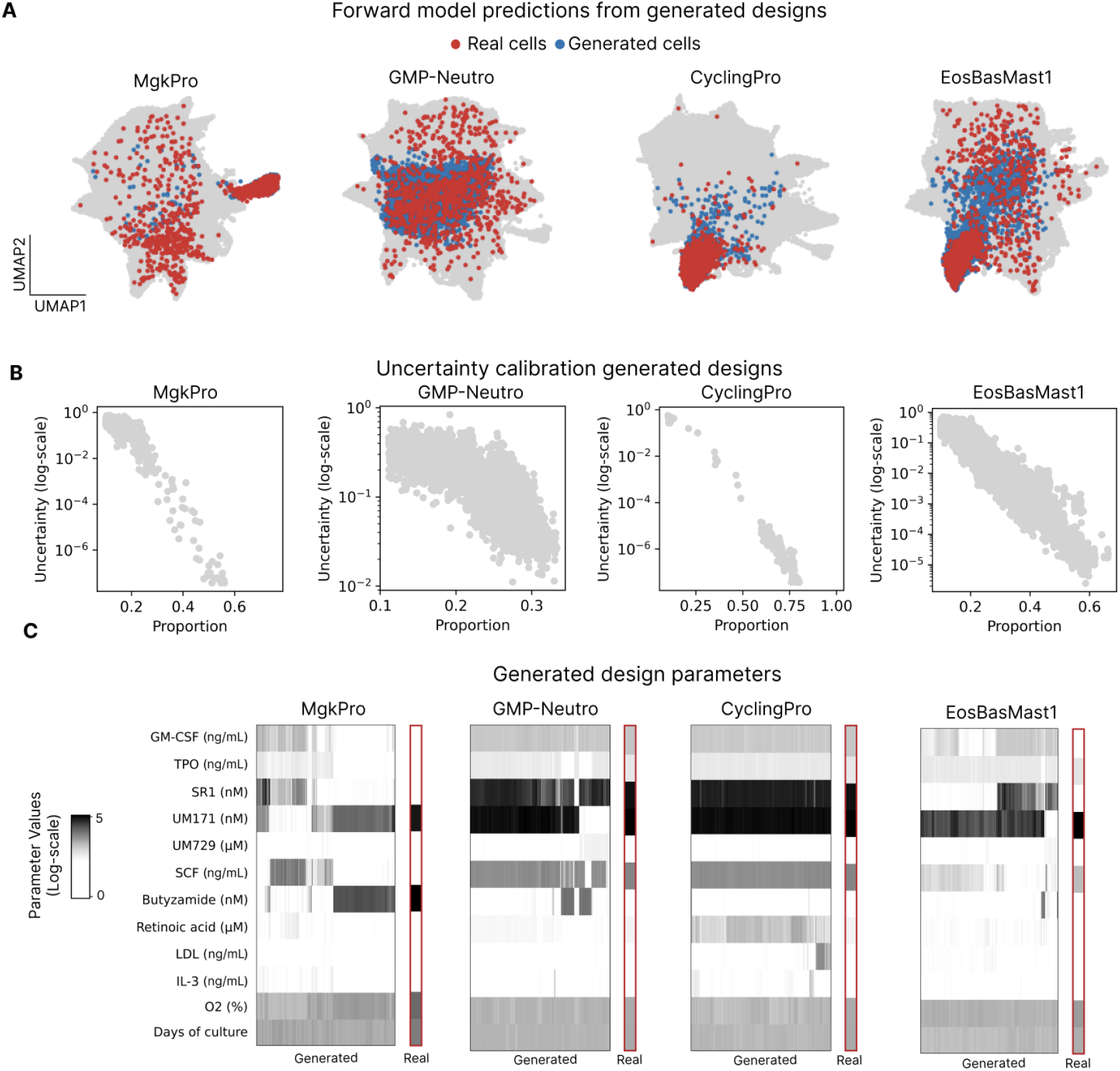
orward model predictions and inverse design generation for MgkPro-, GMP-Neutro-, CyclingPro-, and EosBasMast1-enriching conditions. **A:** UMAP visualization of the hematopoietic landscape, showing real cells (red) and generated cells (blue) for each target cell type before Loop 1. **B:** Predicted target cell-type proportion (x-axis) versus the forward model’s uncertainty (y-axis, log scale) for each target cell type. **C:** Heatmap of culture parameters for LabCompass-generated designs enriching MgkPro, GMP-Neutro, CyclingPro, and EosBasMast1, compared with real experimental designs (right-hand column, outlined in red).

**Supplementary Figure 6.**
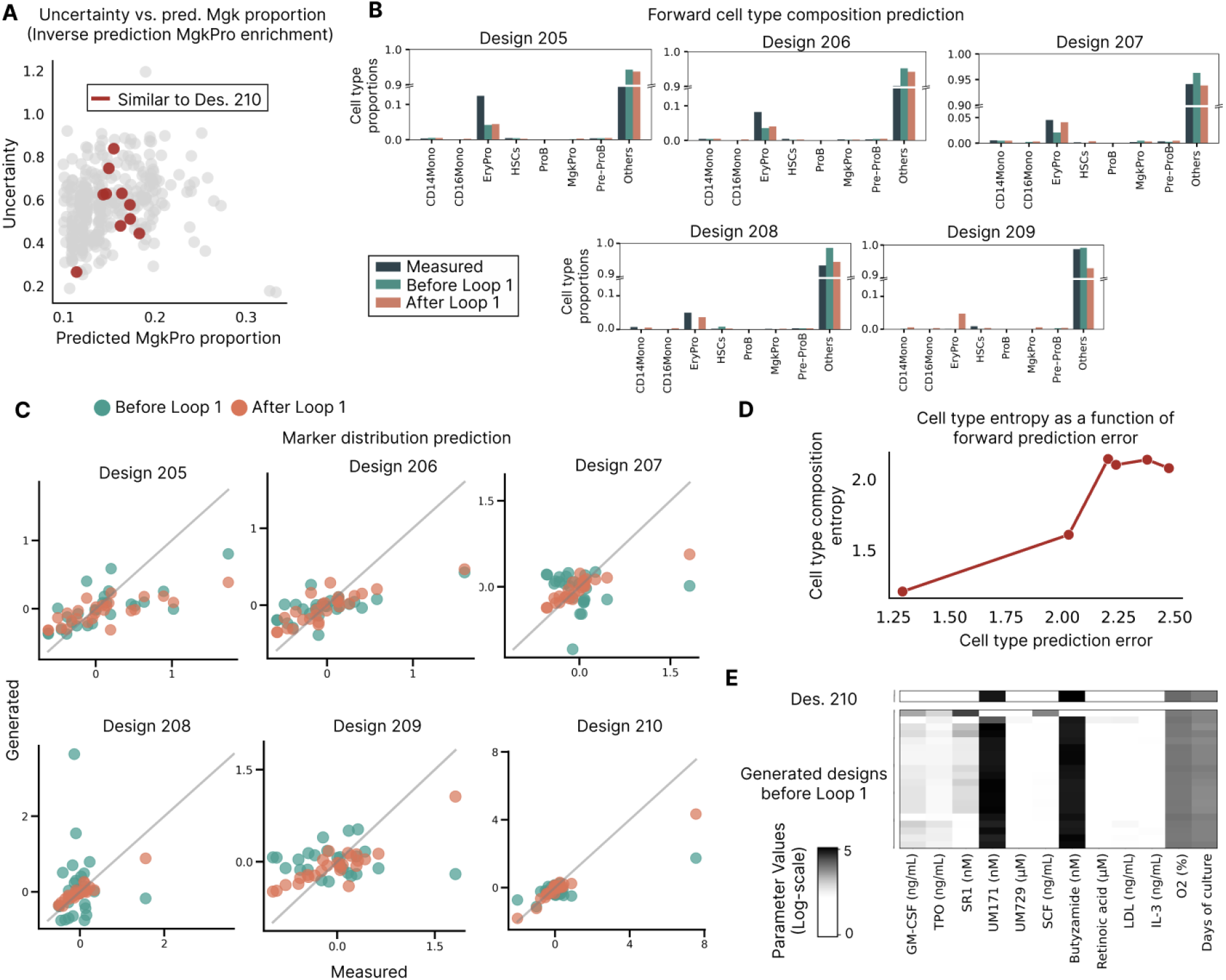
orward model uncertainty, marker prediction accuracy, and design similarity to experiment 210 across held-out experiments before and after Loop 1. **A:** Predicted MgkPro proportion (x-axis) versus the uncertainty of the forward model trained before Loop 1 (y-axis). **B:** Measured cell-type composition compared with model predictions before and after Loop 1, for each of the five held-out designs (205–209). **C:** Measured versus predicted mean marker expression for the held-out experiments (205–210), before and after Loop 1. **D:** Cell-type entropy of the predicted composition as a function of forward-prediction error across held-out designs. **E:** Heatmap of culture parameters for design 210 compared with the set of generated designs preceding Loop 1, highlighting parameter ranges (e.g., UM171, butyzamide) shared between design 210 and the generated designs.

**Supplementary Figure 7.**
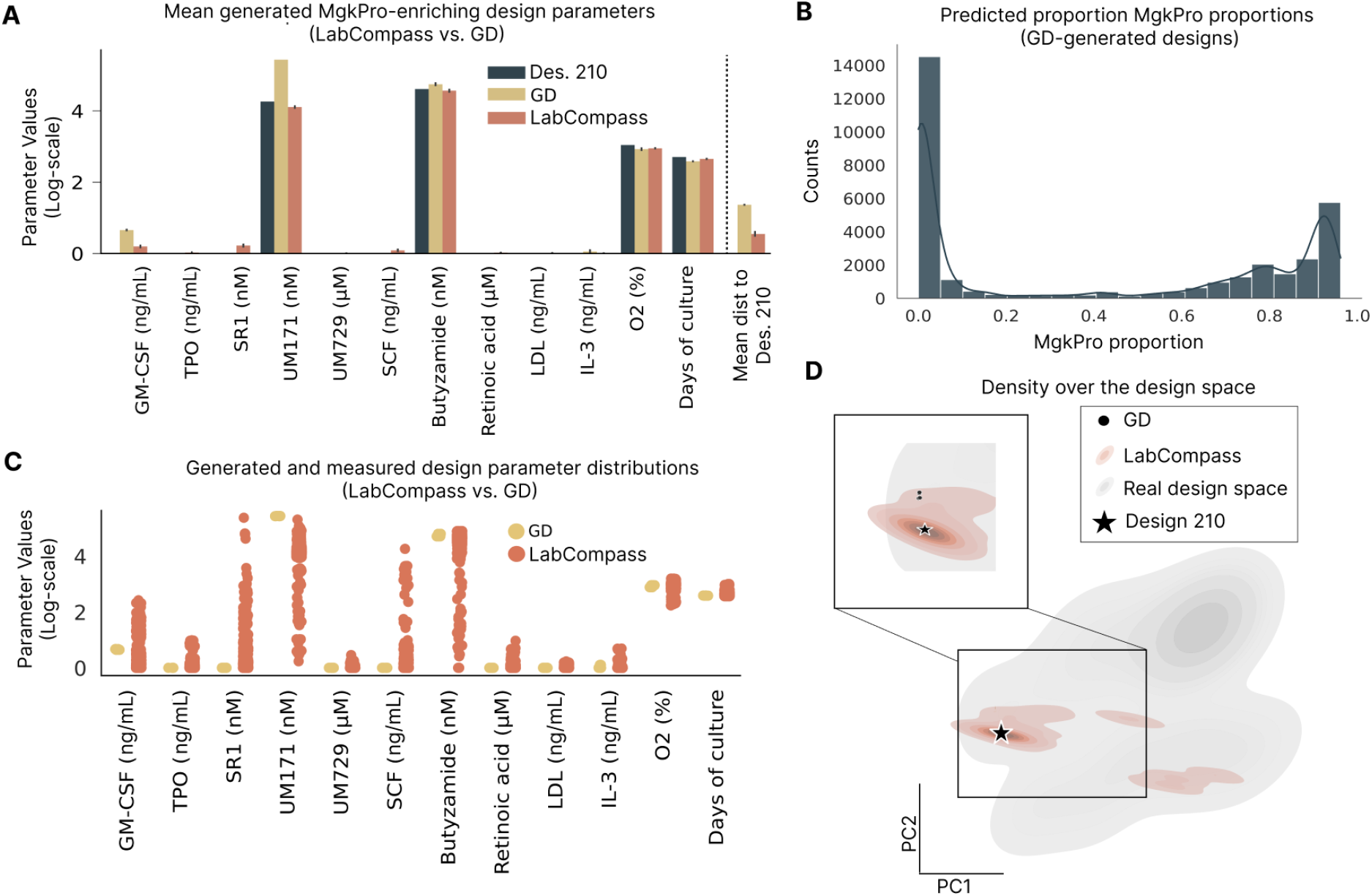
Comparison of gradient-descent (GD) and LabCompass-generated designs for MgkPro enrichment after Loop 1. **A:** Mean value of each design parameter for GD- and LabCompass-generated designs, compared with design 210, alongside the mean Euclidean distance to generated designs to design 210. **B:** Distribution of predicted MgkPro proportion across designs generated by GD. **C:** Distribution of individual design-parameter values for GD- and LabCompass-generated designs. **D:** Two-dimensional principal component projection of the design space, showing projected densities of the real design space (gray) and of LabCompass-generated designs (red), with GD solutions overlaid as individual points (black). The star indicates the position of design 210. GD solutions collapse to a single location and are therefore indistinguishable at this scale. The inset shows a zoomed view of that region, where the GD solutions and the densest LabCompass solutions overlap.

**Supplementary Figure 8.**
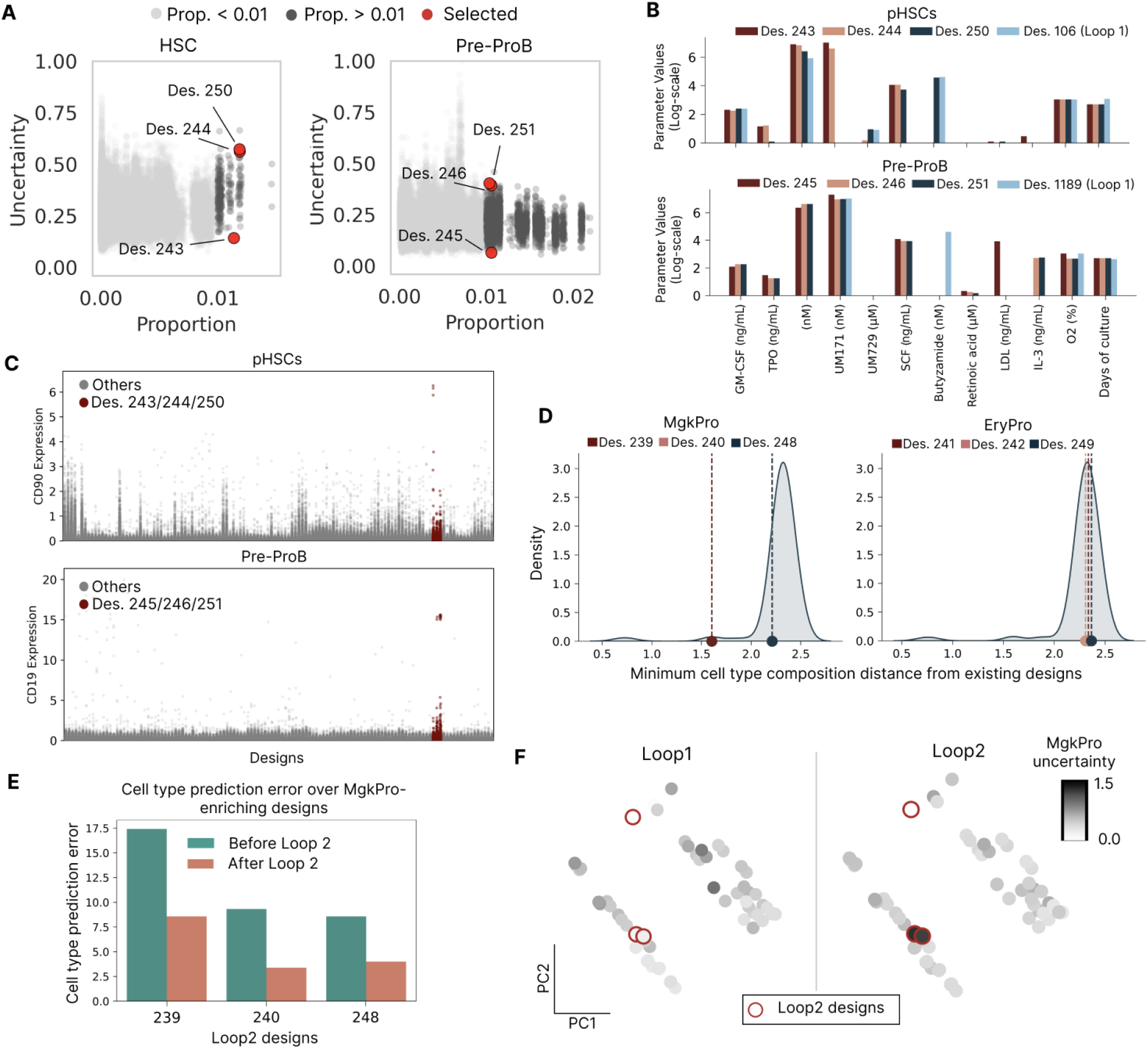
Design selection, marker expression, and prediction quality for HSC-, Pre-ProB-, MgkPro-, and EryPro-enriching designs in Loop 2. **A:** Predicted proportion (x-axis) versus forward-model uncertainty (y-axis) for pHSC- (left) and Pre-ProB-enriching (right) designs, shaded by predicted proportion (light gray, prop. < 0.01; dark gray, prop. > 0.01), with selected designs highlighted in red. **B:** Log-scale concentration of each design parameter for selected pHSC designs (Des. 243, 244, 250) and the most pHSC-enriching design from Loop 1 (Des. 106; top), as well as selected Pre-ProB designs (Des. 245, 246, 251) and the most Pre-ProB-enriching design from Loop 1 (Des. 1189; bottom). **C:** CD90 expression (top, pHSC-selected designs) and CD19 expression (bottom, Pre-ProB-selected designs) across all designs (gray), with selected designs highlighted in red. **D:** Density distribution of minimum cell-type composition Euclidean distance from existing designs for LabCompass-generated MgkPro-enriching (left; Des. 239, 240, 248) and EryPro-enriching (right; Des. 241, 242, 249) designs, with each selected design marked by a dashed line. **E:** Cell-type prediction error measured as cross entropy for LabCompass-generated MgkPro-enriching designs (239, 240, 248), shown before and after Loop 2. **F:** PCA projection of experimental designs for Loop 1 (left) and Loop 2 (right), colored by MgkPro uncertainty, with Loop 2 designs outlined in red.

**Supplementary Figure 9.**
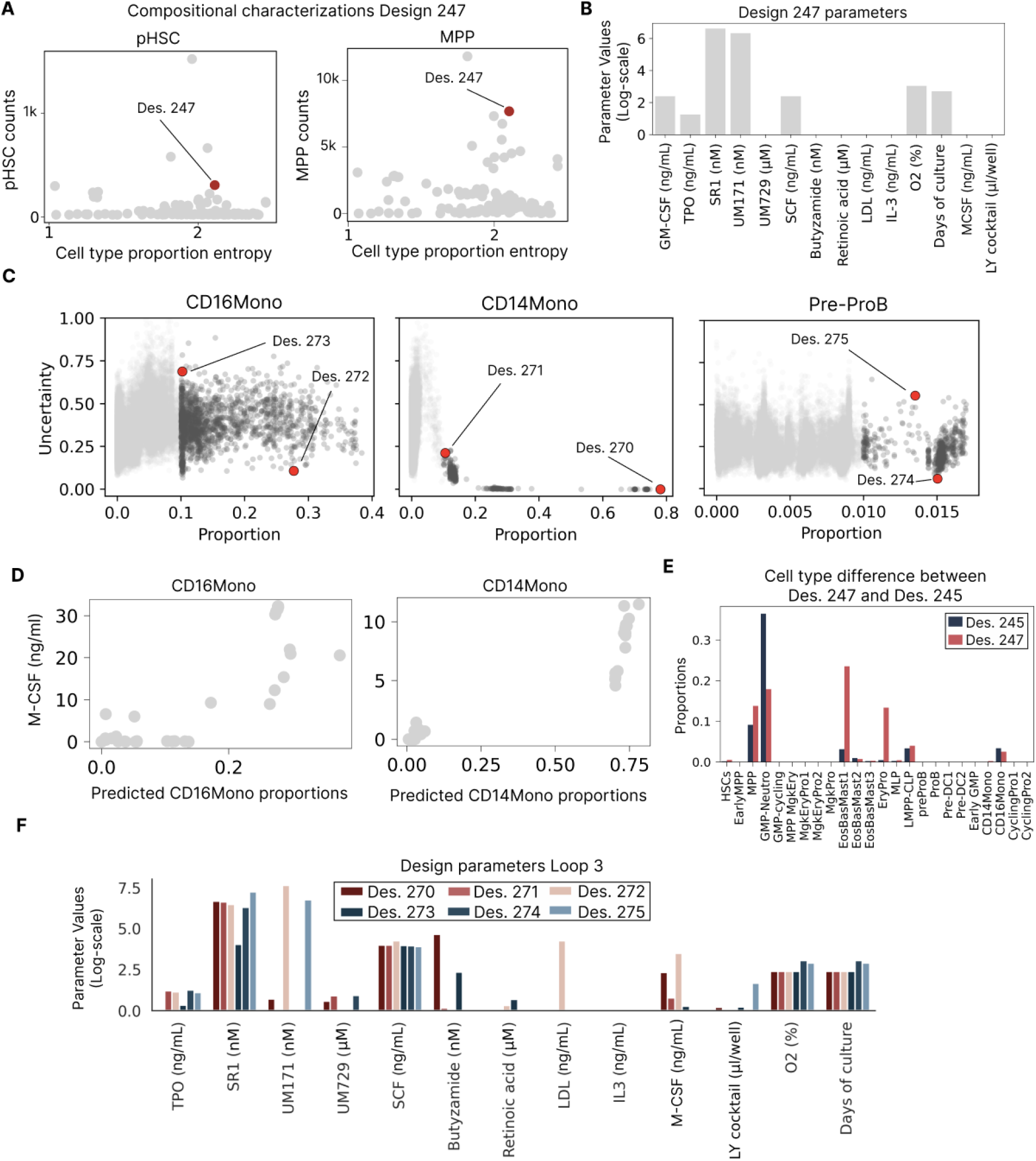
Design selection and parameter analysis for HSC/MPP, CD16Mono-, CD14Mono-, and Pre-ProB-enriching designs in Loop 3. **A:** HSC counts (left) and MPP counts (right) versus cell-type proportion entropy across designs, with design 247 highlighted in red. **B:** Log-scale value of each design parameter for design 247. **C:** Predicted proportion (x-axis) versus forward-model uncertainty (y-axis) for CD16Mono- (left), CD14Mono- (middle), and Pre-ProB-enriching (right) designs, with selected designs (Des. 270–275) highlighted in red. **D:** M-CSF concentration versus predicted CD16Mono proportion (left) and predicted CD14Mono proportion (right) across designs. **E:** Cell-type proportion difference between design 245 and design 247 across all cell types. **F:** Log-scale concentration of each design parameter for generated and measured designs (Des. 270–275).

**Supplementary Figure 10.**
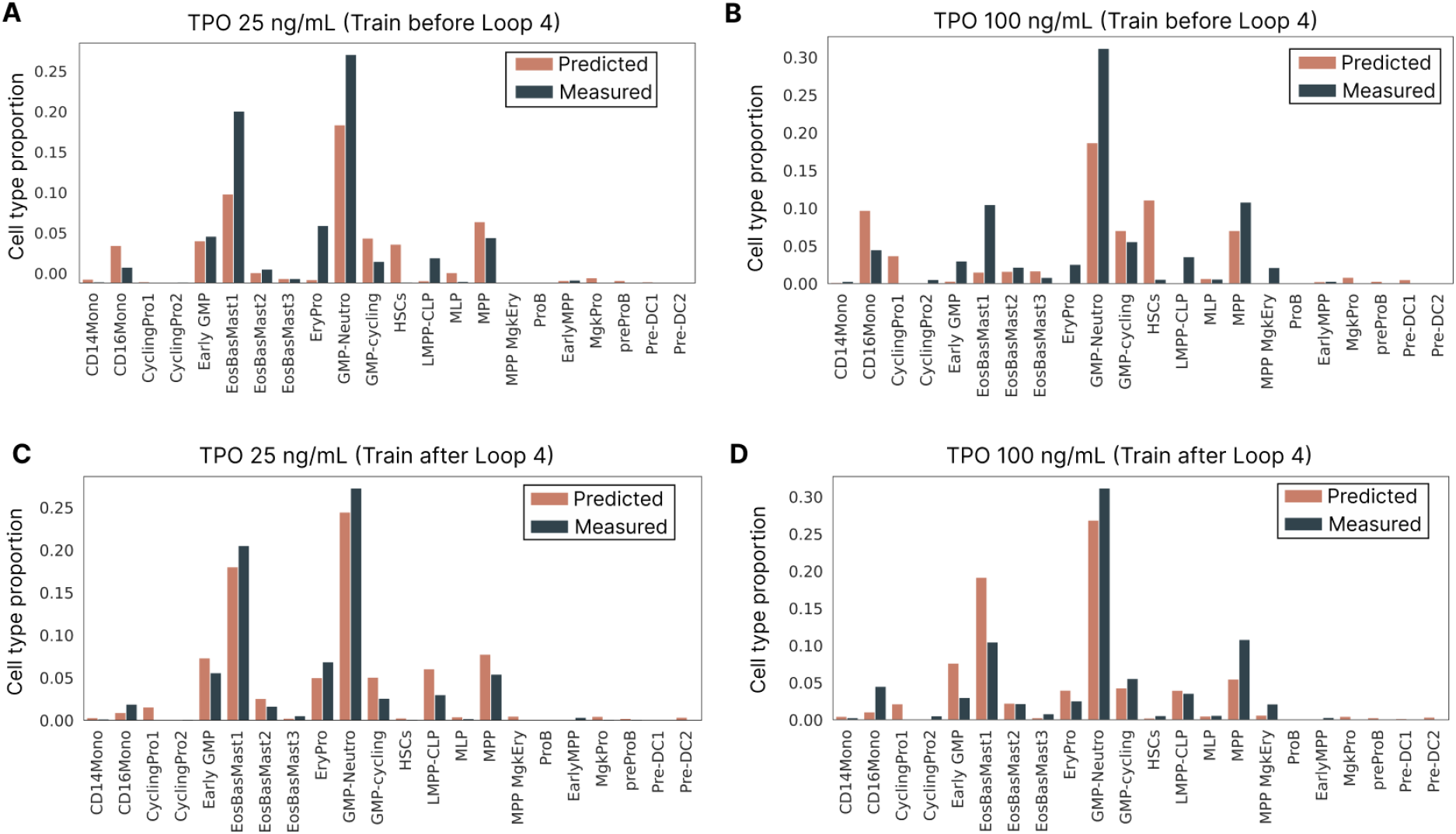
Predicted versus measured cell-type composition at TPO 25 ng/mL and TPO 100 ng/mL, before and after Loop 4. **A:** Predicted and measured cell-type proportions at TPO 25 ng/mL, using the forward model trained before Loop 4. **B:** Predicted and measured cell-type proportions at TPO 100 ng/mL, using the forward model trained before Loop 4. **C:** Predicted and measured cell-type proportions at TPO 25 ng/mL, using the forward model trained after Loop 4. **D:** Predicted and measured cell-type proportions at TPO 100 ng/mL, using the forward model trained after Loop 4.

**Supplementary Figure 11.**
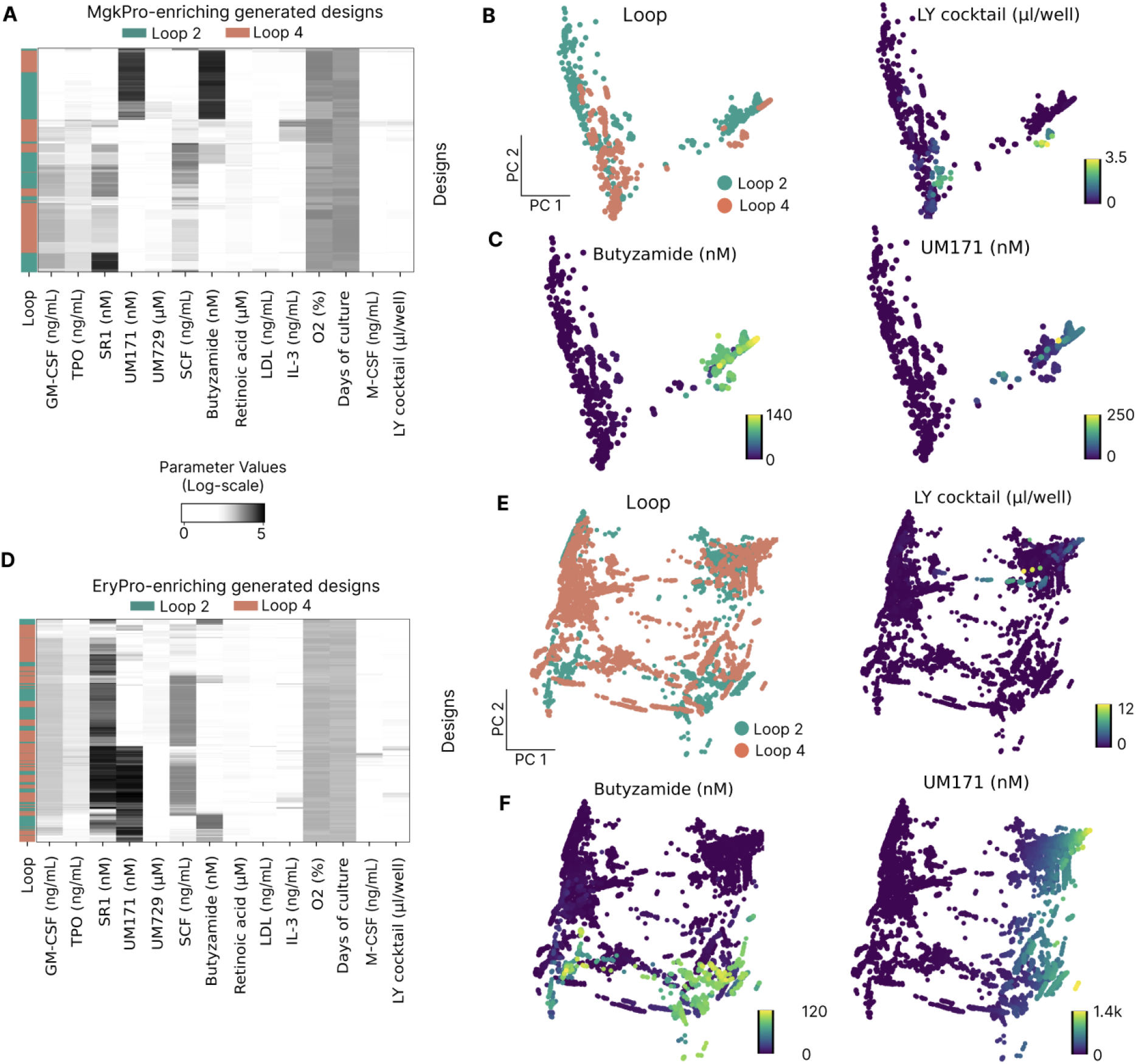
MgkPro- and EryPro-enriching designs generated in Loop 2 and Loop 4. **A:** Heatmap of design parameters for MgkPro-enriching designs generated in Loop 2 and Loop 4. **B:** PCA projection of MgkPro-enriching designs, colored by loop (left) and LY cocktail concentration (right). **C:** PCA projection of MgkPro-enriching designs, colored by butyzamide concentration (left) and UM171 concentration (right). **D:** Heatmap of design parameters for EryPro-enriching designs generated in Loop 2 and Loop 4. **E:** PCA projection of EryPro-enriching designs, colored by loop (left) and LY cocktail concentration (right). **F:** PCA projection of EryPro-enriching designs, colored by butyzamide concentration (left) and UM171 concentration (right).

**Supplementary Figure 12.**
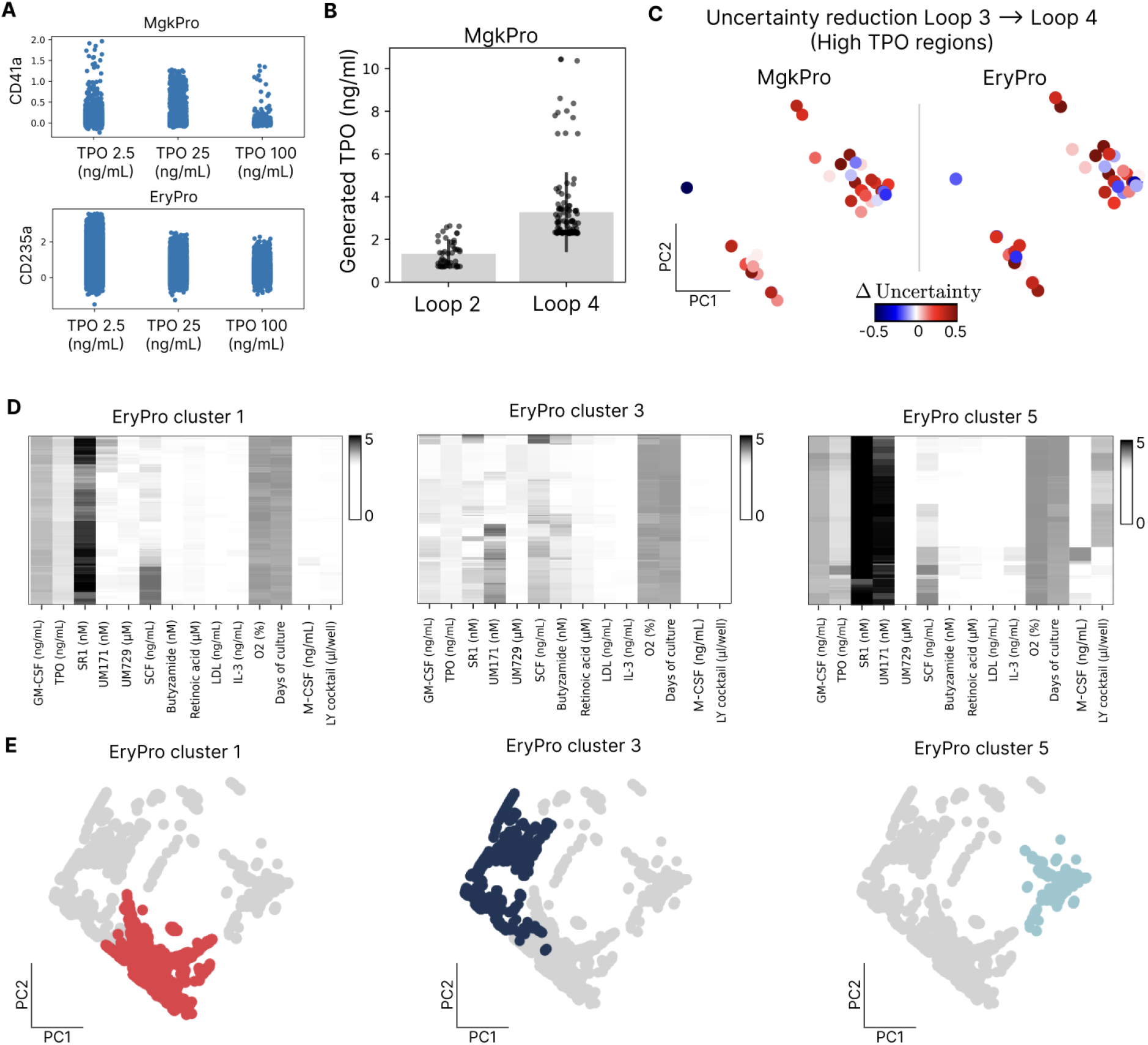
TPO expansion and uncertainty for MgkPro- and EryPro-enriching designs, and clustering of EryPro-enriching designs. **A:** CD41a expression across MgkPro-enriching designs (top) and CD235a expression across EryPro-enriching designs (bottom), at TPO concentrations of 2.5, 25, and 100 ng/mL. **B:** Generated TPO concentration for MgkPro-enriching designs in Loop 2 and Loop 4. **C:** PCA projection of MgkPro-enriching (left) and EryPro-enriching (right) designs in high-TPO regions, colored by change in uncertainty (Δ uncertainty). **D:** Heatmap of design parameters for EryPro-enriching designs in clusters 1, 3, and 5. **E:** PCA projection of EryPro-enriching designs, with cluster 1, cluster 3, and cluster 5 designs highlighted against all other designs (gray).

**Supplementary Figure 13.**
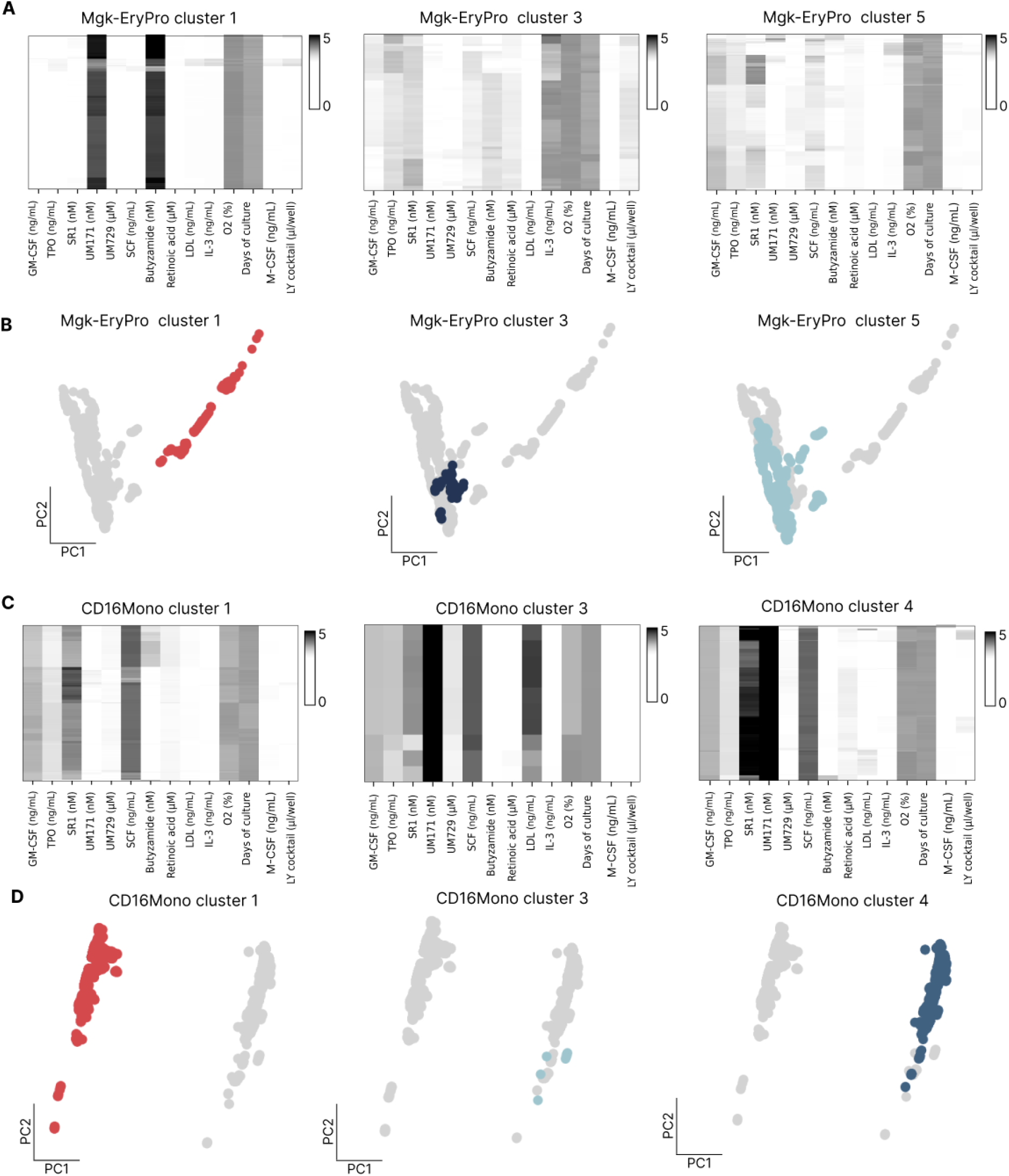
Clustering of MgkPro- and CD16Mono-enriching designs. **A:** Heatmap of design parameters for MgkPro-enriching designs in clusters 1, 3, and 5. **B:** PCA projection of MgkPro-enriching designs, with cluster 1, cluster 3, and cluster 5 designs highlighted against all other designs (gray). **C:** Heatmap of design parameters for CD16Mono-enriching designs in clusters 1, 3, and 4. **D:** PCA projection of CD16Mono-enriching designs, with cluster 1, cluster 3, and cluster 4 designs highlighted against all other designs (gray).

**Supplementary Figure 14.**
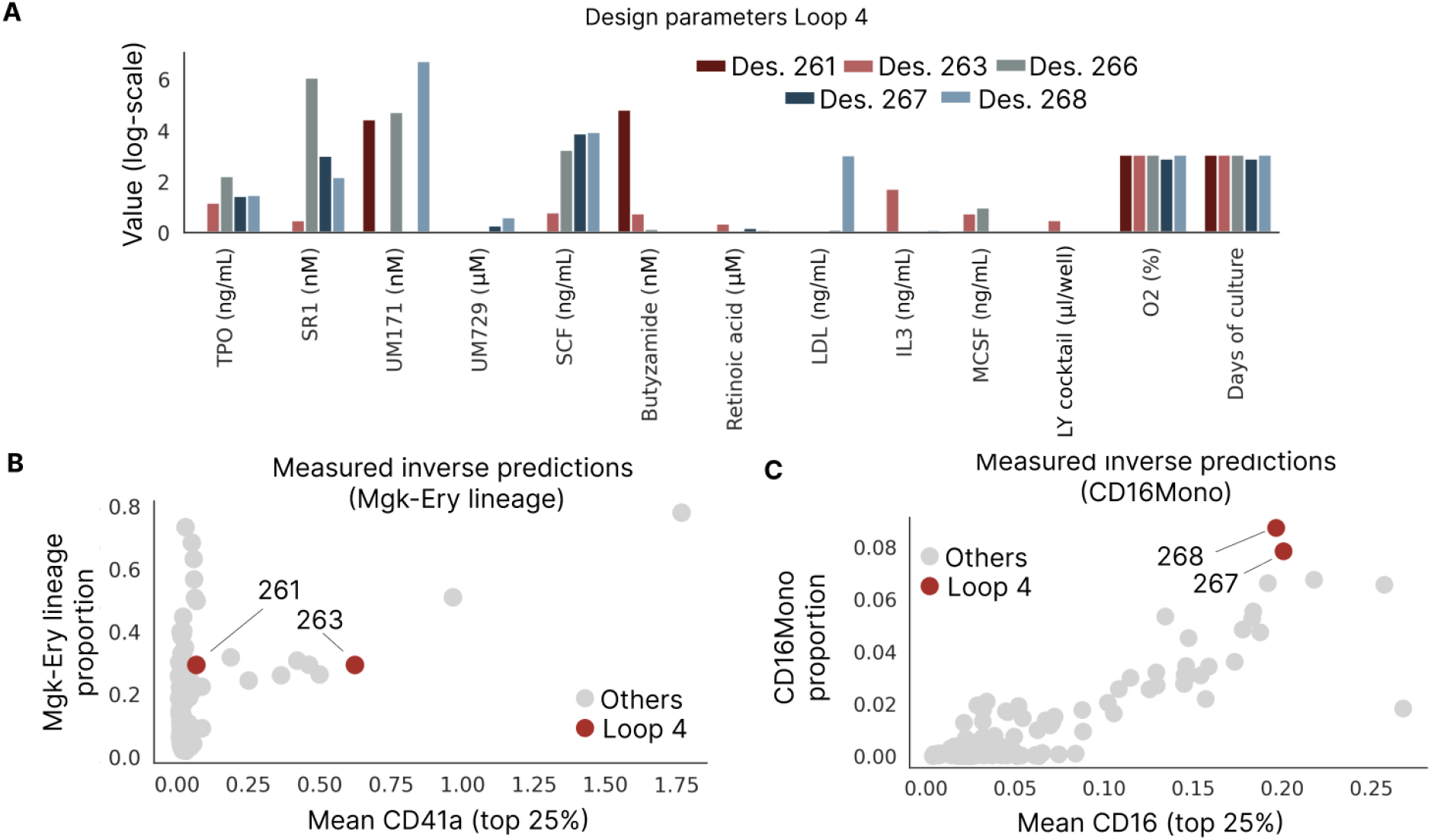
Design parameters and inverse-prediction validation for designs 261, 263, 266, 267, and 268. **A:** Log-scale concentration of each design parameter for designs 261, 263, 266, 267, and 268. **B**: Measured Mgk-Ery lineage proportion versus mean CD41a expression (top 25%) for designs 261 and 263 (Loop 4, red), compared with all other designs (gray). **C:** Measured CD16Mono proportion versus mean CD16 expression (top 25%) for designs 267 and 268 (Loop 4, red), compared with all other designs (gray).

**Supplementary Figure 15.**
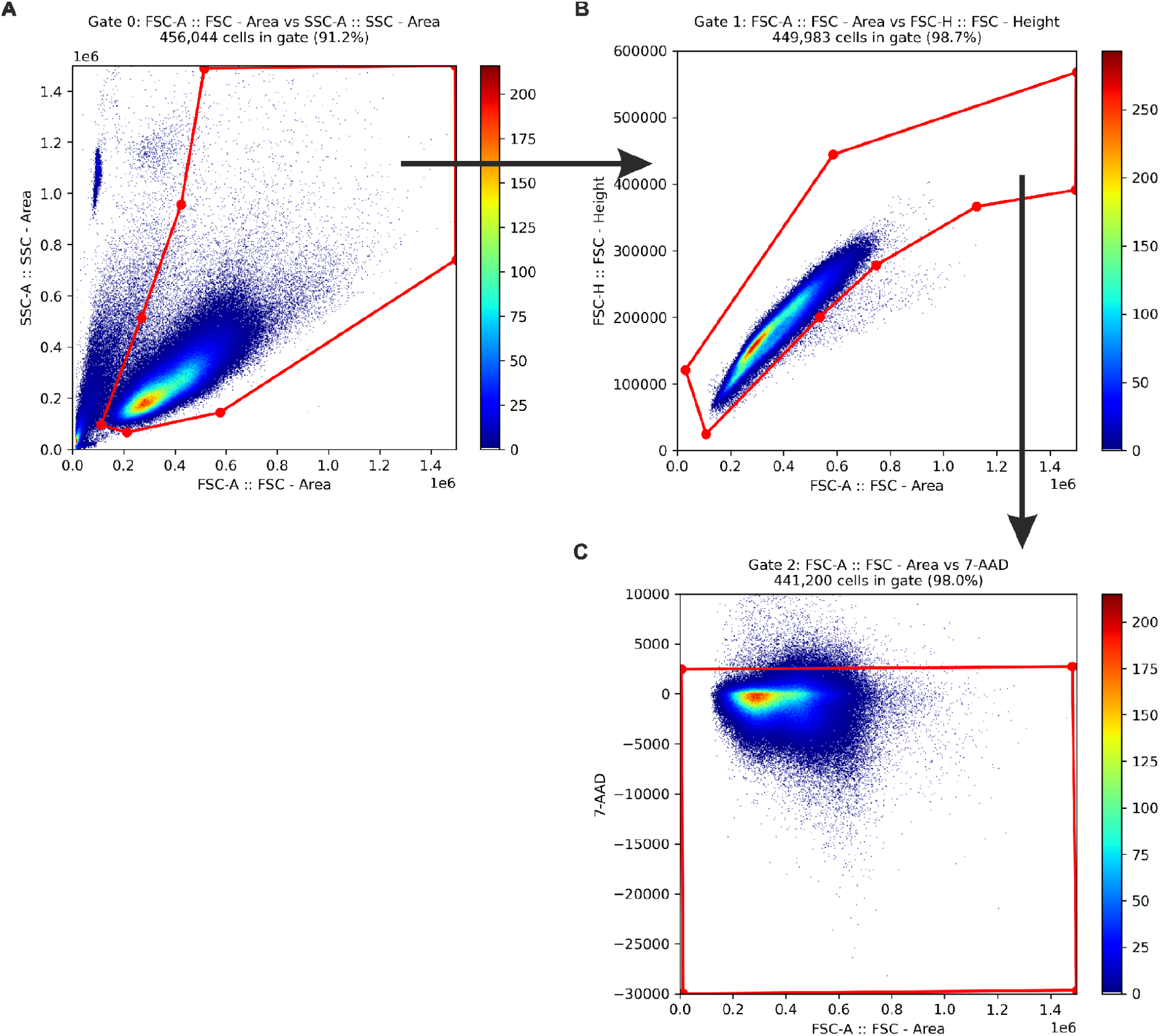
Gating strategy applied programmatically to all experiments. Gates were manually defined once on a representative sample and applied identically to every experiment via processing pipeline (as described in **Methods**), without per-sample adjustment. **A:** Cells were distinguished from debris using FSC-A and SSC-A. **B:** Single cells were selected from A using FSC-H versus FSC-A to exclude doublets. **C:** Live cells were selected from B as 7-AAD-negative events. Numbers indicate percentage of parent gate.

**Supplementary Figure 16.**
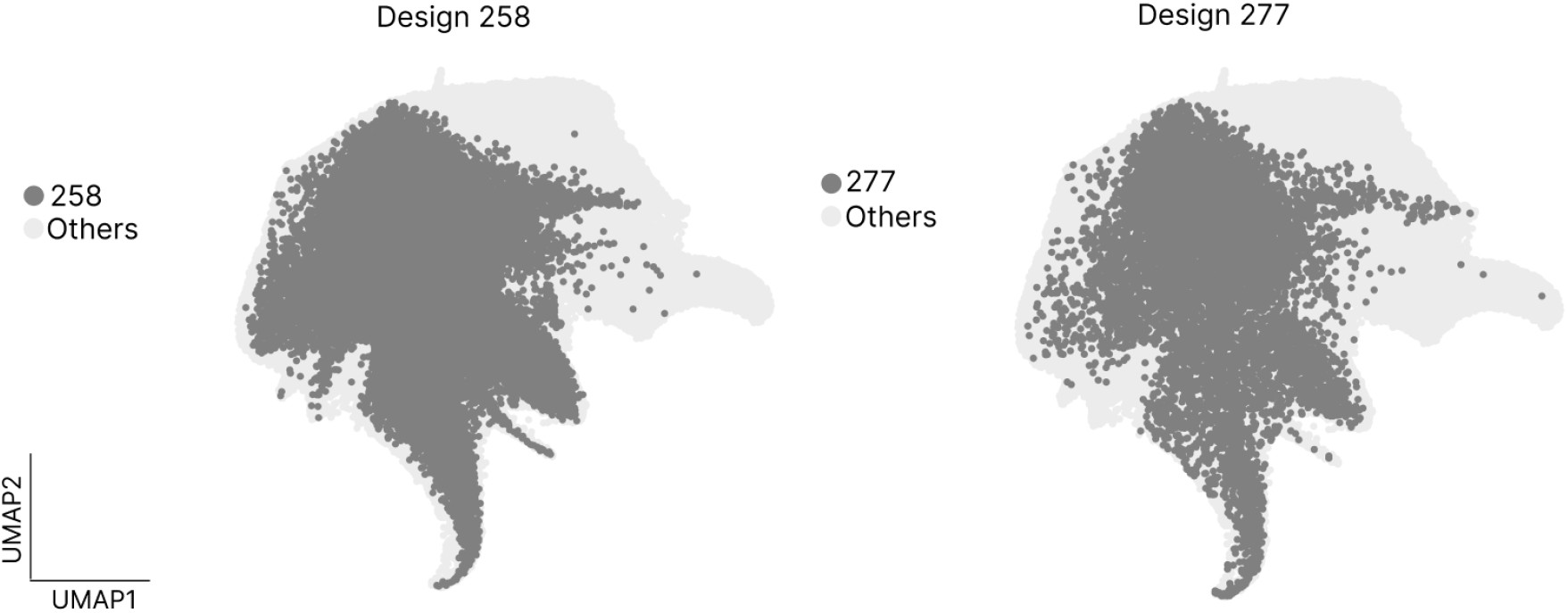
Batch consistency maintained through standardized spectral flow cytometry acquisition settings. Designs 258 and 277 were derived from the same donor (NCB040). Both designs produced highly similar hematopoietic landscapes despite having been analyzed in different experimental batches (Loop 3 and 4). UMAP visualization of the hematopoietic landscape, with cells from Design 258 (left) and Design 277 (right) highlighted in dark gray against all other cells (light gray, “Others”).

## Methods

### LabCompass

#### General description of LabCompass

LabCompass is a generative framework for designing experimental conditions that induce a user-defined target phenotype in single-cell data. We formulate this as an inverse modeling problem: a pretrained forward model predicts cellular responses to candidate experimental designs, and these predictions steer a generative model toward designs expected to induce the target phenotype.

At its core, LabCompass learns a Continuous Normalizing Flow (CNF) over the space of experimental designs using the Conditional Flow Matching (CFM) objective^27,29^. The experimental design space is represented continuously, for example as a high-dimensional vector encoding medium composition. The CNF models the prior distribution of feasible experimental conditions and is coupled to a differentiable forward model that predicts cellular responses conditioned on these variables. This enables efficient exploration and optimization of the experimental design space.

To generate conditions that induce the target phenotype, LabCompass uses loss guidance^62^ during the ODE (Ordinary Differential Equation) sampling process. At each generation step, we:

1. Predict the final, denoised experimental condition from the current noisy state.
2. Use the forward model to predict the cellular response under this condition.
3. Evaluate the predicted phenotype against the target using a differentiable loss.
4. Update the generative trajectory using an augmented vector field that combines the generative velocity with the negative loss gradient.

Importantly, all optimization occurs at inference time, without retraining the generative or forward models for different query optimization phenotypes. This allows the same model to efficiently optimize for different target cell states with no additional training overhead. In what follows, we introduce the spaces LabCompass operates on.

#### The experimental design space

Let *X* ⊆ℝ^*d*^ denote the design space: the set of experimental parameters used to culture cells, such as molecule concentrations, oxygen levels, or culture duration; experimental designs are assumed to be drawn from the *design prior distribution, p* ϵ *P*(*X*), used to unconditionally sample experimental designs *x ∼ p*(·). Given a target phenotype, LabCompass’s objective is to generate an optimal design *x*^⋆^·*X* expected to induce it.

#### The cellular state space

Cells are represented in an embedding space *ε* ⊆ ℝ^*g*^. Each design *x* maps, via the forward surrogate *F*, to a conditional density over cellular states, *p*(*e* | *x*) · *P*(*ε*), from which synthetic cellular profiles ê ∼ *p(*· |*x)*are generated. Identifying *x*^⋆^ therefore amounts to finding designs whose generated profiles ê are consistent with the desired phenotype.

To accommodate the stochasticity of the surrogate forward model *F*(*x*) in our notation, we rewrite it as taking an extra random argument *z* ∼ ζ (·), where ζ (·)^;^· *P* (*ε*)is the tractable latent distribution over the cellular expression space *ε* – we set ∼ (·) : = *N*(· |O_*ε*_ I_*ε*_). Hence, the forward model is defined as ê : = *F*(*x, z*) · *ε*.

#### Phenotypic readout

A phenotypic readout function *h* : *ε* → *y* extracts a biological property of interest, ŷ = *h* (ê), from a generated cellular profile ê — for example, cell-type proportions, marker expression statistics, or distributional distances to a reference population. LabCompass optimizes designs so that the resulting readout ŷ matches a user-specified target phenotype *y*^⋆^ ·*y*.

#### The probability model for *in vitro* hematopoiesis

The generated dataset *D* of *N*_*D*_ spectral flow cytometry events is defined as the collection of triplets 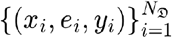 of experimental designs *x*_*i*_ ·*X*, cellular states *e*_*i*_ · *ε* and phenotypic readouts *y*_*i*_ *· y*; each triplet is assumed to be drawn from the joint probability density (*x*_*i*_,*e*_*i*_,*y*_*i*_) ∼ *p* (*x,e,y*) · *P (X* ×*ε*×*Y)*. We assume that the joint density factorizes as:

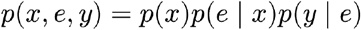

That is, we assume phenotypic readouts y·*Y* to be conditionally independent of designs *x* · *X*, given cellular states *e* ·×.

While we resorted to this probability model for the hematopoiesis experiments, it can be flexibly adapted to accommodate more complex scenarios; for example, one can model the system in different contexts *c* ×*C* by simply conditioning the experimental prior and the forward surrogate on them.

#### The likelihood over the phenotypic readouts and the posterior over designs

From the above probability model, it follows that the likelihood over phenotypic readouts *y* × *Y*, given experimental designs is *x* × *X*:

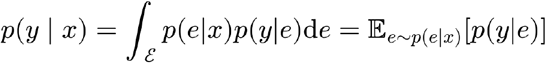

LabCompass aims to approximate the *posterior distribution* over the experimental designs *x* ×*X*, given a phenotypic readout *y* × *Y* – that is *p*(*x* | *y*); from Bayes’s theorem, we can write the posterior distribution as:

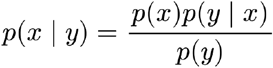

Where *p* (*y*) is the marginal density over the phenotypic readout, given by:

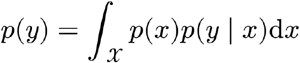

#### Learning the prior distribution with Conditional Flow Matching (CFM)

The design prior is learned using CFM. It can be expressed in terms of a time-dependent probability path {*p*_*t*_(·)}*t*ϵ[0,1], interpolating between a tractable noise distribution *q*_0_ (·) ϵ *P* (X)at time *t* = 0 and the design distribution *q*_1_ (·) ϵ *P* (X)at time *t* = 1. A time-dependent marginal velocity field*υ*_*t*_(·) is said to be generating the path {*p*_*t*_(·)}*t* ϵ[0,1] if *p*_*t*_ (·) and *υ*_*t*_ (·) satisfy the continuity equation *∂*_*tpt*_ *+* ∇ · (*p*_*t*_υ_t_) = 0. The integration of the Ordinary Differential Equation (ODE)

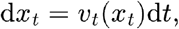

with *x*_0_ ∼ *q*_0_ (·) over the time interval *t* ϵ [0,1]yields samples from the design distribution *q*_1_ (·), which defines the learned design prior. We provide the necessary theoretical background on CNFs and CFM in **Theoretical Background**.

#### Inverse models and exponentially-tilted distributions

For potential function Ψ :*X* →ℝ, inverse modeling tasks can be generally interpreted as trying to approximate the exponentially-tilted distribution

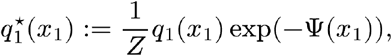

where *Z* is the intractable partition function, defined as:

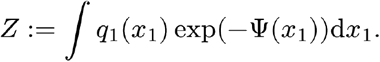

LabCompass uses the forward model *F* (·), the phenotypic readout mapping *h* (·), and the phenotypic loss *ψ (*·,·)to define the potential function Ψ (·). Specifically, the potential function is defined as

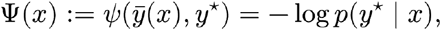

up to a constant independent of *x*, where 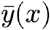 is the phenotypic response predicted by the forward model and *y* ^⋆^ is the target phenotype. The tilted distribution is therefore the posterior over experimental designs given the target phenotype,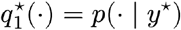.

#### One-step estimator of clean designs

To evaluate the potential function in a semantically meaningful region of the design space, intermediate points along sampling trajectories need to be mapped to their clean fully-generated version. This is achieved with an invertible one-step-estimator as 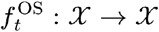 as:

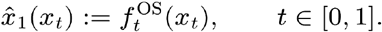

The simplest instantiation is the linear estimator, which extrapolates linearly along the prior velocity field to the endpoint of the time interval:

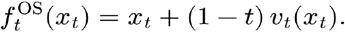

This step is crucial to stabilize the guidance signal, ultimately enhancing the quality of the posterior samples. We parametrize it in terms of a flow map model^63^, distilled from the prior velocity field, which recovers the linear estimator as a special case. We provide more details on flow map distillation in **Theoretical Background**.

#### Inference-time steering of prior generative trajectories

To achieve sampling from the exponentially-tilted distribution, LabCompass biases the generative field using the gradients of the target potential, evaluated on the estimated clean designs. We define the time-dependent intermediate potential function

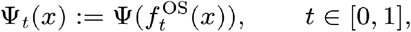

which is evaluated at intermediate points along sampling trajectories, through the one-step estimator. Denoting by *z*_*t*_ the guided trajectory, the steered sampling ODE reads

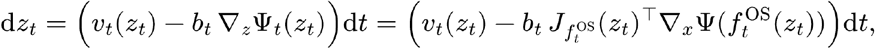

where the second equality follows from the chain rule and *b* : [0,1] ⟶ℝ_≥0_ is the guidance strength scheduler, instantiated depending on the definition of the prior probability path. We provide the derivation of the inverse ODE and of the guidance strength scheduler in **Theoretical Background**. The guidance term can be efficiently evaluated using automatic differentiation libraries.

Therefore, a single generative step at time *t* ϵ [0,1] and intermediate latent design *z*_*t*_ involves the following sequence of operations:

1. Map intermediate latent design *z*_*t*_ to estimated clean design 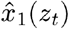 with the one-step-estimator as 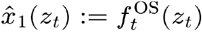.
2. Evaluate the potential function on the estimated clean design, 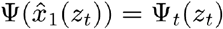.
3. Backpropagate the potential Ψ (·) through the one-step estimator 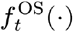 to obtain the gradients 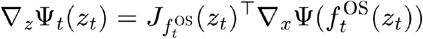.
4. Compute guidance strength scheduler *b*_*t*_ and update the velocity as *υ*_*t*_(*z*_*t*_) − *b*_*t*_ ∇*z* Ψ *t* (*z*_*t*_)

#### LabCompass as a preconditioned gradient flow on the potential landscape

It is possible to show that, when the one-step-estimator is given by the flow map, LabCompass’s inverse sampling dynamics coincides with a preconditioned gradient flow in data-space over the time interval *t* ϵ [0,1]:

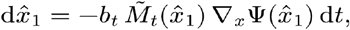

where 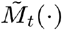 is a symmetric positive-semidefinite matrix that depends on the Jacobian of the flow map. Denoting by 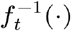 the inverse of the one-step estimator 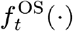, the preconditioning matrix is given by

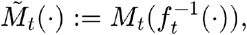

where *M*_t_ (·)is defined as the Jacobian outer product

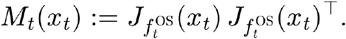

#### Approximating the phenotypic likelihood with the forward model

To approximate the phenotypic likelihood, we draw *M* ϵ ℕ noise samples 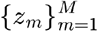, with *z*_*m*_ ∼ ζ (·), and compute the generated cellular profile ê_*m*_ = *F*(*x,z*_*m*_)for each noise sample *z*_*m*_, such that ê_*m*_ ∼ *p* (· ∣ *x*)for all *m*– this allows us to generate synthetic cellular profiles induced by experimental design x ϵ *X*. For each of the *M* samples, the phenotypic readout function *h* (·)maps generated cellular profiles ê to their estimated response ŷ_*m*_ :_*h*_(ê_*m*_)ϵ *y*. Combining all together, we can write ŷ (*x,z*_*m*_) : = *h* (*F*(*x,z*_*m*_)).

To integrate over the induced synthetic cellular profiles, the response is then computed by averaging the response vector across the *M* synthetic samples, that is:

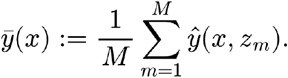

This Monte Carlo integration is justified by the definition of the probability model, and by the derivation of the phenotypic likelihood p(*y* ∣ *x*), an expectation under the conditional distribution over the cellular states. Here *y* is a group-level property of the population induced by design, rather than a property of an individual cell, and the *M* synthetic profiles 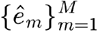 stand in for that population. This averaging form applies to readouts that decompose over cells, as is the case for the cell-type composition used here, where *h*(·) returns per-cell class probabilities and 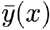 is the mean cell-type probability over the induced population. For readouts that do not decompose over cells, such as distributional distances to a reference population, the readout is instead evaluated on the full set 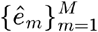.

In simple terms, the phenotypic likelihood given design *x* ϵ *X* is approximated with the following steps:

1. Sample *M* noise samples *z*_*m*_ ∼ ζ (·).
2. Estimate the synthetic cellular profiles with the forward model ê_*m*_ : = *F* (*x,z*_*m*_) for all *m*.
3. Compute the phenotypic response ŷ_*m*_ : = *h* (ê_*m*_) for all *m*.
4. Average the phenotypic response ŷ_*m*_ over the *M* observations, to obtain 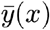.

#### Cell-type composition optimization for *in vitro* hematopoiesis

The inverse experiments for cell-type composition optimization aim to find experimental designs *x* ϵ *X* such that the induced cellular populations will exhibit a user-defined composition in terms of cell types, denoted by the target response *y*^⋆^ ϵ *y*. In this setting, the response space is defined as the *K*-dimensional probability simplex, that is *y* : = Δ^*K*^; the average response of the system 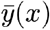 models instead the mean cell type probability over the induced population.

The mean cell type probabilities are then compared to the target composition *y*^⋆^ using the cross-entropy loss:

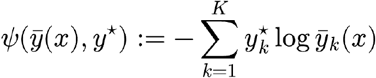

and the potential for inverse sampling is therefore defined as:

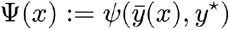

When the goal is to induce a pure population of the target cell type *k*^⋆^ ϵ {1,…,*K*}, the target composition is the one-hot vector with 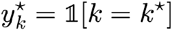, and the potential reduces to 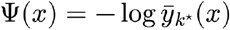 – it will favor conditions that maximize the proportion of the target cell type. This dramatically simplifies the potential landscape, as optimizing across multiple target cell types might involve antagonistic interactions that could increase the difficulty of the optimization task.

Theoretically, the cross-entropy objective is closely related to the entropy of the target cell-type composition *y*^⋆^ ϵ *y*. The minimum attainable value of the cross-entropy loss coincides with the entropy of the target cell-type composition. This can be easily seen by plugging *y*^⋆^ as the predictor, hence obtaining:

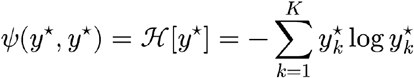

Where is the entropy of the target distribution. Target compositions involving diverse populations will be associated with a higher loss value and will be harder to optimize.

#### Forward model’s uncertainty quantification

The stochasticity of the forward model allows LabCompass to quantify the uncertainty of the estimated response 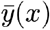. Let *N*_pop_ ϵ ℕdenote the number of generated populations used to estimate the forward model uncertainty. For each of the *N*_pop_ populations, we sample *M* latent codes *z*_*m*_,_*n*_∼ζ (·) – we use these to generate synthetic cellular profiles ê_*m*_,_*n*_ = *F* (*x,z*_m_,_*n*_) and estimate the corresponding phenotypic readout map. For each population, we compute the response as:

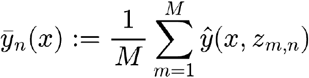

This yields *N*_pop_ estimated cell-type compositions 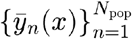 for design *x* ϵ *X*: the variability over the *N*_pop_ populations serves as an uncertainty measure. In this scenario, one can for example assess the variability of the loss function 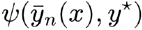, or that of any other quantity computed from 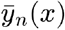. More rigorously, uncertainty quantification amounts to the definition of a function *u* : *X* → ℝ _≥ 0_that maps experimental designs *x* to the uncertainty around the estimated induced response. For both cases just presented, this can be expressed by considering the uncertainty readout function ω : *y* → ℝ – mapping the estimated phenotypic response 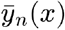 to a scalar whose variability across the *N*_pop_ populations will quantify the response uncertainty. Let

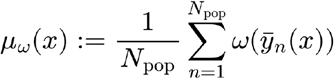

denote the empirical mean over the generated populations; then we define

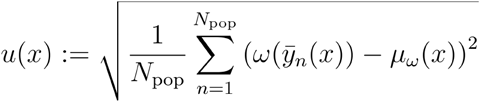

as the uncertainty measure.

To quantify the uncertainty of the enrichment of a target cell type *k* ^⋆^ ϵ {1,…,*K*}for phenotypic response *y*, the uncertainty readout map is 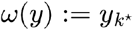 for the uncertainty of the loss function, ω(·) is instead defined as the loss function itself, that is ω (*y*) : = *ψ* (*y, y*^⋆^) (**Supplementary Tab. 1**). We consistently observed that, for the cell-type proportion optimization task, the latter provides a more reliable uncertainty measure, while the former did not exhibit consistent patterns.

**Supplementary table 1:** Definition of the uncertainty readout mapping for the two cases of loss function and target cell type uncertainty.

|  |  |
| --- | --- |
| | $\omega(y)$ |
| Cell-type composition optimization | $y_{k^*}$ for target cell type $k^* \in \{1, \dots, K\}$ |
| Loss function | $\psi(y, y^*)$ |

### Theoretical Background

#### Continuous Normalizing Flows: dynamical approaches to generative models

CNFs are a class of generative models that aim to approximate an intractable target data distribution *q*_1_ (·)ϵ *P* (*X*) by defining a time-dependent marginal probability path {*p*_*t*_(·)}_*t*_ϵ[0,1], interpolating between a tractable noise distribution *q*_0_(·) ϵ *P*(*X*) at time *t* = 0 and the data distribution *q*_0_(·) ϵ *P*(*X*) at time *t* = 1 A time-dependent marginal velocity field *υ*_*t*_ (·) is said to be generating the path {*p*_*t*_(·)}_*t*ϵ[0,1]_ if *p*_*t*_ (·) and *υ*_*t*_ (·)satisfy the continuity equation ∂_*t*_*p*_*t*_ + ∇ ·(*p*_*t*_*υ*_*t*_) = 0. The integration of the Ordinary Differential Equation (ODE)

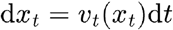

over the time interval *t* ϵ [*t*_0_,*t*_1_]is the flow map diffeomorphism from time *t* = *t*_0_ to time *t* = *t*_1_, that is:

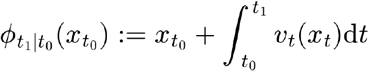

For 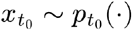, it follows that 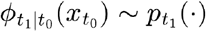. Sampling is then obtained by drawing latent noise from the tractable noise *x*_0_ ∼ *q*_0_ (·)distribution, and computing the flow map *ϕ*_1|0_from time *t*_0_ = 0 to time *t*_1_ = 1, which requires solving an Initial Value Problem (IVP) on the generative dynamical system.

According to our notation, we will have that 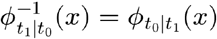 for all (*t*_0_,*t*_1_) ϵ [0,1]^2^ and *x* ϵ*X*. The flow map therefore satisfies the consistency conditions 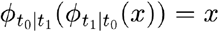 for all (*t*_0_,*t*_1_) ϵ [0,1]^2^and *x* ϵ*X*. Being a diffeomorphism, the flow map 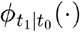 can be intended as a change of variable, transforming the density 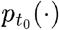 at time *t* = *t*_0_ into the density 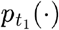 at time *t* = *t*_1_; this relation is characterized by the push-forward equation, reading:

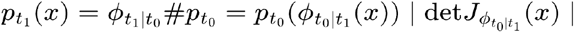

#### Conditional Flow-Matching: simulation-free training of CNFs

To learn the generative ODE, Flow Matching regresses a neural velocity field 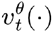 against the marginal velocity field *υ*_*t*_(·) using the objective function:

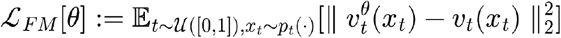

However, this naive formulation requires sampling from the intractable marginal probability path *p*_*t*_(·) and access to the unknown marginal velocity field *υ*_*t*_(·).

Conditional Flow-Matching (CFM) overcomes these issues by introducing a tractable conditional probability path {*p*_*t*|1_ (·|*x*_1_)}_*t*∈[0.1]_ conditioned on data points to obtain the marginal probability path with the marginalization:

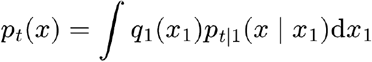

The conditional probability path {*p*_*t*|1_ (·|*x*_1_)}_*t*∈[0.1]_ is itself generated by a tractable conditional velocity field *υ*_*t*|1_ (·|*x*_1_)} – together, they satisfy the continuity equation *∂*_*t*_*p*_*t*|1_ + ∇ · *(p*_*t*|1_*υ*_*t*|1_) = 0. The conditional probability path is commonly instantiated as 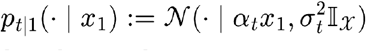, where *α*_*t*_ : [0,1] → [0,1] is the interpolation schedule and *σ*_*t*_ : [0,1] → [*σ*_*min*_, 1] is the noise schedule, with [*σ*_*min*_ ∈ ℝ ≥0; the pair {(*α*_*t*_, *σ*_*t*_*)}*_*t*∈[0.1]_ is referred to as the schedule. The schedule is designed to satisfy the boundary conditions *α*_*t*_ |_t=0_ = 0, *α*_*t*_ |_t=1_ = 1 and *σ*_*t*_ |_t=0_ = 1, *σ*_*t*_ |_t=1_ = *σ*_*min*_ 27. In this case, the tractable conditional velocity field is given by:

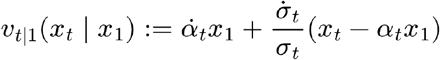

where 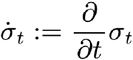 and 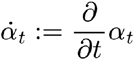. The marginal velocity field can be expressed in terms of the conditional velocity field as:

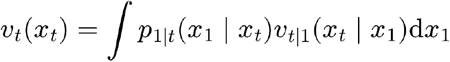

where 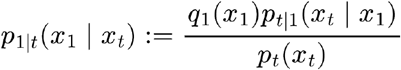 is the posterior on data-points *x*_1_, given intermediate latent codes *x*_*t*_. It is then possible to show that, up to a constant term independent of *θ*, the tractable objective function

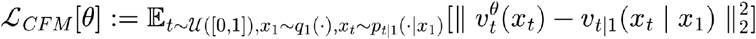

shares the same gradients as *L*_*FM*_, that is ∇_*θ*_*L*_*CFM*_ = ∇_*θ*_*L*_*FM*_. Hence, it is possible to learn the marginal velocity field *υ*_*t*_(·) *L*_*CFM*_ by minimizing instead of the intractable *L*_*FM*_, providing an avenue toward simulation-free training of CNFs.

#### Velocity, Score and Denoiser fields: a unifying view on Conditional Flow Matching

Under the assumption of the conditional probability path being given by 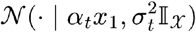, we can rewrite the marginal velocity field as:

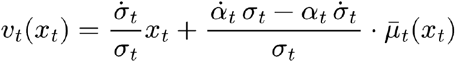

where 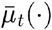 is the denoiser field, defined as 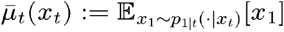. From the definition of the marginal probability path, it is possible to show that the marginal score field ∇ *x*_*t*_ log *p*_*t*_(*x*_*t*_) can be expressed in terms of the tractable conditional score field ∇ *x*_*t*_ log *p*_*t*|1_ (*x*_*t*_|*x*_1_) as:

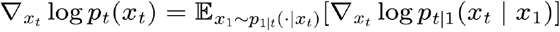

From Gaussianity of the conditional probability path, it follows that:

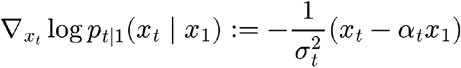

It is then possible to express the marginal score field in terms of the denoiser field as:

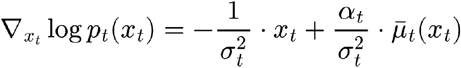

By simple algebraic manipulations, this makes it possible to relate the velocity field to the score field as:

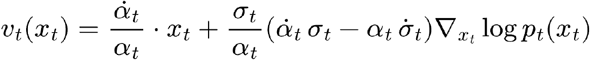

Defining 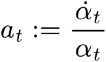 and 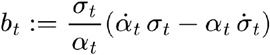, we can write:

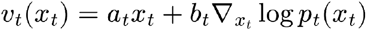

This relationship is crucial for the definition of the augmented dynamics used by the inverse sampler. For the schedule *α*_*t*_ := *t* and *σ*_*t*_ := (1−*t*), the rescaling coefficients will be given by 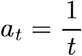 and 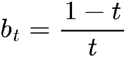. The *b*_*t*_ coefficient will coincide with the guidance strength scheduler for inverse sampling.

#### Flow Maps: distilling a one-step-estimator from CFM velocity field

While CFM models enable efficient, simulation-free training of normalizing flows, their inference procedure is expensive as it requires the simulation of continuous-time dynamics - this is done by stacking several forward passes on the neural velocity field, significantly increasing the computational overhead. This bottleneck has justified several works on consistency models, attempting to reduce inference complexity by one-step or few-steps estimators^*63,64*^.

Generally, the approximation target of consistency models is the integral solution of the flow map *ϕ*_*t*|*s*_*(·)*, instead of its velocity. *Lagrangian map distillation learns a neural flow map* 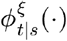 with neural parameters ξ ∈ Ξ, from a pre-trained neural velocity field 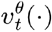, by minimizing the objective function:

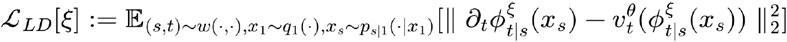

Where *w(*·, ·) ∈ *P* ([0,1]^2^)is the joint distribution over time indices (*s, t) ∈*[0,1]^2^. The neural flow map is designed to satisfy the condition 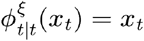. LabCompass utilizes neural flow maps as one-step estimators to map intermediate latent states *x*_*t*_ to estimated clean data points 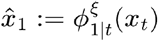, then used to evaluate the potential function Ψ(·). This step is necessary to ensure that the surrogate forward model *F*(·) is evaluated on well-supported regions of the design space where its estimations are accurate - crucial requirement for a stable and reliable optimization signal. In what follows we assume, as is standard for distilled maps, that the learned one-step estimator inherits the invertibility of the exact flow map it approximates.

#### Inverse sampling by inference-time steering of generative models

For potential function Ψ : *X* → ∝, inverse modeling tasks can be generally interpreted as trying to approximate the exponentially-tilted distribution

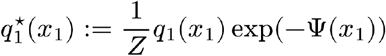

Where

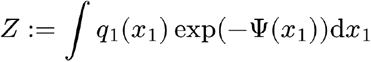

is the intractable partition function. As universal density approximators, continuous normalizing flows are often used as expressive priors to solve this task – by either changing the sampling ODE during generation (inference-time steering), or updating the prior weights (fine-tuning), it is possible to turn a CNF prior into an inverse sampler. LabCompass approaches inverse sampling with inference-time steering, using the forward surrogate model *F*(·), the phenotypic readout mapping *h*(·) and the phenotypic loss *ψ*(·) to define the potential function Ψ(·). We provide here a rigorous mathematical description of the steering mechanism.

To obtain a general formulation of the inverse dynamics, we need to derive the inverse marginal probability path 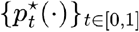 that samples from the exponentially-tilted distribution 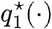; by substituting 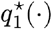 in place of *q*_1_(·) in the definition of the marginal probability path, we obtain:

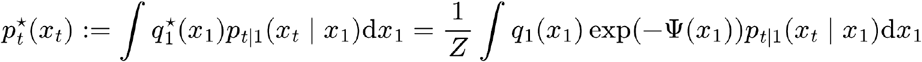

After replacing *q*_1_(*x*_1_*)p*_*t*|1_ (*x*_*t*_ | *x*_1_) *p*_*t*_ (*x*_*t*_*) p*_1_(*x*_1_ | *x*_*t*_), it follows that:

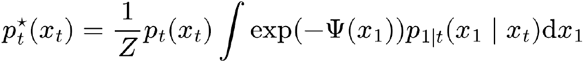

Defining the time-dependent intermediate potential function:

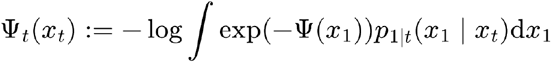

we can finally rewrite:

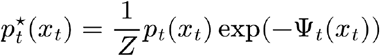

From the relationship between the velocity and the score field, it follows that the inverse velocity field 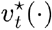 generating the probability path 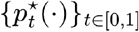 is given by 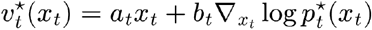; substituting 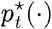 with its exponentially-tilted expression, we have:

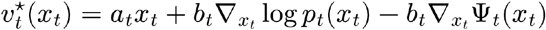

From the definition of the prior velocity field, we can finally derive the expression for the inverse velocity field as 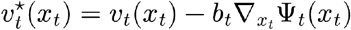. Hence, inverse sampling can be achieved by integrating the ODE:

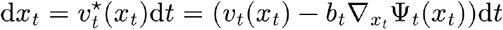

As the prior velocity *υ*_*t*_*(·)* is assumed to be learned upfront by the prior neural velocity field 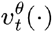, the evaluation of the inverse velocity 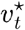 field hinges on the possibility to compute the gradients of the intermediate potential function 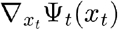. As we have shown, Ψ_*t*_ (*x*_*t*_) is defined as the logarithm of an expectation under the intractable posterior distribution *p*_*t*|1_ (·| *x*_*t*_). In practice, this quantity is approximated by evaluating the potential at the flow-map estimate of the clean data point, that is 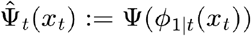, thereby exchanging the expectation of the exponential for the exponential evaluated at a single representative point. This corresponds to replacing the posterior *p*_*t*|1_ ·| *x*_*t*_) with the Dirac measure *δ*(*x*_*t*_ *− ϕ*_*1|t*_(*x*_*t*_)); the approximation is accurate when the posterior is sharply concentrated around *ϕ*_*1|t*_(*x*_*t*_), which holds increasingly well as *t* → 1, and it is the choice that makes the guidance term tractable with a single evaluation of the one-step estimator. From this approximation, it follows that 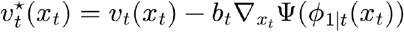. Using the chain rule, we have that 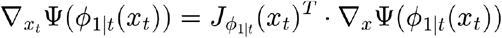, where 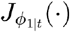 is the Jacobian of the flow map. Plugging back in the inverse velocity field, it follows that:

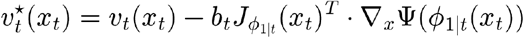

which can be efficiently evaluated using automatic differentiation libraries. The gradients of the potential function will be rescaled using the coefficient. As derived before, for the schedule *α*_*t*_ := *t* and *α*_*t*_ := (1 − *t*), the *b*_*t*_ coefficient exhibits a singularity at *t* = 0, diverging towards infinity; to numerically implement the sampling dynamics, one needs to clip its value to a maximum term, hence yielding the capped coefficient 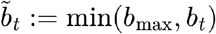, for positive scalar *b*_max_ ∈ ∝ _>0_.

#### The optimization dynamics in data space

Because the flow map is a diffeomorphism, *ϕ*_*1|t*_*(·)* represents the change of variable that transforms intermediate latent states into clean data points 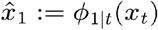. We can use this interpretation to derive the optimization dynamics in the data space from the sampling dynamics in the intermediate latent space. The total differential of 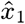 with respect to sampling time *t* ∈ [0,1] is given by 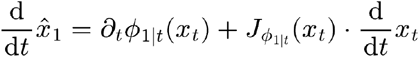, where 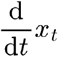 is given by the inverse velocity field 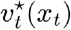; substituting with its expression, we obtain:

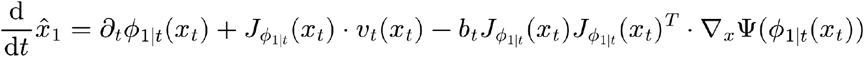

The prior flow map total differential term 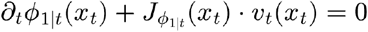 vanishes because *ϕ*_*1|t*_*(·)* is the solution operator of the prior ODE: along a prior trajectory the estimated clean point *ϕ*_*1|t*_*(x*_*t*_) is constant in *t*, so its total derivative is zero. Therefore, we remain with:

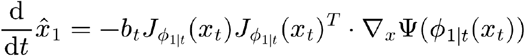

By defining 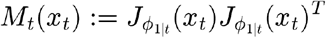, it is possible to rewrite the optimization dynamics as 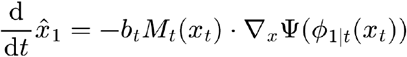. From the definition of the flow map, we have that 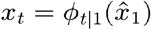; to express the dynamics in terms of clean points 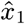, one can rewrite:

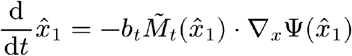

with 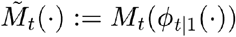 – the generative optimization dynamics act as a gradient flow, preconditioned on the flow map’s Jacobian outer product.

### Comparison with uniform random search

#### Defining the problem settings

For the toy comparison between LabCompass and the random search algorithm, we set the design space to the hypercube *x* := [*x*_min_, *x*_max_]^*D*^, where *D* ∈ ℕ represents the dimensionality of the design space. The hypercube was discretized with resolution *R* ∈ ℕ, *R* > 1 to create the grid 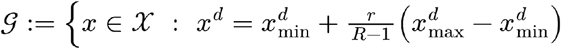, *d ∈ {1, …, D}, r ∈ {0, …, R −* 1}.

Experiments were carried out across resolutions *R* ∈ ℛ :{5,10,100} and dimensionalities *D* ϵ *D* : = {2,5,10,20,30}; in all cases, we set and. The cardinality of the search space is determined by the dimensionality and the resolution ; in particular, we have ∣ *G* ∣ = *R*^*D*^. The highest-dimensional tested combination was *D* = 30 with *R* = 100, implying a grid of around 10^60^ points (see **Supplementary Tab. 2** for the tested simulation grid).

**Supplementary table 2:** The dimensionalities and resolutions that were used to define the grid for the comparison with random search.

|  |  |
| --- | --- |
| **Dimensionality $D$** | 2, 5, 10, 20, 30 |
| **Grid resolution $R$** | 5, 10, 100 |

#### The data splitting regime for the toy comparison

We then sampled a proportion c ϵ [0,1] of the total number of points in the grid, capped at a maximum of 200,000 points. More specifically, we uniformly sampled without replacement from the total grid *G* to obtain an observed subgrid *G*_*observed*_ ⊂ *G* such that ∣ *G*_*observed*_∣ = min(200,000, *cR*^*D*^). The observed subgrid was further partitioned into disjoint training *G*_*train*_, validation *G*_*val*_, and test *G*_*test*_ subgrids, such that *G*_*observed*_ = *G*_*train*_ ∐ *G*_*val*_ ∐ *G*_*test*_. The training grid was therefore used to fit the prior and surrogate forward models, while the validation grid was used for model selection; target designs for the inverse tasks were sampled uniformly from the test subgrid *G*_*test*_. We reserved 70% of observed points for training, 20% for validation, and 10% for testing. For a target design *x*^⋆^ ϵ *G*_*test*_, drawn randomly from the test subgrid, the response was set to the squared Euclidean distance from *x*^⋆^, that is 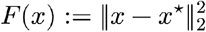; the objective function was instead defined as the identity function on the response,*ψ*(*y*) : = *y*, so that the optimum coincided with *x*^⋆^. A problem is then fully determined by the triplet (*D, R, x*^⋆^).

#### A theoretical analysis on the hit probability

The theoretical analysis on the hit-probability naturally derives from this setting. As seen, the total number of points in a *D*-dimensional grid of resolution *R* is. *R*^*D*^ Under a uniform distribution over the grid points, the probability of sampling *x*^⋆^ is 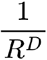. Hence, when executing uniform random search with replacement over a budget B ϵ ℕand *N* ϵ ℕindependent repeats, the probability that no evaluation hits the target is given by 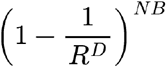. Thus, the probability that at least one of the *NB* trials hits the target *x*^⋆^ is given by 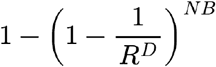, which decreases polynomially with resolution *R* and exponentially with the dimensionality *D*. For each dimensionality *D* ϵ*D* and resolution *R*ϵ*R*, we selected *Q* ϵ ℕoptima 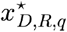, with *Q* = 10, yielding the suite of problems 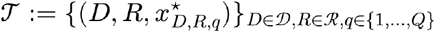.

### The estimation of the prior density over the design space

The prior CFM model was trained to transform an isotropic Gaussian distribution into the data distribution over the grid points. For this experiment, we chose the conditional probability path 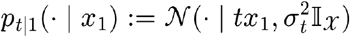, with σ*t* = 1 − *t* + *t*σ, where σ is the prior noise level. Unlike the schedules considered in **Theoretical Background**, here σ is allowed to exceed 1, so that σ*t* increases over the time interval; this was chosen deliberately to smooth the prior approximation in higher dimensions. The neural velocity field 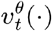was instantiated using a state encoder with one hidden layer of 128 neurons, using Rectified Linear Unit (ReLU) activation function. The state encoder *f*_*state*_ (·) mapped input states *x* to a 32-dimensional latent vector *z* : = *f*_*state*_(*x*). The latent state representation was then concatenated with the continuous time index, to create the concatenated vector *z*_*concat*_ : = [*z,t*]. The concatenated vector was *z*_*concat*_ then fed to the velocity decoder *f*_*velocity*_ *(·)*, to obtain the final velocity field 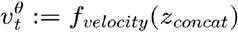. The CFM prior was trained for 100,000 gradient steps, using a training batch size of 512. The prior velocity’s parameters were updated using the AdamW optimizer with learning rate of 0.0005 and default weight decay of 0.01. The architecture for the Flow Map prior was analogous to the CFM one, with the only difference that the concatenated representation contains two time indices, that is *z*_*concat*_ : = [*z,t,s*]. The flow map prior was trained for 150,000 gradient steps, using a training batch size of 512. The flow map’s parameters were updated using the AdamW optimizer with learning rate of 0.0005 and default weight decay of 0.01.

### Potential approximation: surrogate and ground-truth optimization

We ran the comparison in two different settings: either by requiring access to the ground-truth forward model *F*(*x*) (ground-truth optimization), or by using a surrogate model to approximate it (surrogate optimization). Ground-truth optimization assumes a closed-form expression for the forward model; conversely, surrogate optimization learns an approximation 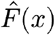 of the forward model from the training dataset 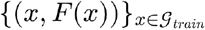. We considered both a linear surrogate model, and a multi-layer perceptron (MLP) with two hidden layers of 32 neurons each and Exponential Linear Unit (ELU) activation. All the surrogate models were trained for 1,000 epochs with a training batch size of 512, using the Mean Squared Error (MSE) loss; to avoid overfitting, early stopping with patience of 100 epochs was enforced. To update the surrogate parameters, we used the Adam optimizer – we swept the optimizer’s hyperparameters over different values for the learning rate and the weight decay. All the used training parameters are presented in **Supplementary Tab. 3**.

**Supplementary table 3:**
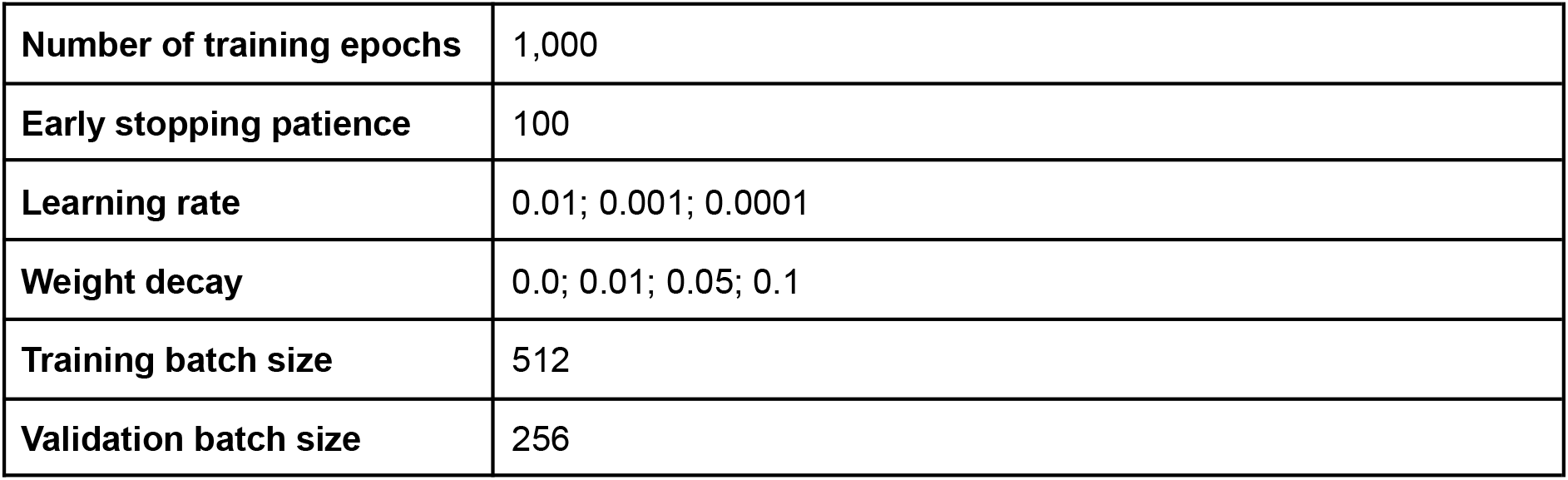
The configurations used to train the surrogate potential for the comparison with random search.

|  |  |
| --- | --- |
| **Number of training epochs** | 1,000 |
| **Early stopping patience** | 100 |
| **Learning rate** | 0.01; 0.001; 0.0001 |
| **Weight decay** | 0.0; 0.01; 0.05; 0.1 |
| **Training batch size** | 512 |
| **Validation batch size** | 256 |

#### The definition of the optimization problems and budget

LabCompass sampling ODE was simulated with Euler discretization – ensuring a single function evaluation per integration step – therefore enabling a fair comparison (**Supplementary Tab. 4**). The capping value for the coefficient 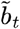 that rescales the potential gradients was swept over *b*_max_ ϵ {0.5, 2.0, 5.0}. The number of discretization steps was set to the reference budget *B*_max_ ϵ ℕ, while the number of generated optimization samples corresponded to the number of independent restarts *N* = 100 for uniform random search. The baseline was executed with increasing budget, until the reference budget *B*_max_ was reached; to determine the budgets to run uniform random search for, a minimum budget *B*_min_ ϵℕ, with *B*_min_, < *B*_max_ was progressively scaled according to a multiplier *m* ϵ ℕ, creating the sequence of evaluation budgets {*m*^k^ *B* _min_}; _*k* ϵℕ_the multiplier was set to m := min (5.0, 2.0 + 0.2ln *D*), while the minimum budget was *B*_min_ = 1.

We determined the reference budget *B*_max_ and the prior noise level σ according to the dimensionality of the problem. While the former was set to achieve a fair comparison with random search, the latter was chosen to increase the level of smoothing of the prior approximation with higher dimensionality; this was believed to reduce the effect of overfitting with higher dimensions by favoring a more robust approximation.

**Supplementary table 4:** The dimensionality-dependent reference budget and prior noise levels for LabCompass in the comparison.

| | $D = 2$ | $D = 5$ | $D = 10$ | $D = 20$ | $D = 30$ |
| --- | --- | --- | --- | --- | --- |
| **Reference Budget**<br>$B_{\max}$ | 10 | 100 | 1,000 | 10,000 | 10,000 |
| **Prior noise level** $\sigma$ | 0.1 | 0.2 | 0.5 | 1.0 | 2.0 |

#### Baseline: uniform random search over the discretized grid

The baseline for this comparison was defined as a uniform distribution over the experimental grid *G*, executed with budget *B* ϵ ℕ over *N* ϵ ℕindependent optimization restarts. For optimization restart *n* ϵ {1,…,*N*}, the uniform baseline samples the set of designs 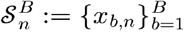, with *x*_*b,n*_ ∼ *U*(*g*) for all *b* and *n*. The designs are used to evaluate the potential function Ψ(·), and are then ranked based on the resulting potential values to obtain:

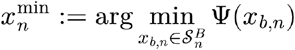

The *N* independent restarts yield the set of candidate solutions 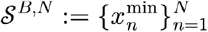. To ensure fair comparison between uniform random search and LabCompass, we considered the budget *B* to be analogous to the number of Euler discretization steps in the ODE integration for the inverse dynamics; the number of independent restarts *N* was then configured as the number of independent samples generated by LabCompass at each optimization round.

### Biological experiments

#### Human umbilical cord blood processing and CD34+ enrichment

CD34+ cells were isolated from human umbilical cord blood (hUCB), obtained from the Nolan Foundation (REC ref. 20/EM/0028) or the Cambridge Blood and Stem Cell Bank (REC ref. 18/EE/0199). hUCB Samples were processed as previously described^65^. Briefly, hUCB-derived mononuclear cells were isolated using density gradient centrifugation. Whole blood samples were first diluted 1:1 in Dulbecco’s PBS (DPBS; Sigma-Aldrich, D8537), then separated using Pancoll (PAN-Biotech, P04-601000) at 500 rcf for 25 minutes with brake off. Erythrocytes were lysed by incubation with ammonium chloride-based Red Blood Cell Lysis Buffer (BioLegend, 420302) for 15 minutes at 4°C in the dark. CD34+ selection was then performed using CD34 MicroBeads and FcR blocking reagent (Miltenyi Biotec, 130-100-453) in DPBS containing 3% fetal calf serum (FCS; PAN-Biotech, P40-37500) for 30 minutes at 4°C, followed by separation using LS Columns (Miltenyi Biotec, 130-042-401). Purified CD34+ cells were cryopreserved in FCS containing 10% DMSO (Sigma, D2650) and stored at -150 °C until use.

#### Culture condition screening

CD34+ cells were seeded at 10,000 cells per well in a 96-well CellBIND tissue culture plate (Fisher Scientific, 10510733). Cells were cultured at 37°C with 5% CO_2_ and 20% O_2_, unless stated otherwise, in base media and to which the different molecules were added. Base media was composed of: IMDM medium (Gibco, 12440-053), 1% (v/v) Insulin-Transferrin-Selenium-Ethanolamine (ITS-X) (Gibco, 51500-056), 1% (v/v) Penicillin-Streptomycin-Glutamine (P/S/G) (Gibco, 10378-016), 0.1% Soluplus (BASF, 50539897), 5 μM 740Y-P (Selleckchem, S7865), 100 μM α-Monothioglycerol (MTG) (Sigma, M1753) and 10 ng/mL rhFlt3L (Miltenyi, 130-06-479 and Qkine, Qk087-0100). Molecules used throughout the loops are listed in **Supplementary Tab. 5** and concentrations and days of culture are indicated in **Supplementary Tab. 6**. Complete media changes were performed every 2-3 days. To account for the limited CD34+ cell yield from a single cord blood collection, each round of 6 to 16 designs used cells from a different donor.

#### DLL4-coated bead and lymphoid (LY) cocktail preparation

Recombinant human biotinylated DLL4 (Sino Biological, 10171-H49H-B) was resuspended in sterile water (Sigma-Aldrich, W3500-100ML) to a concentration of 250μg/mL. Streptavidin-coated polystyrene particles (1% w/v stock) (Spherotech, 0007-SVP1-200-4) were diluted 1:1 in DPBS (Sigma-Aldrich, D8537) to a final concentration of 0.5% w/v. Biotinylated DLL4 was added at a concentration of 1.5 µg per mL of bead suspension and incubated with diluted streptavidin-coated polystyrene particles on a rocker at room temperature for 30 min. Following incubation, beads were washed twice by adding DPBS followed by centrifugation at 4000 rcf for 20 min. After each centrifugation, the supernatant was slowly removed. Following the second wash, the beads were resuspended with DPBS to obtain a concentration of 15 μg/mL for downstream usage. To prepare lymphoid cocktail, beads were vortexed for 1 minute immediately before each use and added to base media, as described above, supplemented with 100 ng/mL IL7 (Miltenyi, 130093937) and 100 ng/mL rhFLt3 (Qkine, Qk087-0100).

#### Spectral flow cytometry

##### Sample Preparation

At indicated timepoints, samples were collected into polystyrene tubes (Corning, #352058) or V-bottom plates (Greiner Bio-one, #651261) and centrifuged at 300 rcf for 6 minutes at 4°C. Samples were stained with 50 μL of fluorochrome-conjugated antibody cocktails: CD49c BUV395, CD41a BUV563, CD117 BUV615, CD38 BUV661, HLADR BUV737, EPCR BV421, CD135 BV480, CD56 BV605, CD15 BV650, CD123 BV711, CD45 BV785, CD16 SparkBlue550, CD7 RealBlue780, CD90 PE, CD10 PE-Dazzle594, CD14 PE-Fire700, CD45RA PE-Cy7, CD34 APC, CD235a AlexaFluor700, and CD19 APC-Cy7 (**Supplementary Tab. 7**) and incubated for 30 minutes at 4°C and washed once with DPBS and stained with viability dye 7AAD (Fisher Scientific, #A1310) prior to data acquisition. Stained, unstained, fluorescence-minus-one (FMO) cells, and single-stained compensation beads (Thermo Fisher, #01-2222-41) were run on Sony ID7000 Spectral Cell Analyzer (Sony Biotechnology) with 5-lasers (355 nm, 405 nm, 488 nm, 561 nm, 637 nm) and 147 detectors.

To assess potential batch effects and ensure consistency, we used the instrument’s built-in standardization mode and repeatedly profiled control designs at multiple timepoints throughout the screen, following established flow cytometry best practice^66^. Control designs showed some variability in population frequencies across donors (**Fig. 2C**), consistent with donor-to-donor biological variation, though drift cannot be fully excluded given the study design. Given the stable instrument performance, no batch correction was applied to the spectral flow cytometry data. Spectral unmixing followed by per-feature normalization placed marker and scatter measurements from all conditions on a common scale, enabling their combination with LabCompass.

##### Pre-processing and Quality Control of the raw spectral flow data

Acquired data were exported as Flow Cytometry Standard (FCS) files (version 3.1) and were later imported into R (version 4.5.2) for denoising and spectral unmixing. Multiple distinct autofluorescence profiles were extracted from unstained samples of each condition using unsupervised clustering FlowSOM (version 2.18.0)^67^ implemented through *AutoSpectral* (version 1.0.0)^68^. Spectral signatures of each of the fluorochromes were isolated from single-stained compensation beads to create the spectral library. By utilizing the corresponding unstained samples for each condition, *AutoSpectral* applied the best-fitted unmixing matrix for each cell via minimal residual error matching to reduce unmixing-dependent spreading error.

Each fluorochrome channel was transformed using Logicle transformation implemented through *flowCore* (version 2.22.1)^69^ where the parameters were calculated using singlets and live cells obtained through *ggcyto* (version 1.38.1)^70^. Transformation was then applied to all ungated cells. Automatic detection of outliers due to unstable flow rate, signal drift, and other technical issues were performed using *PeacoQC* (version 1.20.0)^71^. Detected outliers were subsequently removed from untransformed *Autospectral* unmixed samples and saved as FCS files.

##### Gating, Logicle transformation and dimension reduction of the unmixed spectral flow data

Unmixed FCS files were read as annotated data objects using Pytometry (v0.1.6), metadata was assigned using a lookup table based on their experiment identifier, see **Supplementary Tab. 6**. Additional metadata from the cytometer were assigned programmatically (including acquisition date, well of origin, and instrument identifier). The files were appended and the marker labels were renamed for simplicity, followed by removal of debris, single cell filtering, and dead cell exclusion using a series of programmatically predefined gates (debris: FSC-A versus SSC-A; singlets: FSC-A versus FSC-H; viability: 7-Aminoactinomycin D; gating strategy shown in **Supplementary Fig. 15**). The viability parameter was used for this exclusion step only and was not retained as a clustering feature in subsequent analysis. Twenty fluorescence minus one (FMO) files (one per marker in the panel) were also run through the same pipeline, with the addition of a pipeline for FMO automated extraction of constants for Logicle scale transformation (applied uniformly across all experiments). The single cell object of the experiment designs were then further soft cleaned (removal of cells with negative values in all markers, and outliers, without additional event gating) before applying the Logicle transformation from Pytometry^72^, using the extracted constants from the FMOs pipeline. No additional computational batch correction was applied across experiments; batch consistency was maintained through standardized spectral flow cytometry acquisition settings and uniform application of FMO derived Logicle transformation parameters across all experiments (**Supplementary Fig. 16**). The transformation was applied channel by channel using `pytometry.tl.normalize_logicle` (Pytometry 0.1.6). The four Logicle parameters were derived per channel together from the FMO controls and the fully stained samples (the cutoff c_i of marker i is defined as the 99.5th percentile of its FMO distribution; the display range T_i is the 99.75th percentile of the fully stained signals; *M*_*i*_ = *log*_10_(*T*_*i*_), *W*_*i*_ = *log*_10_(*c*_*i*_), *A*_*i*_ = −*log*_10_(*c*_*i*_) = − *W*_*i*_). Data was further processed using Scanpy’s^73^ (v1.12) preprocessing scale (no clipping was applied), principal component analysis (n=19, corresponding to the number of clustering markers in the panel (20) minus one, to maximize use of degrees of freedom), neighbors (n=15) and UMAP using default settings (Scanpy v1.12 defaults: min_dist=0.5, spread=1.0, random_state=0). Clustering was performed using Leiden clustering^74^ (leidenalg v0.11.0, resolution=0.8, random_state=51) using a downsampled file of 500k cells, with each experiment contributing an equal number of cells (4,098 cells per experiment). Cell type annotation was performed at the cluster level, based on literature-defined marker expression profiles (**Fig. 2F, Supplementary Fig. 3**), with expert curation using cell-surface markers.

#### Applied transformation for the modeling tasks

Spectral flow cytometry events capture two complementary types of features: immunophenotypic marker expression and morphological scatter properties of the cellular states. In all experiments, cellular states 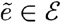 were constructed as the concatenation of the 20 marker features in the immunophenotypic panel 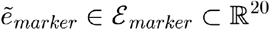, with the 6 morphological scatter features 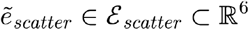; that is, 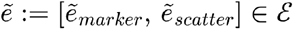, with *ϵ* := *ϵ*_*marker*_ × *ϵ*_*scatter*_. These two modalities exhibit significantly different ranges, making their contribution to the modeling task unbalanced. To rescale the values comparably, we applied independent *z*-standardization; each dimension 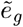 of the raw 26-dimensional spectral flow cytometry space was shifted and rescaled using the marginal mean μ_*g*_ and standard deviation σ_*g*_, estimated from the training data. This yielded the *z*-standardized cellular states:

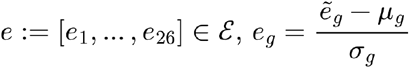

The experimental design space modeled for the inverse tasks consisted of the continuous concentration levels of 10 molecules and cytokines, along with the oxygenation level and the culture duration measured in days; therefore, the experimental design space is 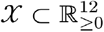. Because titration experiments usually involve exponential increments of the concentration levels, we applied the log1p transformation to the raw design values, which also accommodates zero concentration values. Formally, let 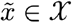 denote the raw concentration values for experimental designs; we then constructed the log1p-scaled designs as 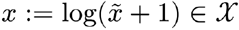. Throughout the Methods, the dimensionality of *X* counts every modelled axis, culture duration included. The schematics in Figs. 2B, 3A, 4A, 5A and 6A instead report the number of medium components together with the oxygenation level, listing culture duration separately, and therefore state 11 design parameters before the Loop 3 expansion and 13 thereafter.

#### The definition of the surrogate forward model for *in vitro* hematopoiesis

We employed a CFM-based surrogate forward model for the hematopoietic experiments; the continuous-time formulation yields marginal probability paths conditioned on experimental designs *x* ϵ *X* – we denote this path as {*p*_*t*_(·|*x)*}_*t*ϵ[0,1]_, where each element *p*_*t*_(·|*x)* ϵ *P* (ε)is a conditional distribution on the cellular state space ε for all *t* ϵ[0,1]. To learn the forward model, we chose the conditional probability path *p*_*t* ∣1_ (·|*e) = N (*·|*te*, (1−*t*)^2^I_ε_) – the conditional probability path is independent of experimental designs X ϵ x, given cellular states *e* ϵ ε. The forward model’s generating velocity field was learned with a simple modification of the standard CFM objective:

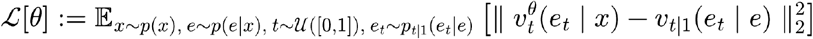

The neural architecture of the velocity field employed an MLP cell state encoder, *f*_*cell*_ (·), mapping cellular states *e*_*t*_ ϵ *ε* to their encoded representation *z*_*cell*_ : = *f*_*cell*_(*e*_*t*_). The continuous-time index t ϵ [0,1]was instead passed through the time features extractor τ_*time*_(·), to create spectral features 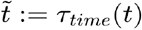 which were then further encoded using the MLP time encoder *f*_*time*_(·), mapping spectral features to their encoded representation 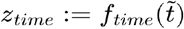. Experimental designs *x* ϵ *X* were also processed using a condition encoder *f*_*design*_ (·), mapping input designs to their encoded representation *z*_*design*_: = *f*_*design*_ (*x*). To combine these three elements, a conditioning layer *f*_*cond*_(·, ·, ·), taking as input the encoded representation of cellular states *z*_*cell*_, continuous-time index *z*_*time*_, and experimental designs *z*_*design*_, was used to create the conditioned representation 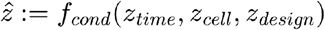. The conditioned representation 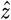 was then passed through a velocity field decoder *f*_*velocity*_ (·), to obtain the velocity 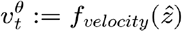. All the MLP blocks employed a Scaled Exponential Linear Unit (SELU) activation function. All the surrogate forward models were trained for 2,000,000 microbatches, using the Adam optimizer with learning rate of 0.0001. See **Supplementary Tab. 8** for more details on the explored grid.

**Supplementary table 8:**
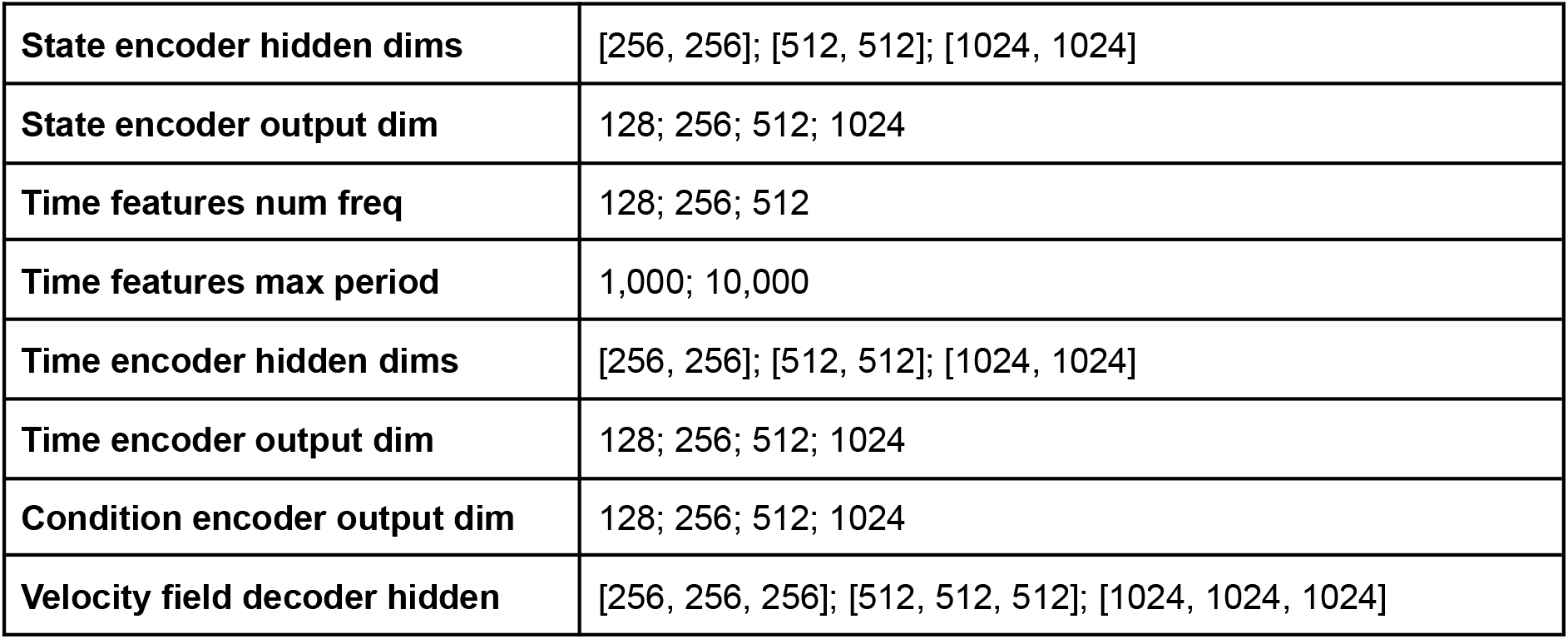
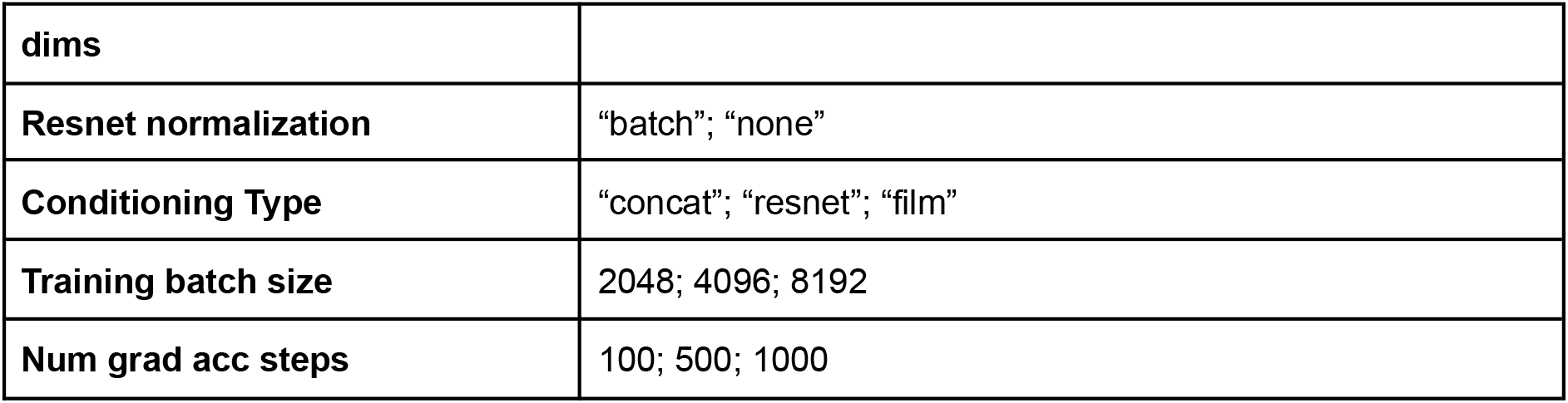
The hyperparameters grid used to sweep the forward model configurations for the biological experiments. 25 runs were executed, using random search.

#### The phenotypic readout function for cell-type composition optimization

To approximate the phenotypic readout model for the cell-type composition experiment, we used an MLP cell-type classifier, mapping cellular states *e* ϵ ε to points on the *K*-dimensional probability simplex *y* : =Δ^*K*^; the architecture consisted of a state encoder *f*_*state*_, extracting an encoded representation of the cellular states *z*_*state*_ : = *f*_*state*_ (*e*). To obtain the class logits *z* ϵ ℝ^*K*^, the encoded cellular states were passed through a linear head *W* to yield the logits as *z* = *z*_state_ *W*. Cell-type probabilities were then computed from the logits using the softmax activation. To increase the level of regularization, we added to the hyper-parameter search grid an optional dropout with 0.1 probability on the state encoder connections, and different values of weight decay for the optimizer. All the MLP blocks employed a Scaled Exponential Linear Unit (SELU) activation function. All the phenotypic prediction models were trained for 100,000 gradient steps, using the Adam optimizer. See **Supplementary Tab. 9** for more details on the hyperparameter grid.

**Supplementary table 9:** The hyperparameters grid used to sweep the phenotypic readout function configurations for the biological experiments. 30 runs were executed, using random search.

|  |  |
| --- | --- |
| **State encoder hidden dims** | [128, 128]; [128, 128, 128]; [256]; [256, 256]; [256, 256, 256] |
| **State encoder output dims** | 16; 32; 64 |
| **Use batchnorm** | False; True |
| **Dropout rate** | 0.0; 0.1 |
| **Optimizer learning rate** | 0.0005; 0.0001 |
| **Use class weights** | False; True |
| **Training batch size** | 32; 64; 128; 256; 512; 1024; 2048; 4096; 8192 |

#### Learning the prior model over the design space of *in vitro* hematopoiesis

To approximate the prior distribution over the experimental design space, we employed a CFM model – we used the conditional probability path *p*_*t*|1_(·|*x*) := *N*(·*tx*,(1 − *t*)^2^ I*x*) to learn the neural prior velocity field 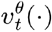. The architecture of the neural prior velocity consisted of a design encoder *f*_*design*_(·), mapping designs *x* ∈ *X* to their encoded representation *z*_*design*_ := *f*_*design*_(*x*). The continuous-time index *t* ∈ [0,1] was instead passed through the time features extractor *τ*_*time*_(·), to create spectral features 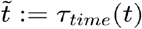 which were then further encoded using the MLP time encoder *f*_*time*_(·), mapping spectral features to their encoded representation 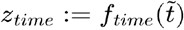. To combine these two elements, a conditioning layer *f*_*cond*_(·, ·), taking as input the encoded representation of the continuous-time index *z*_*time*_ and experimental designs *z*_*design*_, was used to create the conditioned representation 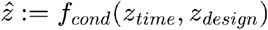. The conditioned representation 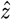 was then passed through a velocity field decoder *f*_*velocity*_(·), to obtain the velocity 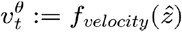. The CFM prior was trained for 200,000 steps and its parameters were updated using the Adam optimizer with learning rate of 0.0001. The flow map prior architecture entailed an additional encoder for the extra time index *s ∈* [0,1], denoted by 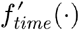 and configured with the same settings as for its counterpart *f*_*time*_(·). Both time indices shared the same time features extractor *τ*_*time*_(·); hence, the architecture first extracts the time features as 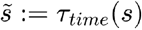, then encodes them using the additional MLP block, to obtain 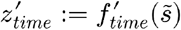, which is then concatenated with *z*_*time*_, to obtain the concatenated representation 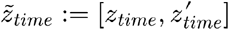. The conditioning layer then takes this concatenated representation to create the conditioned features 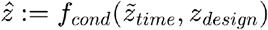. The flow map decoder *f*_*flow map*_(·) works analogously to the velocity field decoder and it is initialized with the same settings. The configurations to distill the flow map prior were the same ones used to train the prior velocity field. See **Supplementary Tab. 10** for the hyperparameter configuration used for the prior model.

**Supplementary table 10:** The hyperparameters used to train the prior velocity field configurations for the biological experiments.

|  |  |
| --- | --- |
| State encoder hidden dims | [256] |
| State encoder output dim | 16 |
| Time features num freq | 256 |
| Time features max period | 100 |
| Time encoder hidden dims | [256] |
| Time encoder output dim | 8 |
| Velocity field decoder hidden dims | [256] |
| Resnet normalization | “none” |
| Conditioning Type | “resnet” |
| Num Resnet blocks | 3 |
| Training batch size | 2048 |
| Num grad acc steps | 20 |

#### Optimizing cell-type composition: experimental configurations

For all experiments, we used Euler discretization with the number of integration steps *S* ∈ {100,200, 500} to simulate LabCompass’s inverse ODE. The capping value for the coefficient 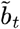 that rescales the potential gradients was swept over *b*_max_ ∈ {1.0, 5.0, 7.0, 10.0}. We set *M* = 350 for the number of samples used to approximate the response.

### The sequential experimental design loop

LabCompass was applied iteratively, alternating between *in silico* design generation and wet-lab validation. Each loop comprised four stages: generation of candidate designs by the inverse model for a specified target cell state; selection of a small number of candidates for experimental evaluation; culture and spectral flow cytometry profiling of the selected designs; and retraining of the models on the augmented dataset before the subsequent query. Loop 1 served as an in silico validation of retrieval and model updating: six previously measured designs (205–210), spanning a range of distances from the training data, were withheld from training and used as ground truth, and no designs generated by LabCompass were cultured. Loops 2 to 4 were model-informed wet-lab loops, in which LabCompass-generated candidates were selected, cultured and profiled. Each loop comprised 6 to 16 designs in total. All designs evaluated in each loop, with their parameter values, are listed in the **Supplementary Tab. 6**.

#### Generation of candidate designs

For each target cell state, the inverse model was queried as described in “Optimizing cell-type composition: experimental configurations”, drawing 50 candidate designs per run. Generation was swept exhaustively over the number of integration steps (*S* ∈ {100, 200, 500) and over the guidance-strength schedule. Two schedules were used: the reciprocal schedule derived in **Theoretical Background**, *b*_*t*_ = (1 − *t*)/*t*, capped at *b*_max_ ∈ {1, 5, 7, 10}; and a constant schedule, *b*_*t*_ := *b*_max_ with *b*_max_ ∈ {1, 3, 5, 7, 10}. This gave 12 and 15 unconstrained runs per target, respectively. Candidates were generated both without constraints and under the experimental-bound penalty described in “Enforcing the experimental constraints”, in two variants penalising oxygenation and culture duration alone, or all design axes. The penalised grids additionally swept the penalty warm-up (*t*_warmup_ ∈ (0, 0.1, 0.25, 0.5}) and strength bounds (*υ*_min_ ∈ {1, 2, 5}; *υ*_max_ ∈ {7.5, 10, 15}); these grids exceed the 225-trial budget and were therefore sampled rather than exhaustively enumerated. Candidates from all runs were pooled per target cell state before filtering. Each candidate *x* was then characterized by its predicted target-cell-type proportion 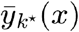 and its predictive uncertainty, both evaluated under the current forward surrogate using the loss-function uncertainty *u*(*x*) readout (see “Forward model’s uncertainty quantification”) with *N*_pop_ = 10 generated populations of 5,000 synthetic profiles each. This post hoc characterization of generated candidates is distinct from the response approximation used within the inverse ODE itself, for which 350 synthetic profiles were drawn per guidance step.

#### Filtering of generated candidates

Before selection, candidates were discarded if the predicted proportion of the target cell type fell below a per-target threshold; if oxygenation lay outside 5–25% or culture duration outside 12–20 days; or if any design parameter fell outside the experimentally measured range for that axis, allowing a fractional margin. Because targets differed in attainable enrichment, the proportion threshold was set per loop and per target cell type, ranging from 0.01 for rare lineages to 0.20 for lineages already well represented in the screen. The bound margin was set to 0.5 of the measured range for a given axis.

#### Selection of designs for experimental validation

To meet practical experimental constraints, we select a subset of possible designs for wet-lab validation. Because filtering had already removed candidates with low predicted enrichment, selection among the surviving candidates was made on predictive uncertainty alone, quantified as the standard deviation of the loss across the generated populations. Filtered candidates were ranked by this uncertainty, and for each target lineage we selected the single lowest-ranked candidate as the exploitative design and the two highest-ranked candidates as the exploratory designs. Each loop therefore both banked an expected gain and resolved the region in which the surrogate was least certain.

From Loop 3 onward, the several hundred candidates with the lowest predictive uncertainty for a given target were retained and clustered by hierarchical clustering with average linkage on Euclidean distances in design space. The dendrogram was cut into a fixed number of clusters, between 4 and 7 depending on the target, chosen by visual inspection of the dendrogram. Low-uncertainty representatives were drawn from distinct clusters, so that the selected designs probed different regions of the design space rather than a single mode. In Loop 4, clusters with similar mean predicted target-cell-type proportions were treated as alternative routes to the same target state and compared directly; the lowest-uncertainty member of each such cluster was selected, yielding designs 261 and 263 for the Mgk-Ery lineages and designs 267 and 268 for CD16+ monocytes.

#### Model update between loops

Following each round of measurements, the newly acquired spectral flow cytometry events were pooled with all data from previous loops. The forward surrogate was retrained on the pooled dataset at each loop, re-running the hyperparameter search over the grid given in **Supplementary Tab. 8** (25 random-search runs) and retaining the configuration with the best validation performance, so that the surrogate architecture was re-selected at every loop rather than carried over. The design prior was likewise retrained on the pooled dataset at each loop, using the fixed configuration given in **Supplementary Tab. 10**, and was trained from the forward-model checkpoint fitted to that loop’s data. Unlike the forward surrogate and the design prior, the phenotypic readout classifier was not retrained at every loop. It was updated only when the cell-type annotation it reproduces was revised, and reused unchanged otherwise.

#### Expansion of the design space

In Loop 3, two parameters absent from the initial screen, M-CSF and the lymphoid (LY) cocktail, were added to the design space, increasing its dimensionality from 12 to 14 (including oxygen and number of days). Designs from earlier loops were assigned a value of zero along these axes, the concentration at which they were in fact held. Because no measurements existed along these axes, the expanded space was first populated experimentally: design 247 was supplemented with M-CSF at 0.2, 2 and 20 ng/mL (designs 258, 257 and 256, respectively), and designs 245 and 247 were supplemented with the LY cocktail at 1.5 and 15 µL/well (designs 252 to 255). These measurements were added to the training data before the prior and forward model were retrained over the expanded space and queried for the new target lineages.

#### Held-out evaluation of model improvement

To quantify the improvement in forward-model performance attributable to a loop’s measurements, a subset of validated designs was withheld from the model update and used as an evaluation set. In Loop 1, six designs (205 to 210) spanning a range of distances from the training data were held out before training. In Loop 2, four target lineages were queried and one uncertain design per lineage was withheld: designs 248, 249, 250 and 251 for MgkPro, EryPro, pHSC and Pre-ProB, respectively. Hold-out prediction analysis was only carried out on MgkPro- and EryPro-enriching designs, as they showed promising experimental results. Predictions from the updated model were then compared with those of the model that had not observed the new measurements, in terms of the Pearson correlation between predicted and measured marker statistics and of the predictive uncertainty *u*(*x*).

#### Counterfactual parameter masking

To assess whether a predicted enrichment was attributable to a specific design parameter, generated designs were re-evaluated with that parameter set to zero and the resulting change in predicted target-cell-type proportion was recorded. This masked evaluation was applied to M-CSF across the CD14Mono- and CD16Mono-enriching designs generated in Loop 3 (**Fig. 5C**).

#### Controls

In Loop 4, design 279, a re-measurement of design 210, served as the positive control for Mgk-Ery enrichment, and design 277, a repeat measurement of the unsupplemented base condition 247, served as the positive control for CD16+ monocytes.

### Inverse design of viral expression states in HCMV infection

#### Dataset

We applied LabCompass to the single-cell HCMV infection time course of Hein and Weissman^57^, in which human fibroblasts were infected and profiled across 8 time points (12,919 cells), together with its accompanying Perturb-seq host-factor screen. Following the original study, genes were split into host and viral sets and filtered to those with more than 10,000 total counts, yielding 3,588 genes of which 3,482 were host genes. The 21 knockdown targets of the screen were retained in addition. Host counts were normalized to 10,000 counts per cell and log1p-transformed. Viral counts were log1p-transformed only, as viral load is itself the signal of interest. Each modality was reduced to 50 principal components, with loadings and means retained so that generated components could be decoded back to gene space. Infection-stage labels were taken from the original annotation. In this application the design space is the viral expression space and the cellular state space is the host expression space, both 50-dimensional.

#### Models

The forward model was a CFM model mapping an isotropic Gaussian to host components, conditioned on the viral state of the cell, with a one-hot encoding of the 8 time points as an additional conditioning branch for the Perturb-seq experiments. Its architecture followed the hematopoietic forward model, with a ResNet-conditioned velocity decoder (3 blocks, SELU activations) and sinusoidal time features. It was trained for 400,000 gradient steps with Adam, learning rate 0.0001 and batch size 256, holding out 10% of cells for validation. The prior over viral expression was an unconditional CFM model on the 50 viral components, trained for 50,000 gradient steps with Adam, learning rate 0.001 and batch size 1,024. Because infection stages are strongly imbalanced, training batches were stratified over Leiden clusters of the viral components.

#### Inverse queries

Phenotypic readouts were the log_2_ fold changes of query host genes between the generated population and naive cells, and the guiding loss function was the mean squared error between generated and target fold changes. The linear one-step estimator was used in place of the distilled flow map, the design space being low-dimensional and the prior trajectories short. Query genes were curated from the original study and their target fold changes measured in the data. For the Infected abortive target these were NFKBIA, JUN, IER2, DDIT3, CCNL1 and IRF1. Guidance strength and the number of integration steps were swept for each query, and generated viral states were assigned to infection stages by nearest neighbors in viral principal-component space.

#### Held-out phenotype retrieval

Each of the clusters Infected 1 to 6 was removed in turn and the embeddings, prior and forward model refit on the reduced data. LabCompass was then queried with the log_2_ fold changes of CD55 and HES6 measured between the held-out cluster and naive cells, over three replicates, retaining per cluster the configuration that minimized the mean energy distance between generated and held-out profiles in viral principal-component space. As baselines we report the energy distance between the held-out cluster and its closest and second-closest observed clusters, and a random subset of training viral states. Synthetic knockdown of host restriction factors. Using the Perturb-seq arm of the same study and the time-conditioned forward model, the potential was set to the absolute decoded expression of a target host gene in the generated host population, so that guidance drives the viral state toward conditions in which the gene is not expressed. Queries were run independently for each time point. Generated viral states were pushed through the forward model with fresh noise, and the resulting host states were annotated by nearest neighbors on real host components. Stage proportions were then compared against cells carrying the corresponding guide in the screen, against non-targeting guides, and against an equally sized random sample of screen cells. Results are reported for STAT2 and IFNAR2.

### Comparison with gradient descent

#### GD as a natural baseline for gradient based optimization

Since LabCompass is a gradient-based optimization algorithm, it appears natural to perform a comparison with the simplest instantiation of gradient-based optimization algorithms: Gradient Descent (GD). In addition, recent work has employed vanilla GD^48^ for design optimization in single-cell biology studies, further establishing the necessity of such comparison.

Gradient descent is an optimization algorithm that updates its solutions based on the gradients of the target potential. For total number of steps *T* ∈ ℕ_≥0_, GD can be expressed as the update rule:

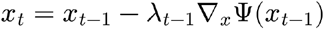

Where the sequence 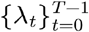, with *λ*_*t*_ ≥ 0 for all *t*, is referred to as the *learning rate schedule*. The initial sample are drawn from the initialization distribution *χ*(·) h *P*(*X*), that is *x*_0_ ∼ *χ*(·).

#### Initialization distributions and learning rate schedulers

As we have just seen, the two hyperparameters that GD entails are the initialization distribution *χ*(·) and the learning rate schedule 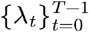. We set the initialization distribution either to an isotropic *χ*(·) := *N* (· *|* **0**_*x*_,I_*x*_) *Gaussian*, or to a mixture of Dirac unit masses centered at the unique measured designs, that is:

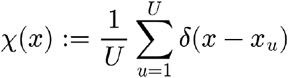

where each *x*_*u*_ is one of the *U* ∈ ℕ unique designs measured in the spectral flow cytometry dataset D. In the simplest setting, we employed a constant learning rate schedule 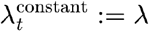 for all *t* with *λ ∈ ℝ*_≥0_. Otherwise, we employed a linear learning rate schedule: for (*λ*_min_, *λ*_min_) ∈ ℝ^2^, with *λ*_min_ < *λ*_max_, the learning rate for step *t* is defined as 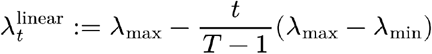. Additionally, we also used the exponential learning rate, defined in terms of decay factor *μ ∈ [*0,1] and maximum learning rate *λ*_max_ ∈ ℝ_≥0_ as 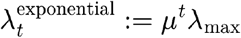. Hyperparameter configurations explored during the experiments are reported in **Supplementary Tab. 11**.

**Supplementary table 11:** The hyperparameters grid used to sweep the Gradient Descent baseline configurations for the comparison with LabCompass on MgkPro enrichment in Loop 1. The grid was exhaustively swept, and all the runs were executed.

|  | Definition | Swept Hyperparameters |
| --- | --- | --- |
| **Constant** | $\lambda_t^{constant} := \lambda, \lambda \in \mathbb{R}_{\geq 0}$ | $\lambda \in \{10.0, 1.0, 0.1, 0.01, 0.001, 0.0001\}$ |
| **Linear** | $\lambda_t^{linear} := \lambda_{\max} - \frac{t}{T-1}(\lambda_{\max} - \lambda_{\min})$<br>$\lambda_{\min}, \lambda_{\max} \in \mathbb{R}^2, \lambda_{\min} < \lambda_{\max}$ | $\lambda_{\max} \in \{10.0, 1.0, 0.1, 0.01, 0.001, 0.0001\}$<br>$\lambda_{\min} = 0.0$ |
| **Exponential** | $\lambda_t^{exponential} := \mu^t \lambda_{\max} \mu \in [0, 1],$<br>$\lambda_{\max} \in \mathbb{R}_{\geq 0}$ | $\lambda_{\max} \in \{10.0, 1.0, 0.1, 0.01, 0.001, 0.0001\}$<br>$\mu \in \{0.9, 0.99, 0.999\}$ |

#### Enforcing the experimental constraints: penalization term on the potential function

Gradient descent does not constrain the optimization trajectory using any information on the experimental design space, substantially limiting its capacity to generate realistic design configurations. To enforce such a constraint on the optimization trajectory, we added a quadratic penalization on the loss function, penalizing designs that exceeded the experimentally measured bounds. For each design axis *j* ∈ {1, …, 12}, denote by 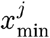 and 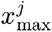 the minimum and maximum values experimentally evaluated. The quadratic penalty was then defined as:

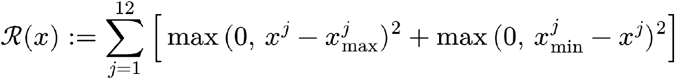

The optimization was then performed on the penalized potential Ψ_*R*_(*x*) := Ψ(*x*) + *γ*_*R*_ *R*(*x*), for scalar penalization strength ; we swept for the comparison.

### Additional Specifications

#### Sinusoidal time features

To increase the expressiveness of the neural velocity field’s time-dependency, it is common practice to use closed-form sinusoidal expressions to extract features from the time index *t* ∈ [0,1] – previously denoted by *τ*_min_(·); we provide here the exact definition of such a function. Let *P*_max_ ∈ ℕ denote the maximum period; *N*_*freq*_ ∈ ℕ represents the number of frequencies, such that *N*_*freq*_ = 2*N*_*half*_ for *N*_*half*_ ∈ ℕ. For each frequency index *n* ∈ {0, …, *N*_*half*_ − 1} we define the frequency vector:

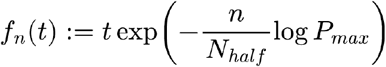

where *t* ∈ [0,1] is the time index. Time features are ultimately constructed as:

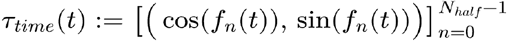

yielding an *N*_*freq*_ -dimensional vector.

#### ResNet conditioning

The ResNet layer stacks a sequence of *K*_*blocks*_ ∈ ℕ layers, to condition the latent state representation on external information. Formally, let *z* ∈ ℝ^*N*^ denote the *N*-dimensional vector that we want to condition, and denote by the *y* ∈ ℝ^*M*^ *M*-dimensional external information we want to inject in *z*.

Let 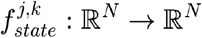 for *j* ∈ {1, 2}, and *g*^*k*^ : ℝ^*M*^ → ℝ^*N*^ denote respectively the state and condition feature extractors for block *k* ∈ {1, …, *K*_*blocks*_} – the residual block is then defined as:

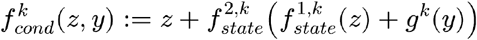

The conditioning representation is defined by the sequential composition of the 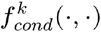 blocks. Each of 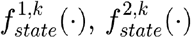 processes its input by first applying an optional batch or layer normalization, which is then passed to a Sigmoid Linear Unit (SiLU) non-linearity; next, an optional dropout is applied, with probability *p* ∈ (0, 1). The resulting representation is passed through a dimensionality-preserving linear layer. The condition feature extractor first applies the SiLU activation on the input vector, and then passes it through a linear layer, mapping to the state space dimension.

#### FiLM conditioning

Let *z* ∈ ℝ^*N*^ denote the *N*-dimensional vector that we want to condition, and denote by *y* ∈ ℝ^*M*^ the *M*-dimensional external information we want to inject in *z*. Let *W*_*γ*_ *∈* ℝ^*M,N*^ and *W*_*β*_ *∈* ℝ^*M,N*^ be two matrices. The FiLM conditioned representation is obtained as:

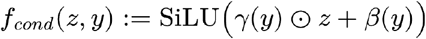

where *γ* := *y* · *W*_*γ*_ and *β*(*y*) := *y* · *W*_*β*_; SiLU(·) denotes the Sigmoid Linear Unit non-linearity.

## Data availability

The data from different experimental loops, as well as selected generated designs across lab-in-the-loop iterations are available at the following link: https://huggingface.co/datasets/theislab/LabCompass

## Code availability

The core software package for LabCompass is available at https://github.com/theislab/LabCompass. The code to reproduce our analyses on spectral flow cytometry data and sequential experimental loop is hosted at https://github.com/theislab/LabCompass-reproducibility and https://github.com/Gottgens-lab/labcompass_preprocessing.

## Acknowledgements

We are grateful to the Anthony Nolan Tissue Bank and the Cambridge Blood and Stem Cell Biobank for provision of cord blood samples and their corresponding donors. We would also like to thank NIHR Cambridge BRC Cell Phenotyping Hub. The Göttgens laboratory is supported by Wellcome (206328/Z/17/Z, 309075/Z/24/Z), Blood Cancer UK and Blood Cancer United (7035-24), and Aging Biology Foundation. Licyel Paulas-Condori is funded by the Wellcome Trust PhD fellowship [318995/Z/24/Z] and the Cambridge Trust. Deborah G. Martinuzzi is funded by the Kay Kendall Scholarship in Leukaemia Research (KKL1355) and the Cambridge Trust. This research was funded in whole, or in part, by the Wellcome Trust [226795/Z/22/Z]. This project has received funding from the European Research Council (ERC) under the European Union’s Horizon Europe research and innovation programme under grant agreements No. 101054957 (DeepCell) and 101248740 (CellCourier). This work was also supported by the Helmholtz Association within the framework of the Helmholtz Foundation Model Initiative (VirtualCell). We also thank Luca Eyring for his fruitful feedback on generative modeling for inverse problems. We are also thankful to Matthew Nicholls and Adam Wilkinson for sharing their expertise in preparing the lymphoid cocktail. For the purpose of open access, the author has applied a CC BY public copyright licence to any Author Accepted Manuscript version arising from this submission.

## Contributions

L.C., A.P., L.P.-C. and J.R.-L. developed the study, designed and performed experiments, analyzed data, and wrote the paper. A.P. and L.C. conceptualized the model, developed the LabCompass software and performed downstream analysis of the results. J.R.-L., D.G.M. and O.H. developed the pipeline for spectral flow cytometry pre-processing and processing. W.Z. provided critical analysis of the results and proofread the spectral flow cytometry analysis code. Z.W. performed experiments. D.K. and A.D. contributed to the technical and theoretical conceptualization of the framework. All authors reviewed and approved the final manuscript. N.K.W., B.G., and F.J.T. conceived, conceptualized, and supervised the study.

## Competing interests

F.J.T. consults for Immunai, CytoReason, Valinor Industries, Bioturing and Phylo Inc., and has ownership interest in RN.AI Therapeutics, Dermagnostix, and Cellarity. The remaining authors declare no competing interests.

## Supplementary information

### The prediction gap increases with dimensionality and resolution

In **Supplementary Tab. 12**, we present the numerical results for the comparison with uniform random search. This table displays the relative performance gap between the compared methods; they were both executed using the same number of function evaluations, to ensure a fair assessment of their performance differences.

**Supplementary table 12:**
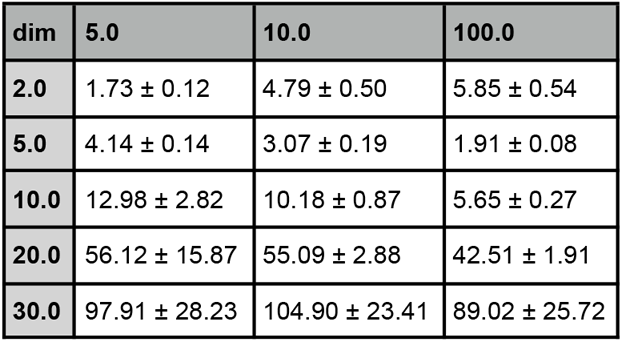
Relative performance of uniform random search with respect to LabCompass, used in the comparison as a reference method. The objective value was scaled by a factor of 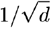 to make the results comparable across dimensions. We can observe a general trend, with the performance ratio generally worsening with the dimensionality; however, this tendency was not apparent along the resolution axes, except for the 2-dimensional problems. As the comparison involved 10 problems for dimensionality and resolution, the standard deviation was taken over such instantiations.

### The prediction quality improves between Loop 1 and Loop 2

To evaluate the approximation quality of the forward model, for each of *x*^*j*^ ϵ *X* the held-out designs, we computed the empirical mean and marginal standard deviation over *N*_*j*_ the is the number of measured cells arising from each design.

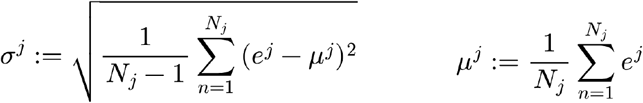

In parallel, we generated *M* ϵ ℕ_≥0_synthetic profiles using each forward model, and analogously computed their empirical mean 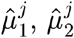 and standard deviation 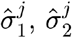. We then compared the two predicted statistics against those experimentally observed, using the Pearson correlation coefficient; we report the obtained values in **Supplementary Tab. 13**.

**Supplementary table 13:** Pearson correlation between predicted and measured mean and standard deviation across experimental loops 1 and 2 for each of the unseen conditions. The inclusion of the uncertain design significantly improves the approximation quality of the forward model on the held-out protocols for both evaluated statistics.

| Experiment Number | Mean Loop 1 | Mean Loop 2 | Std Loop 1 | Std Loop 2 |
| --- | --- | --- | --- | --- |
| 248 | 0.440 | 0.933 | 0.458 | 0.814 |
| 249 | 0.434 | 0.812 | 0.264 | 0.930 |

### The predictive uncertainty decreases between Loop 1 and Loop 2

Similarly, to investigate the beneficial effect of the uncertain design on the predictive uncertainty of the forward model across loops, we quantified the uncertainty of the loss function, that is we used ω(*y*) = *ψ*(*y, y*^⋆^) for the two models. To compute it, we generated *M* = … synthetic cellular profiles and independent populations. Compared to the model trained in loop 1, the model trained in loop 2, which includes the uncertain design, showed a consistently lower predictive uncertainty on unseen protocols (**Supplementary Tab. 14**), indicating that acquiring data in regions of high model uncertainty effectively improves the model’s confidence where it was previously least informed.

**Supplementary table 14:**
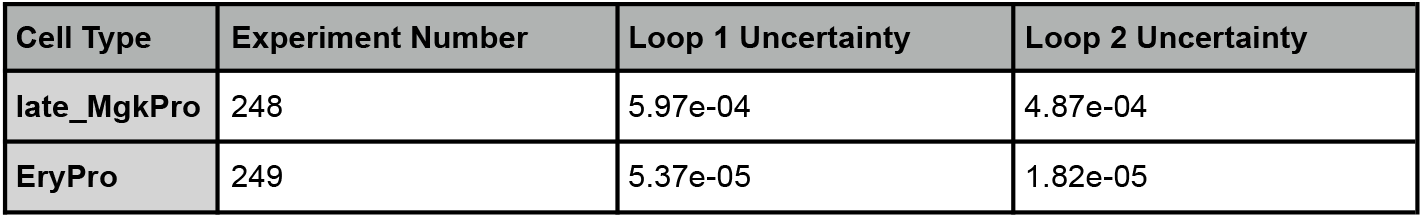
Loss function predictive uncertainty across experimental loops 1 and 2. The inclusion of the uncertain design significantly reduces the model’s uncertainty on the unseen protocols.

### Molecules and cytokines used throughout screening

**Supplementary table 5:**
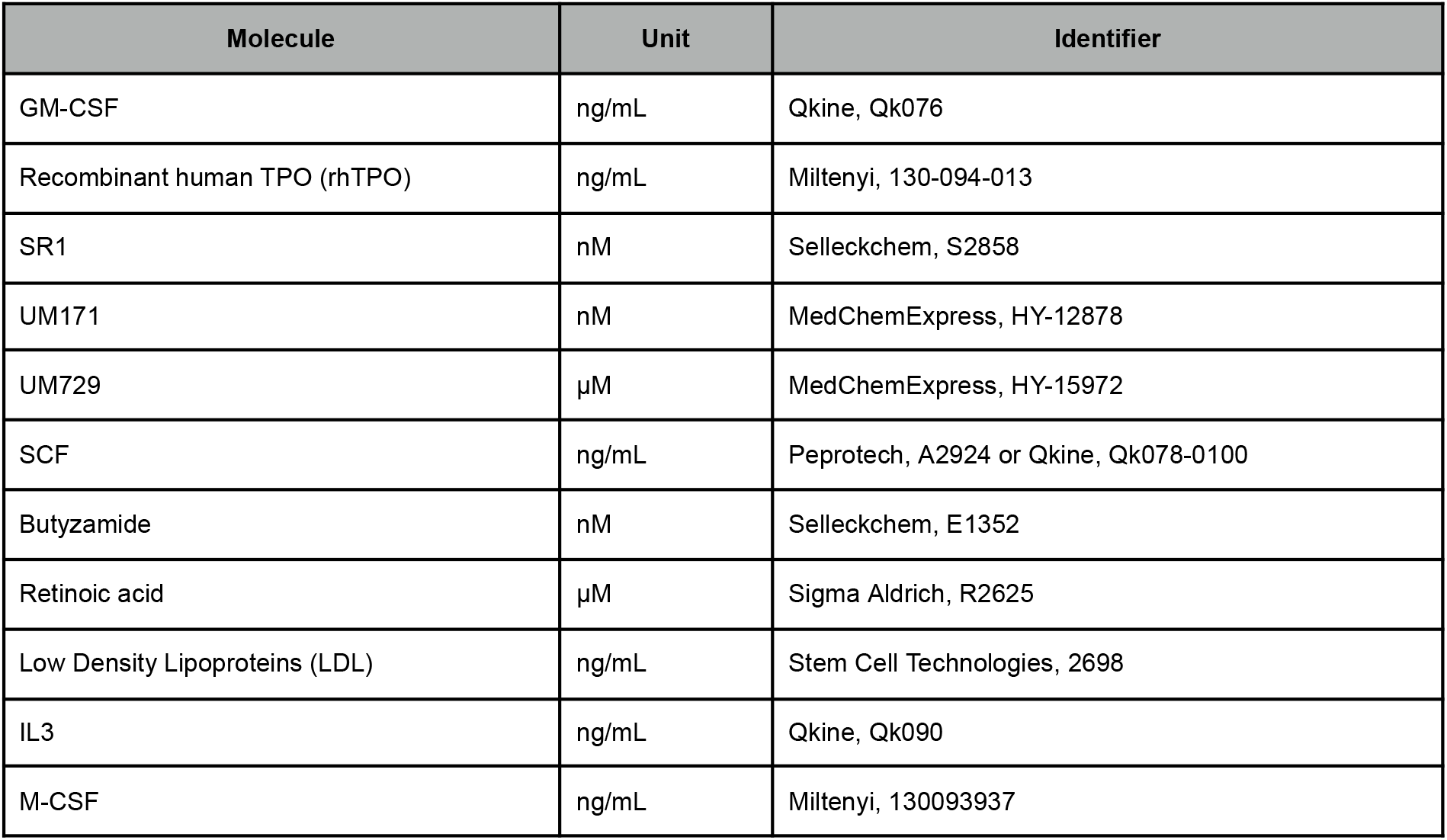
Molecules and cytokines used in addition to the basal media across all loops.

### Design number and parameters

LabCompass was initially trained on 116 designs (40-189 & 1089-4139). For Loop 1, designs 205 to 210 were analyzed. For Loop 2, designs 239 to 251 were analyzed. For Loop 3, designs 252 to 258 and 270 - 275 were analyzed. For loop 4, design 259-269 and 276-280 were analyzed.

**Supplementary table 6:**
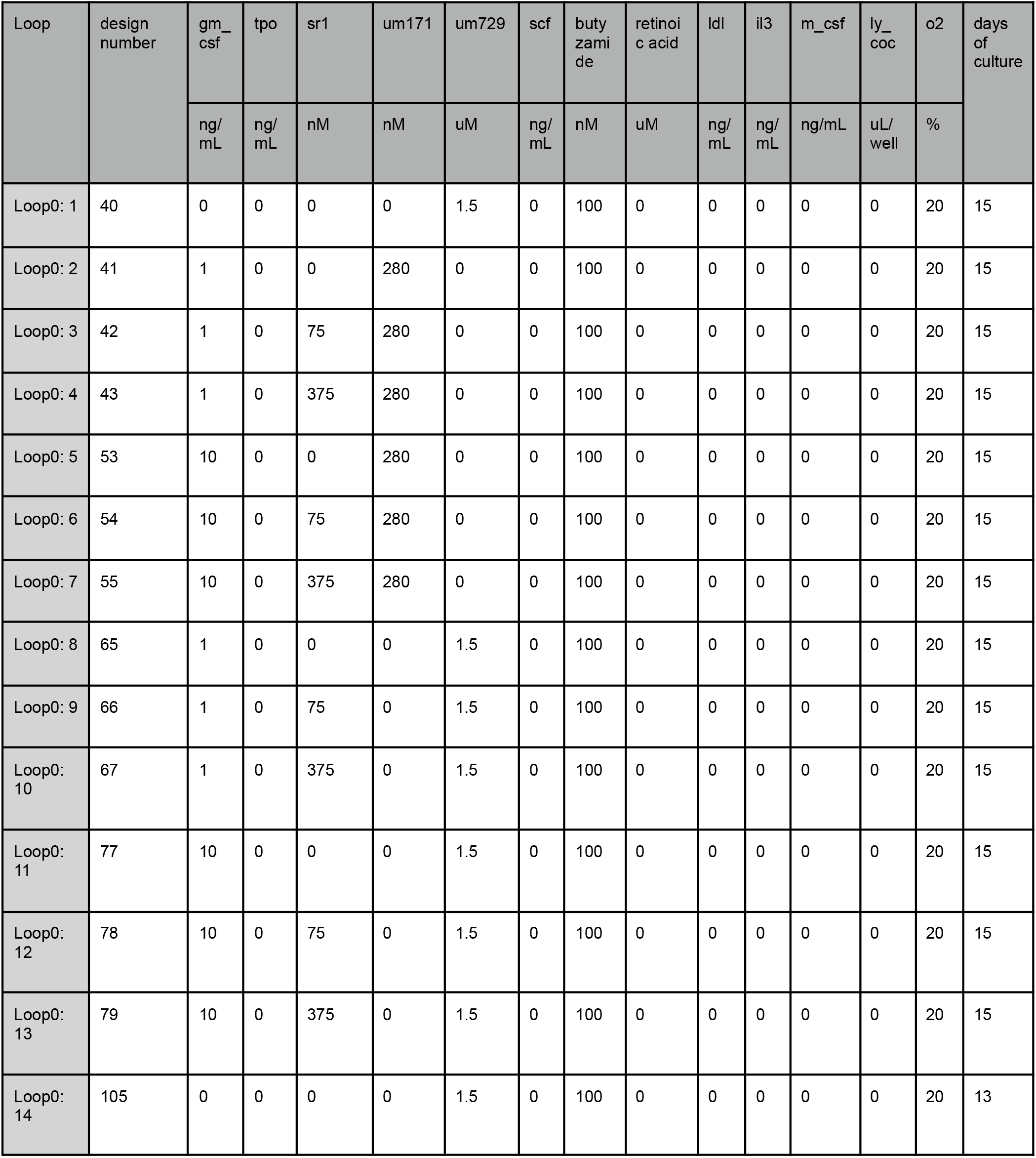
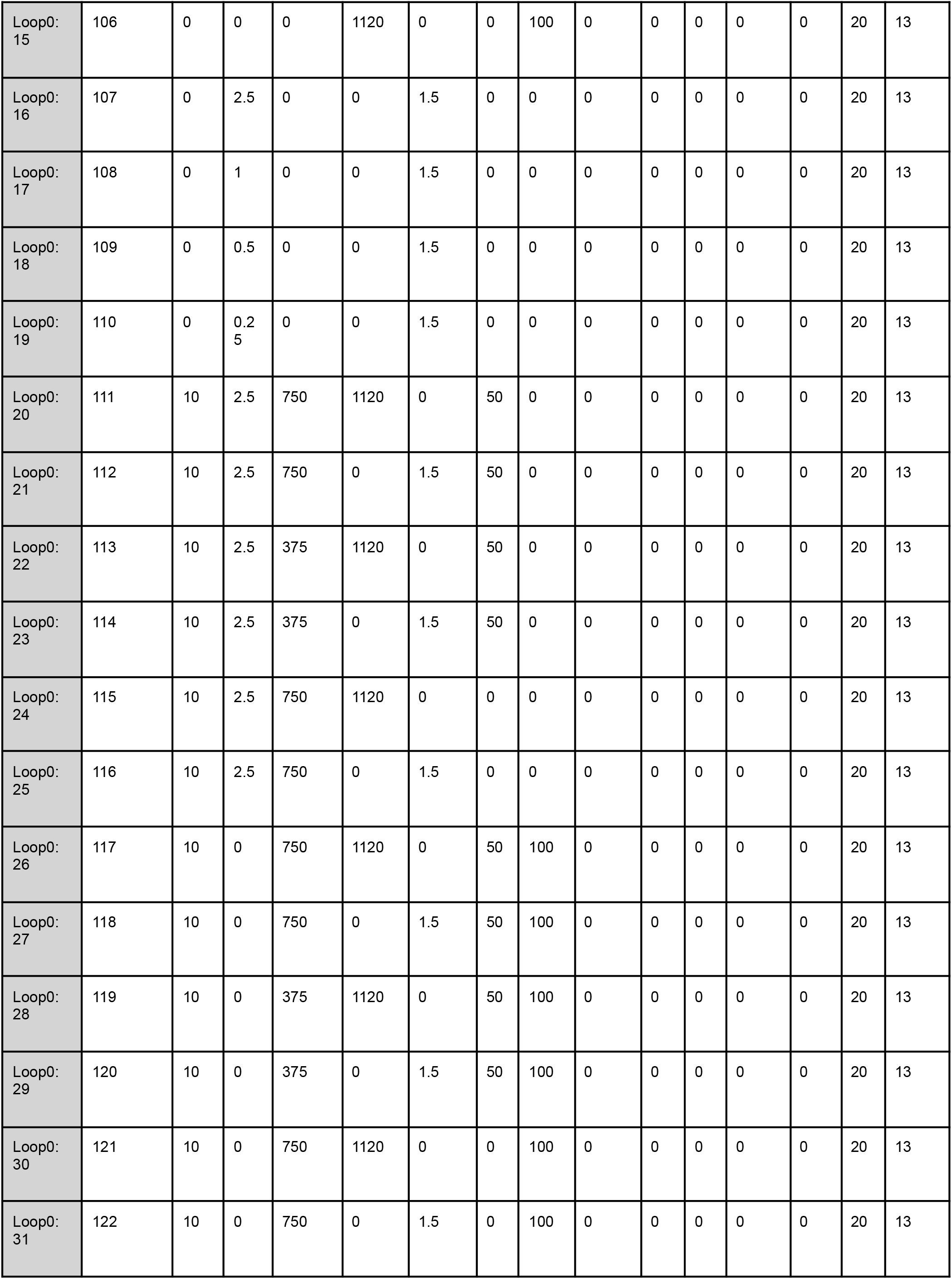
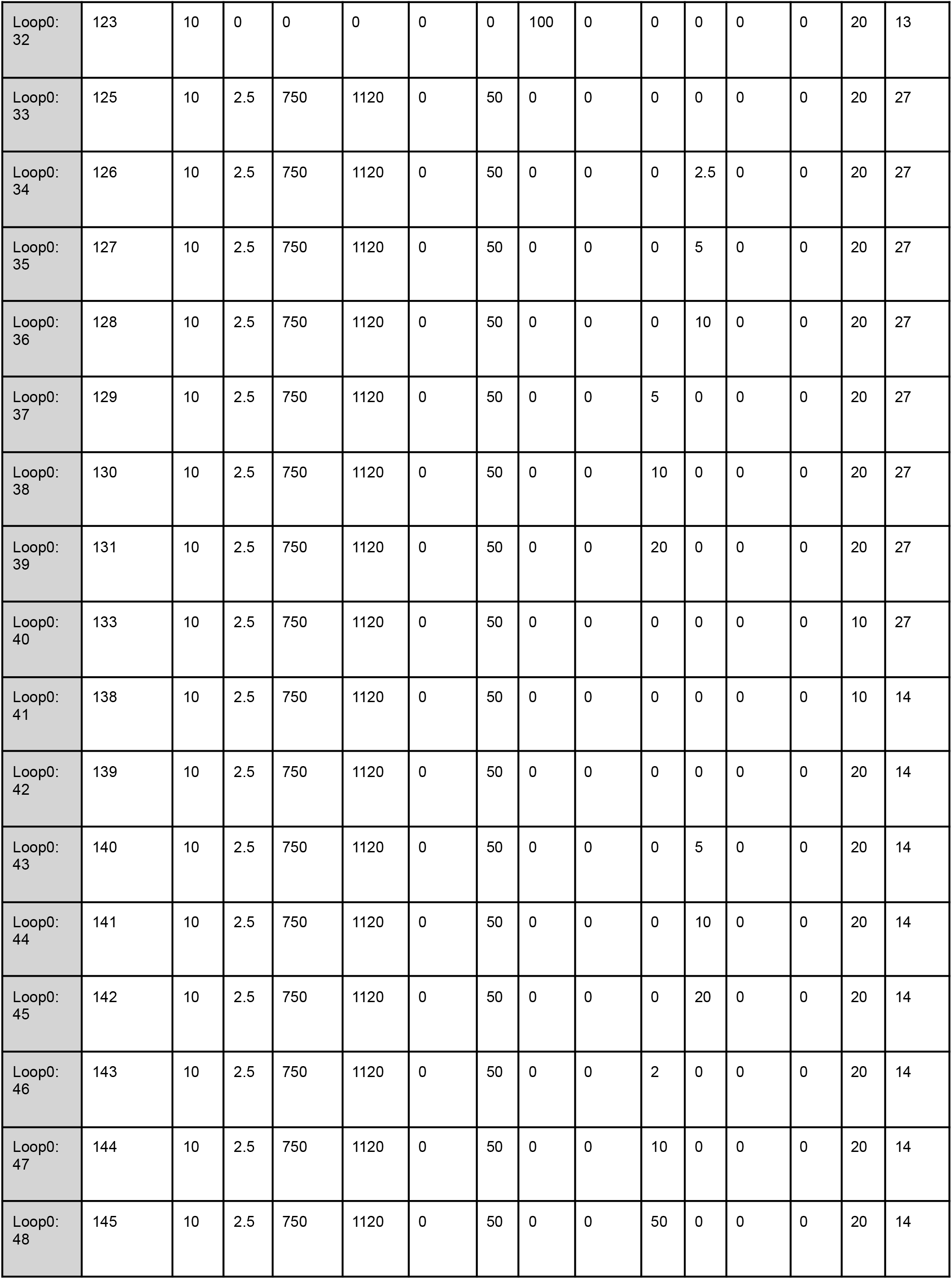
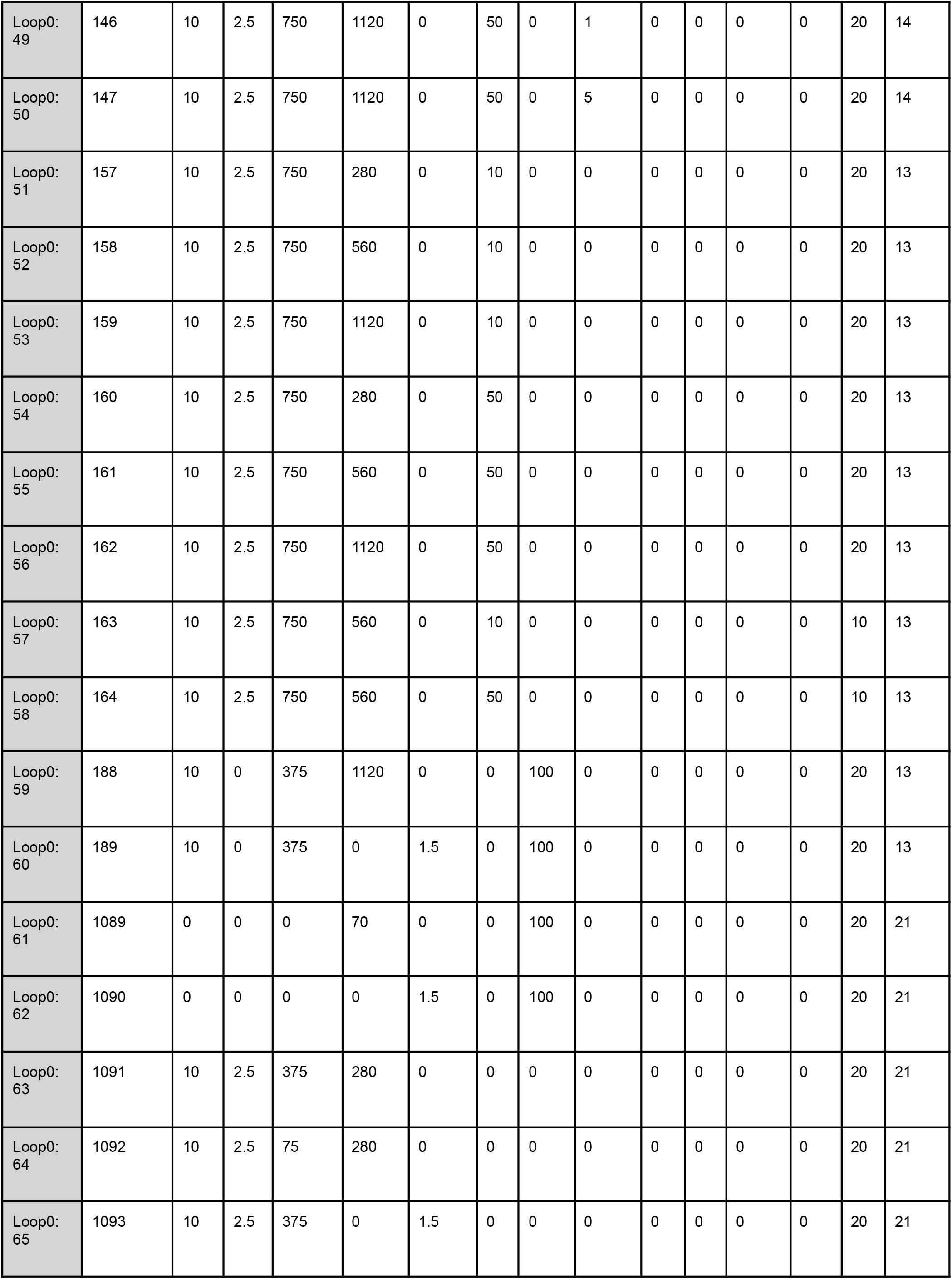
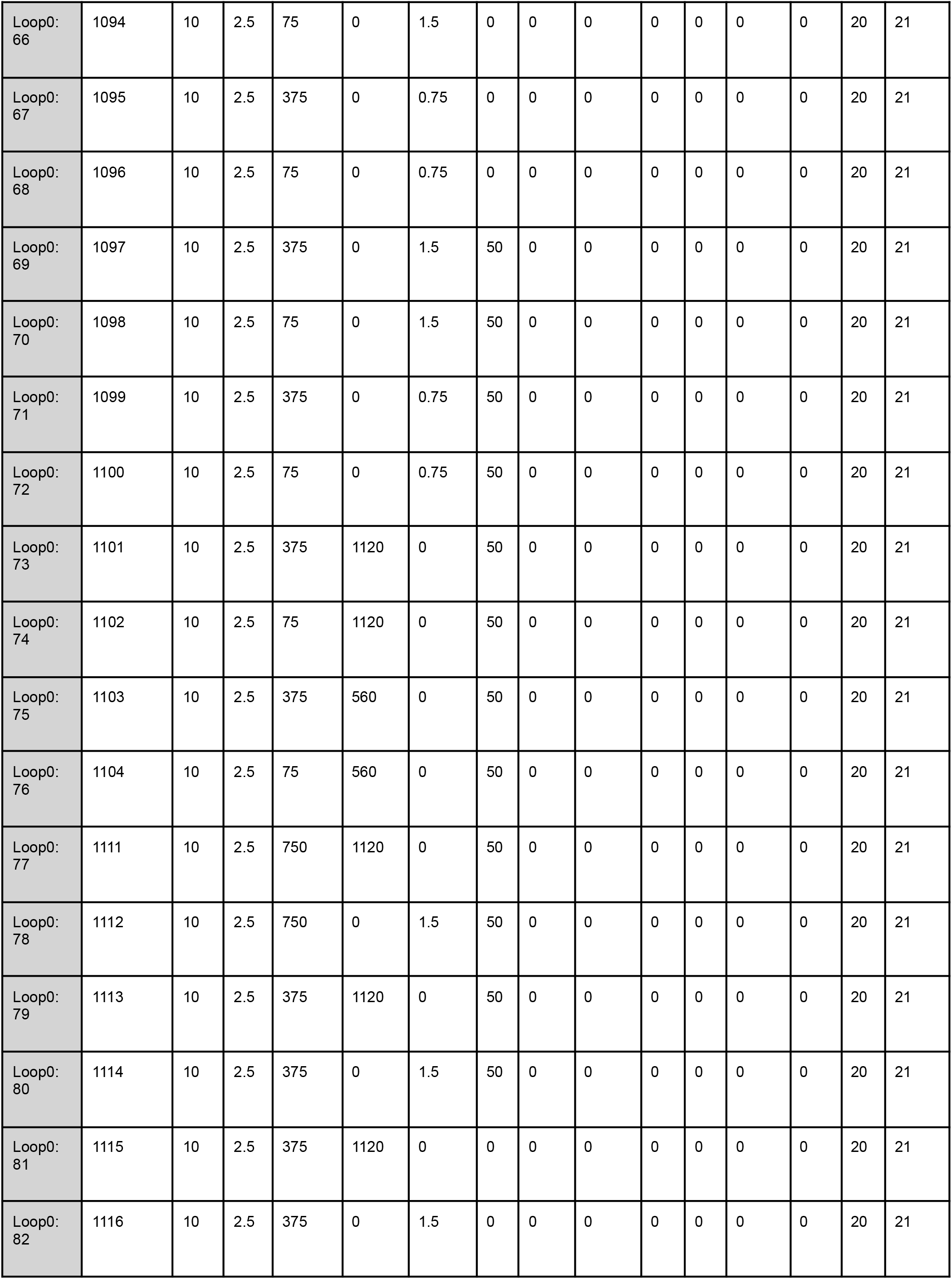
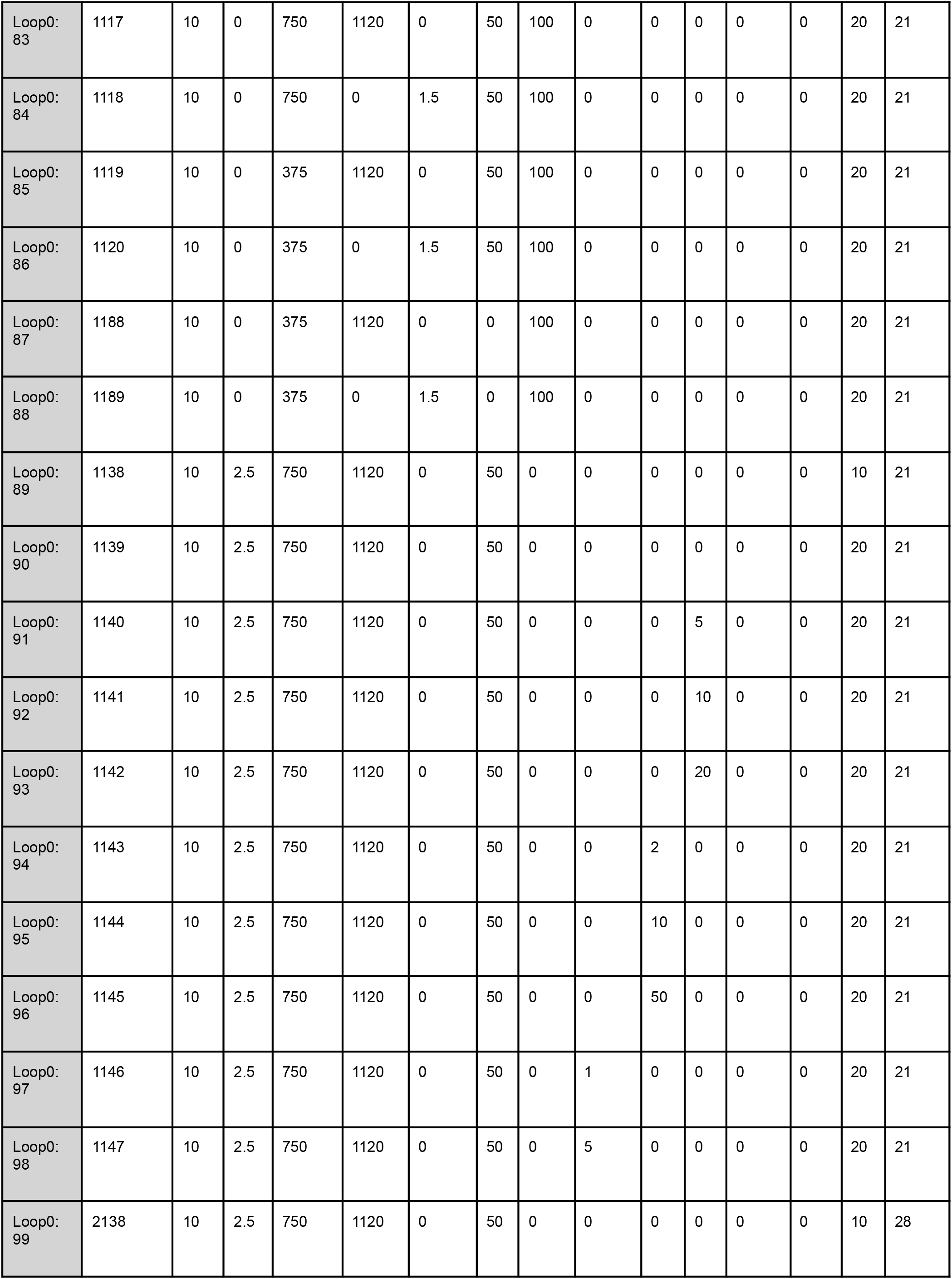
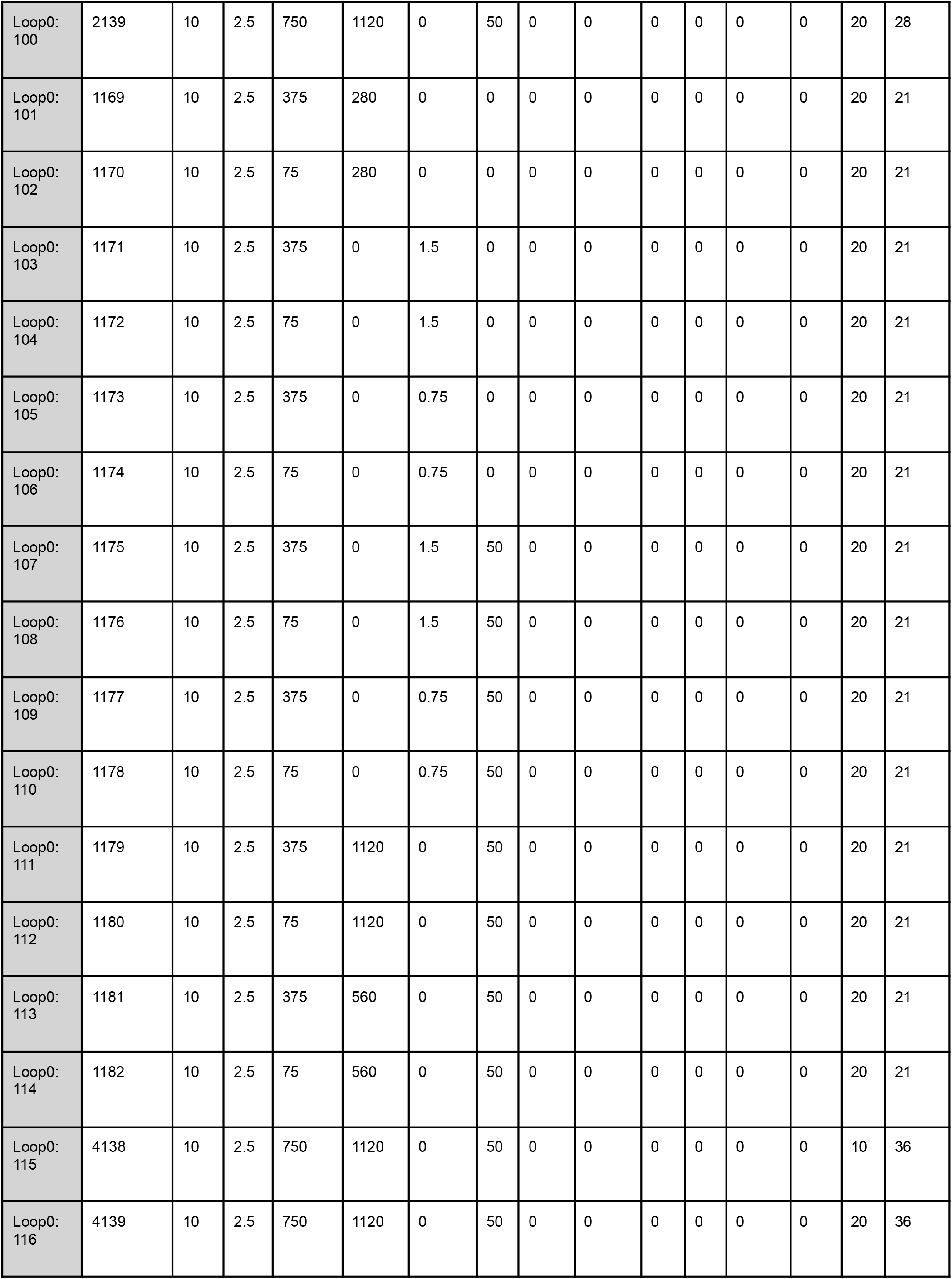
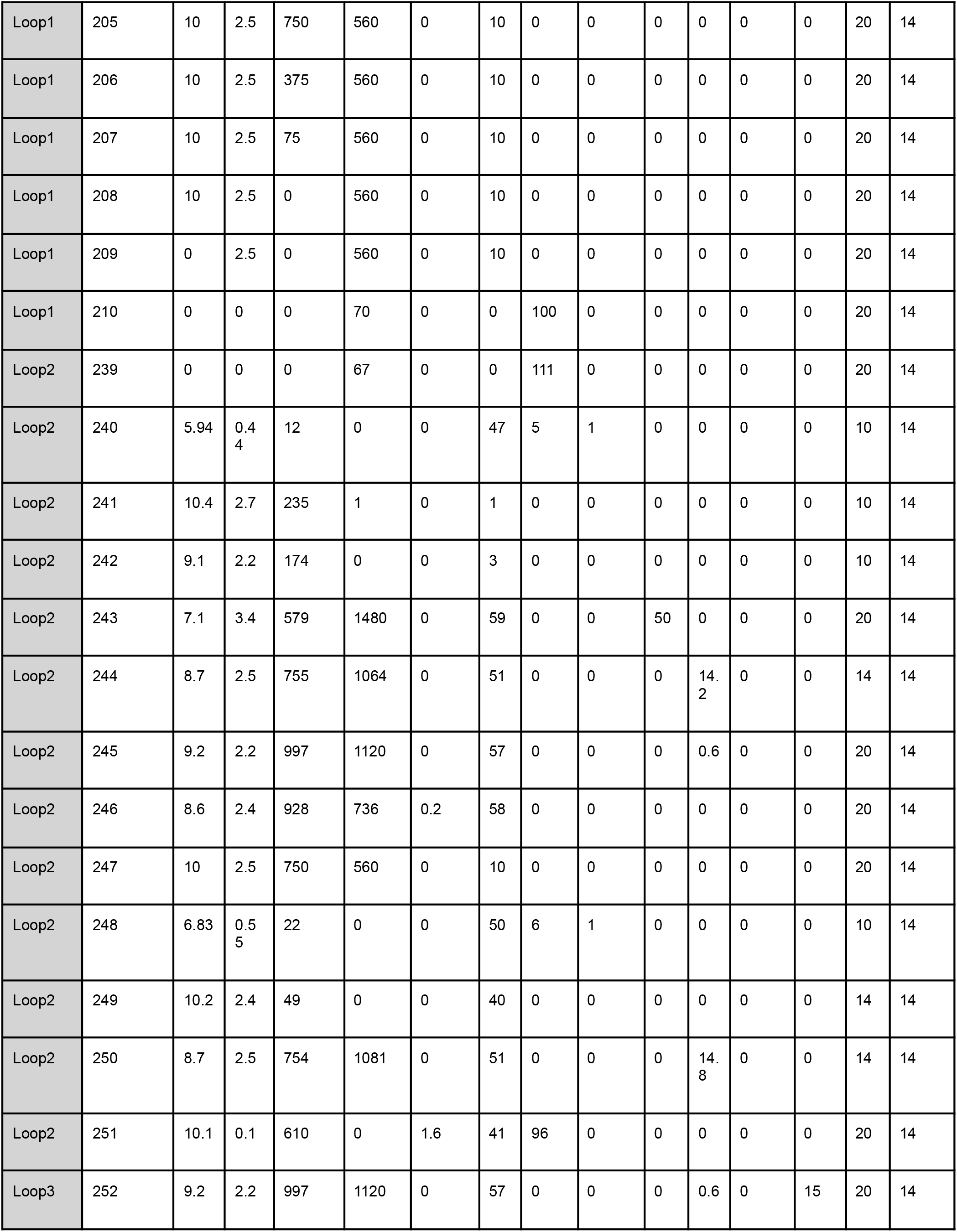
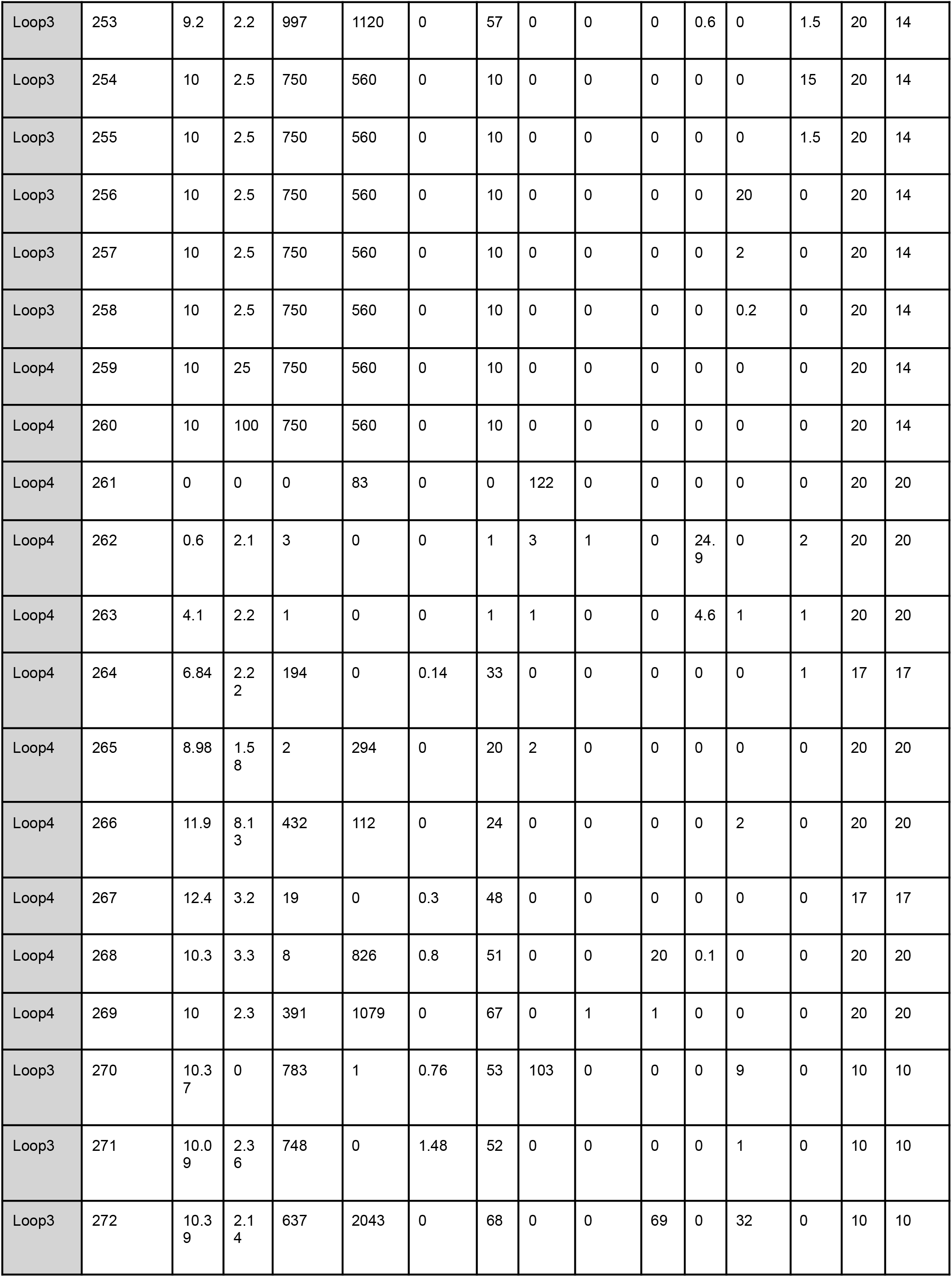
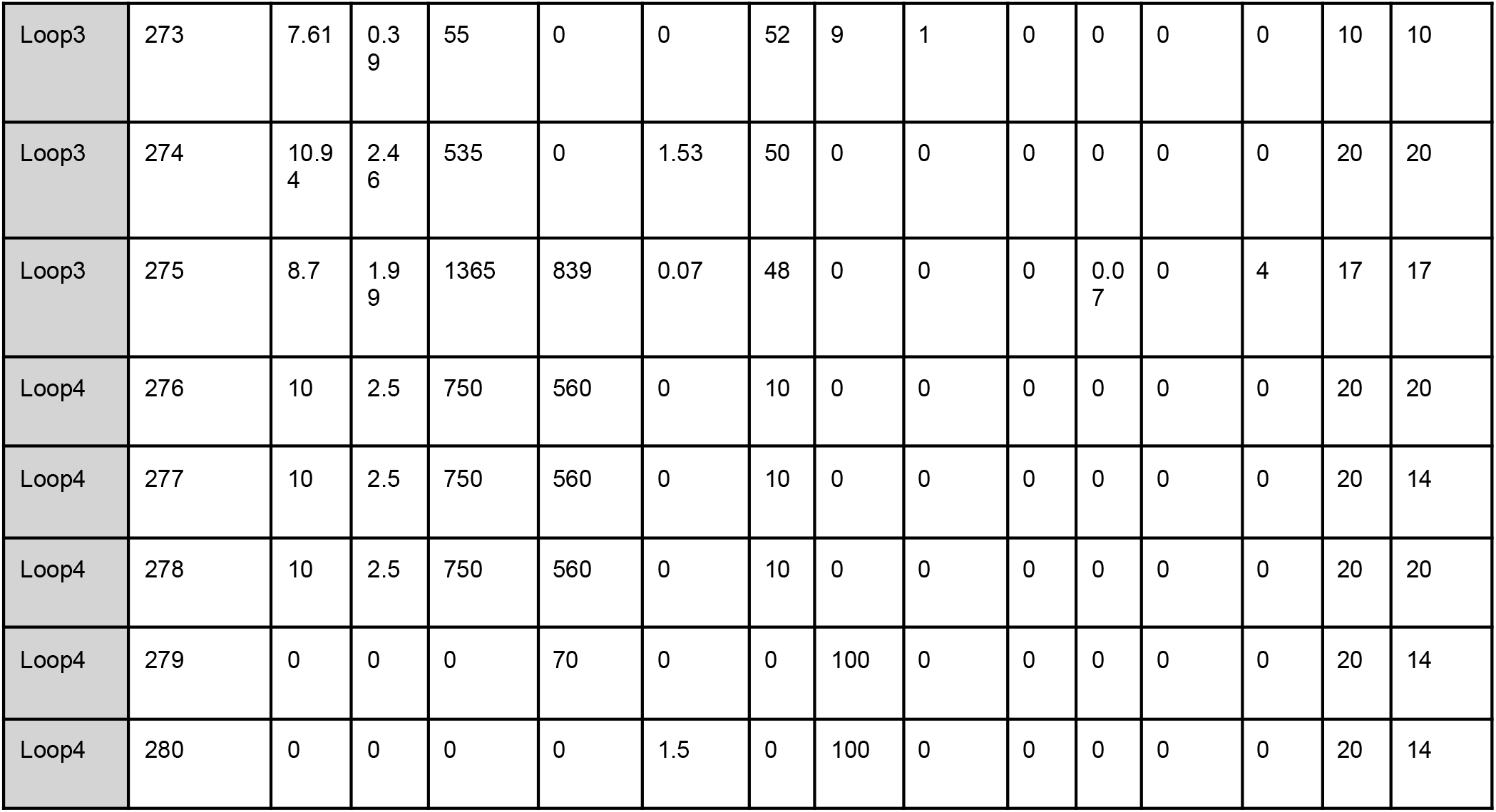
Experimental design numbers and their corresponding culture parameters used across all loops.

### Antibodies and concentrations used

**Supplementary table 7:** Antibodies in spectral flow cytometry panel for all loops.

| Markers | Fluorochrome | Dilution | Manufacturer | Cat. # | Clone |
| --- | --- | --- | --- | --- | --- |
| CD49c | BUV395 | 1 in 250 | BD Biosciences | 755279 | P1B5 |
| CD41a | BUV563 | 1 in 250 | BD Biosciences | 741371 | HIP8 |
| CD117 | BUV615 | 1 in 250 | BD Biosciences | 751616 | 104D2 |
| CD38 | BUV661 | 1 in 250 | BD Biosciences | 612970 | HIT2 |
| HLADR | BUV737 | 1 in 250 | BD Biosciences | 568351 | G46-6 |
| EPCR | BV421 | 1 in 250 | BD Biosciences | 743552 | RCR-252 |
| CD135 | BV480 | 1 in 250 | BD Biosciences | 746703 | 4G8 |
| CD56 | BV605 | 1 in 250 | Biolegend | 318334 | HCD56 |
| CD15 | BV650 | 1 in 250 | BD Biosciences | 564232 | HI98 |
| CD123 | BV711 | 1 in 250 | BD Biosciences | 563161 | 9F5 |
| CD45 | BV785 | 1 in 250 | Biolegend | 304048 | HI30 |
| CD16 | Spark Blue 550 | 1 in 250 | Biolegend | 302084 | 3G8 |
| CD7 | RealBlue 780 | 1 in 250 | BD Biosciences | 569215 | 4H9 |
| CD90 | PE | 1 in 250 | Biolegend | 328110 | 5E10 |
| CD10 | PE/Dazzle 594 | 1 in 250 | Biolegend | 982216 | HI10a |
| CD14 | PE/Fire 700 | 1 in 250 | Biolegend | 301872 | M5E2 |
| CD45RA | PE/Cy7 | 1 in 250 | Biolegend | 304126 | HI-100 |
| CD34 | APC | 1 in 250 | BD Biosciences | 555824 | 581 |
| CD235a | Alexa Fluor 700 | 1 in 250 | Thermo Scientific | 56-9987-42 | HIR2 (GA-R2) |
| CD19 | APC/Cy7 | 1 in 250 | BD Biosciences | 557791 | SJ25C1 |

## Notes

### Competing Interest Statement

Fabian J. Theis consults for Immunai, CytoReason, Valinor Industries, Bioturing and Phylo Inc., and has ownership interest in RN.AI Therapeutics, Dermagnostix, and Cellarity. The remaining authors declare no competing interests.

